# *TSC22D*, *WNK* and *NRBP* gene families exhibit functional buffering and evolved with Metazoa for macromolecular crowd sensing

**DOI:** 10.1101/2024.02.13.579840

**Authors:** Yu-Xi Xiao, Seon Yong Lee, Magali Aguilera-Uribe, Reuben Samson, Aaron Au, Yukti Khanna, Zetao Liu, Ran Cheng, Kamaldeep Aulakh, Jiarun Wei, Adrian Granda Farias, Taylor Reilly, Andrea Habsid, Kevin R. Brown, Katherine Chan, Patricia Mero, Jie Qi Huang, Maximilian Billmann, Mahfuzur Rahman, Chad Myers, Brenda J. Andrews, Ji-Young Youn, Christopher M. Yip, Daniela Rotin, W. Brent Derry, Julie D. Forman-Kay, Alan M. Moses, Iva Pritišanac, Anne-Claude Gingras, Jason Moffat

## Abstract

The ability to sense and respond to osmotic fluctuations is critical for the maintenance of cellular integrity. Myriad redundancies have evolved across all facets of osmosensing in metazoans, including among water and ion transporters, regulators of cellular morphology, and macromolecular crowding sensors, hampering efforts to gain a clear understanding of how cells respond to rapid water loss. In this study, we harness the power of gene co-essentiality analysis and genome-scale CRISPR-Cas9 screening to identify an unappreciated relationship between *TSC22D2*, *WNK1* and *NRBP1* in regulating cell volume homeostasis. Each of these genes have paralogs and are functionally buffered for macromolecular crowd sensing and cell volume control. Within seconds of hyperosmotic stress, TSC22D, WNK and NRBP family members physically associate into cytoplasmic biocondensates, a process that is dependent on intrinsically disordered regions (IDRs). A close examination of these protein families across metazoans reveals that *TSC22D* genes evolved alongside a domain in NRBPs that specifically binds to TSC22D proteins, which we have termed NbrT (<u>N</u>RBP <u>b</u>inding region with <u>T</u>SC22D), and this co-evolution is concomitant with rapid IDR length expansion in WNK family kinases. Our study identifies functions for unrecognized components of the cell volume sensing machinery and reveals that *TSC22D*, *WNK* and *NRBP* genes evolved as cytoplasmic crowding sensors in metazoans to co-regulate rapid cell volume changes in response to osmolarity.

## INTRODUCTION

Precise cell volume measurements in response to osmolarity were first reported nearly a century ago by Ponder and Saslow while measuring red cell volume^1^, and failure to adapt to osmotic challenges is lethal. To protect cells from osmotic stress, functional redundancies have evolved to regulate cell volume (e.g. genes encoding aquaporins and cation chloride co-transporters) and maintain normal cell function in response to osmolarity or the effective osmotic pressure gradient^2–5^. Cell volume changes are generally thought to occur on short time scales (seconds to minutes). During hyperosmotic stress, cells undergo crenation (shriveling) within seconds and then adapt to volume contraction by orchestrating a system of ion transporters, channels and pumps that drive net solute and water influx, resulting in regulatory volume increase (RVI)^6^. All multicellular organisms, unicellular protists and algae encode WNK kinases^7^, while unicellular organisms such as yeast are not thought to have WNKs, although homologies can be found by virtue of the WNK kinase domain^8^. Human WNK1 has recently been shown to form membrane-less phase-separated biomolecular condensates within seconds in response to hyperosmotic stress^9^. These WNK1 biocondensates are thought to be functionally relevant, occurring at native protein concentrations and at levels of hyperosmotic stress that are within the physiological range experienced by different cell and tissue types. The emergent model is that WNK kinases act as crowding sensors (i.e., through IDRs) and signal transducers (i.e. through intrinsic kinase activity) to activate cell volume recovery through the downstream effector kinases OXSR1 and SPAK^9^.

The idea of crowding-induced phase separation caused by hyperosmotic stress as a means of cell volume control was proposed decades ago^10–12^. Aside from a requirement for the C-terminal IDR of WNK1 for formation of WNK1 biocondensates^9^, our current understanding of macromolecular crowd sensing in response to hyperosmotic stress *in vivo* and *in vitro* is poorly understood. For example, the constituents and function of WNK biocondensates remain to be characterized. The WNK kinase signaling cascade is clearly significant for normal cell function as genes involved in this pathway have been associated with numerous diseases including Pseudohypoaldosterism type II and Gordon syndrome^13–15^. We postulate that macromolecular crowd sensing must have evolved distinctly between bacteria, fungi, plants and metazoans, where animal cells refined the cytoplasmic cellular milieu to facilitate biological processes necessary for different cell types and habitats to facilitate organism movement. Surprisingly little is known about molecular-crowd sensing mechanisms in different organisms, tissues, cell types and cellular compartments, potentially due to extensive redundancy and functional buffering.

Mechanisms of functional buffering can be revealed by profiling genetic interactions (GIs), which are defined as the phenotypic consequences of two or more gene perturbations that combine to produce a functional outcome. The most extreme negative GI (i.e. synthetic lethality) has been exploited as a cancer therapeutic strategy^16,17^. While profiling GIs can also reveal the function of paralog pairs, multi-gene families, where functional buffering occurs between more than two paralogs, pose an additional challenge for genome-scale functional annotation efforts^18–20^. One approach to map gene functions is to deconvolve gene-by-gene (GxG) co-dependencies through systematic efforts to generate GI profiles that can be used build interaction networks ^21^. Another approach is to define gene dependency profiles, such as the Cancer Dependency Map (DepMap) which has profiled over one thousand human cancer cell lines with genome-wide CRISPR knockout libraries^22–26^. By systematically surveying DepMap data, we identified one gene cluster composed of three essential genes (*TSC22D2, WNK1*, and *NRBP1*), each within a multi-gene family, that share similar dependency profiles. *WNK1* has long been known to play a key role in cell volume regulation^27^; however, the function of *NRBP1* is poorly characterized and the function of *TSC22D2* is even less well understood. We combine genome-wide genetic screens, live cell microscopy, proximity mapping, evolutionary analyses, drug screens and epistasis to study the WNK-related cytoplasmic macromolecular crowd sensing mechanism and reveal a complex and highly buffered regulatory network involving several multi-gene families that evolved in metazoan cells for co-regulation of cell volume homeostasis.

## RESULTS

### TSC22D2, WNK1 and NRBP1 exhibit functional dominance for cell fitness

Our initial motivation was to comprehend the scope or degree of functional redundancy amongst paralogs in the human genome using data from genome-wide pooled CRISPR fitness screens. To do this, we intersected gene families arising through whole genome duplication (i.e. ohnologs, 4935 genes from 1827 gene families^28^) with three different fitness gene sets defined by thresholding the DepMap data (i.e. DepMap30, DepMap60 and DepMap Common Essentials; see **Data S1**)^22–26^. Amongst ohnologs, we found multi-gene families that include a single fitness gene occur ∼3-fold lower than expected by chance (**Fig. S1A**). The low representation of fitness genes in ohnolog families is consistent with the idea of functional buffering amongst paralogs. However, 7%-11% (**Fig. S1A**) of ohnolog families still retain a single fitness gene, which we interpret as the functionally dominant member of a multi-gene family. Despite potential buffering from paralogs, disruption of a functionally dominant family member is sufficient to produce a measurable fitness defect for a fraction of ohnologs. To further investigate this small fraction of 197 functionally dominant ohnologs, we examined genetic co-dependencies using pairwise correlation analyses of DepMap dependency profiles (19,306 pairs of profiles). Using the DepMap30 list of fitness genes, several highly correlated gene clusters were apparent; for example, those involved in focal adhesion, thiamine pyrophosphate binding, and ARF-COPI pathway function (**Fig. 1A**). Notably, *TSC22D2*, *WNK1*, and *NRBP1* gene dependency profiles were highly correlated in large-scale CRISPR and RNAi datasets, indicating that these three genes act in related functions across a variety of genetic contexts (**Fig. 1A,B**). *TSC22D2, WNK1* and *NRBP1* are fitness genes, are members of multi-gene families (i.e., *WNK1-4*, *TSC22D1-4*, and *NRBP1-2*), and represent functionally dominant members of their gene families for cell fitness. Functional dominance is specific to a given phenotype or trait; for example, *WNK1* is the functionally dominant family member for cell line fitness (**Fig. S1B**). Functionally dominant family members are also the highest expressed gene in >80% of cases (**Fig. S1A**). However, unlike *WNK1* and *NRBP1*, which are the highest expressed genes in their gene families across most cell types and tissues examined, *TSC22D2* is expressed at much lower levels compared with its family members (**Fig. S1C**).

**Figure 1.**
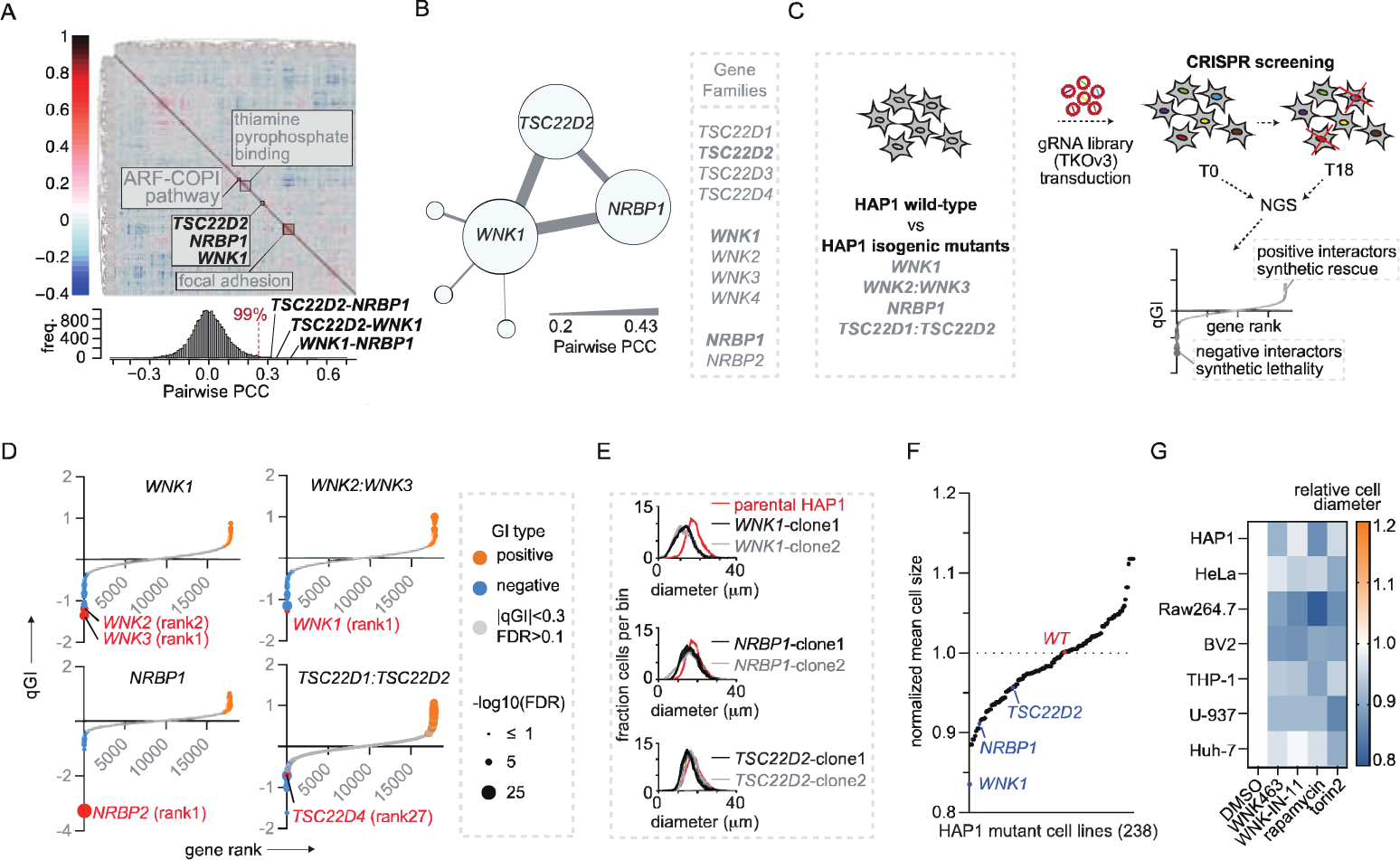
*TSC22D2, WNK1*, and *NRBP1* are co-essential, genetically buffered by family members, and regulate cell volume. **A**. The top panel shows a heatmap of hierarchically clustered pairwise Pearson gene dependency profile correlations (PCCs) derived from 197 functionally dominant ohnologs using |PCC|>0.2. Positive PCC scores represent pairwise gene dependency profiles that are positively correlated (red) and negative PCC scores represent pairwise gene dependency profiles that are negatively correlation (blue). Prominent clusters of positively correlated profiles are highlighted. The bottom panel shows the distribution of all pairwise PCCs of all 197 functionally dominant ohonologs described in Data S1. The 99% percentile of the distribution is annotated as the red dashed line. **B**. Network diagram for *TSC22D2, WNK1,* and *NRBP1* showing interactions with PCC cutoff of greater than 0.2. **C**. Experimental setup for genome-wide pooled CRISPR genetic interaction screens in HAP1 cells using *WNK1* and *NRBP1* single mutant queries and *WNK2:WNK3* and *TSC22D1:TSC22D2* double mutant queries. **D**. Rank plots of qGI scores for *WNK1* (n=2) and *NRBP1* (n=1) single mutant HAP1 query screens (left panels), as well as *WNK2:WNK3* (n=2) and *TSC22D1:TSC22D2* (n=2) double mutant HAP1 query screens (right panels). Legend-defining dot size and dot color are shown on the right. **E**. Cell diameter distributions for two unique single mutant *TSC22D2*, *WNK1*, or *NRBP1* HAP1 clones, shown in grey or black. Parental HAP1 distributions are shown in red for comparison. **F**. Rank order plot of normalized mean cell volume mearurements for 238 isogenic mutant HAP1 query cell lines. **G**. Heatmap of normalized mean cell volume changes across the indicated cell lines following a 24hrs treatment with either WNK463, WNK-IN-11, Rapamycin or Torin 2 at the indicated doses. See also Fig. S1 and Data S1.

To further explore the idea of functional dominance in cultured isogenic cells, we generated stable HAP1 clonal cell lines mutant for *TSC22D2*, *WNK1*, or *NRBP1* (**Fig. 1C**), where the corresponding protein levels were substantially decreased or undetectable **(Fig. S1D)**, and subjected these mutant query cell lines to genome-wide pooled loss-of-function CRISPR fitness screens (**Fig. 1C**) to measure negative and positive genetic interactions (GIs)^21,29^. If gene family members are capable of genetic buffering, the expectation is that family members of the query gene would be negative GIs. In the *TSC22D2* query screen, we observed a negative GI with *TSC22D1* (FDR=8.8E-2), but the reciprocal interaction was not observed in the *TSC22D1* query screen (**Fig. S1E**). We also observed negative GIs in the *WNK1* query screen with both *WNK2* (**Fig. 1D**, FDR=6.5E-10) and *WNK3* (**Fig. 1D**, FDR=7.5E-11), as well as a negative GI in the *NRBP1* query screen with *NRBP2* (**Fig. 1D**; FDR=4.1E-25). *WNK4* is not detectably expressed in HAP1 cells and was not expected to buffer the mutant *WNK1* cells. Also, no GI with *WNK1* was observed in the *WNK2* and *WNK3* single query screens (**Fig. S1E**). Lastly, no GIs were observed between the *TSC22D2*, *WNK1*, and *NRBP1* gene families, despite the reproducible nature of these screens based on within-and-between correlation measurements (**Fig. S1F**)^30^. Together, these results provide evidence that genetic buffering can occur within the *TSC22D*, *WNK* and *NRBP* gene families when systematically surveying digenic interactions and when the functionally dominant family member serves as the query gene.

To enhance our understanding of genetic buffering amongst these gene families, we also constructed isogenic *TSC22D1:TSC22D2* and *WNK2:WNK3* double-mutant HAP1 cells and performed genome-wide pooled loss-of-function CRISPR screens in each of these double-mutant query cell lines. In this context, we found negative trigenic interactions amongst *TSC22D1*, *TSC22D2*, and *TSC22D4* (**Fig. 1D**, FDR=1.8E-4), as well as *WNK1*, *WNK2*, and *WNK3* (**Fig. 1D**, FDR=1.7E-4). Although not comprehensive for all family members, these observed higher-order trigenic interactions provide further evidence of genetic buffering within each of the *TSC22D* and *WNK* gene families in human HAP1 cells.

### TSC22D2, WNK1 and NRBP1 regulate cell volume

Because *WNK1* has been implicated in cell volume regulation ^9^, we next examined whether stable perturbation of *TSC22D2*, *WNK1*, or *NRBP1* impacts cell volume in HAP1 cells. Remarkably, the average cell volume of *TSC22D2*, *WNK1* and *NRBP1* mutant clonal HAP1 cells were smaller than parental HAP1 cells (**Fig. 1E**). We also measured the cell volume of 238 unique HAP1 isogenic query mutant cell lines representing 146 unique genes whose functions spanned biological processes (**Data S1**) and found that *TSC22D2*, *WNK1* and *NRBP1* mutants had cell volumes that were consistently smaller than the average HAP1 mutant cell line (**Fig. 1F**). To corroborate these results, HAP1 cells were treated with the pan-WNK kinase inhibitor WNK463^31,32^ and a dose-dependent decrease in cell volume was observed, comparable to the MTOR inhibitor Torin2, which inhibits both mTORC1 and mTORC2 complexes and is known to cause smaller cell size broadly across eukaryotic models (**Fig. 1G** and **S1G**)^33^. More modest cell volume changes were also observed in HAP1 cells treated with WNK-IN-11, a WNK1 inhibitor, as well as the mTORC1 inhibitor rapamycin ^34^. The decrease in cell volume following WNK463 treatment was consistent in multiple human and mouse cell lines including RAW264.7, BV2, THP-1, HeLa-Kyoto, U-937, and Huh-7 cells (**Fig. 1G** and **S1H**). Thus, both stable genetic and acute chemical perturbation of the *TSC22D2-WNK1-NRBP1* co-essential gene cluster led to cell volume decreases in mammalian cells.

### Endogenous TSC22D2, WNK1 and NRBP1 colocalizes to reversible biocondensates under hyperosmotic stress

To our knowledge, the *TSC22D2* and *NRBP1* genes have not previously been linked to cell volume regulation; however, WNK1 forms biocondensates and regulates cell volume increase by acting as a macromolecular crowding sensor ^9^. To investigate whether the products of *TSC22D2* and/or *NRBP1* are involved in crowd sensing, we first endogenously tagged the canonical isoform of TSC22D2 in HAP1 cells with the fluorescent protein mScarlet at its C-terminus (i.e., *TSC22D2^mScarlet^*) (**Fig. 2A**), confirmed that the endo-tagged protein was properly expressed (**Fig. S2A**), and performed live-cell microscopy under hyperosmotic stress to pinpoint TSC22D2 localization. Treating *TSC22D2^mScarlet^*cells with increasing amounts of sodium chloride (NaCl), potassium chloride (KCl), or sorbitol resulted in the rapid formation of >10 bright cytoplasmic foci per cell that were on average ∼200 nm or less in diameter (**Fig. 2B** and **S2B**). Increasing hyperosmotic stress caused more and denser cytoplasmic foci; the highest stress condition led to the dissolution of the condensates or redistribution of the protein to other cellular sites (**Fig. S2C**)^35^. At medium levels of stress, these foci persisted and then gradually dissolved following prolonged hyperosmotic stress (**Fig. 2B** and **S2C**). TSC22D2 foci were only observed under salt or sorbitol stress, and not using small molecule crowding agents such as urea (**Fig. S2B**), consistent with previous observations with WNK1 foci ^9^.

**Figure 2.**
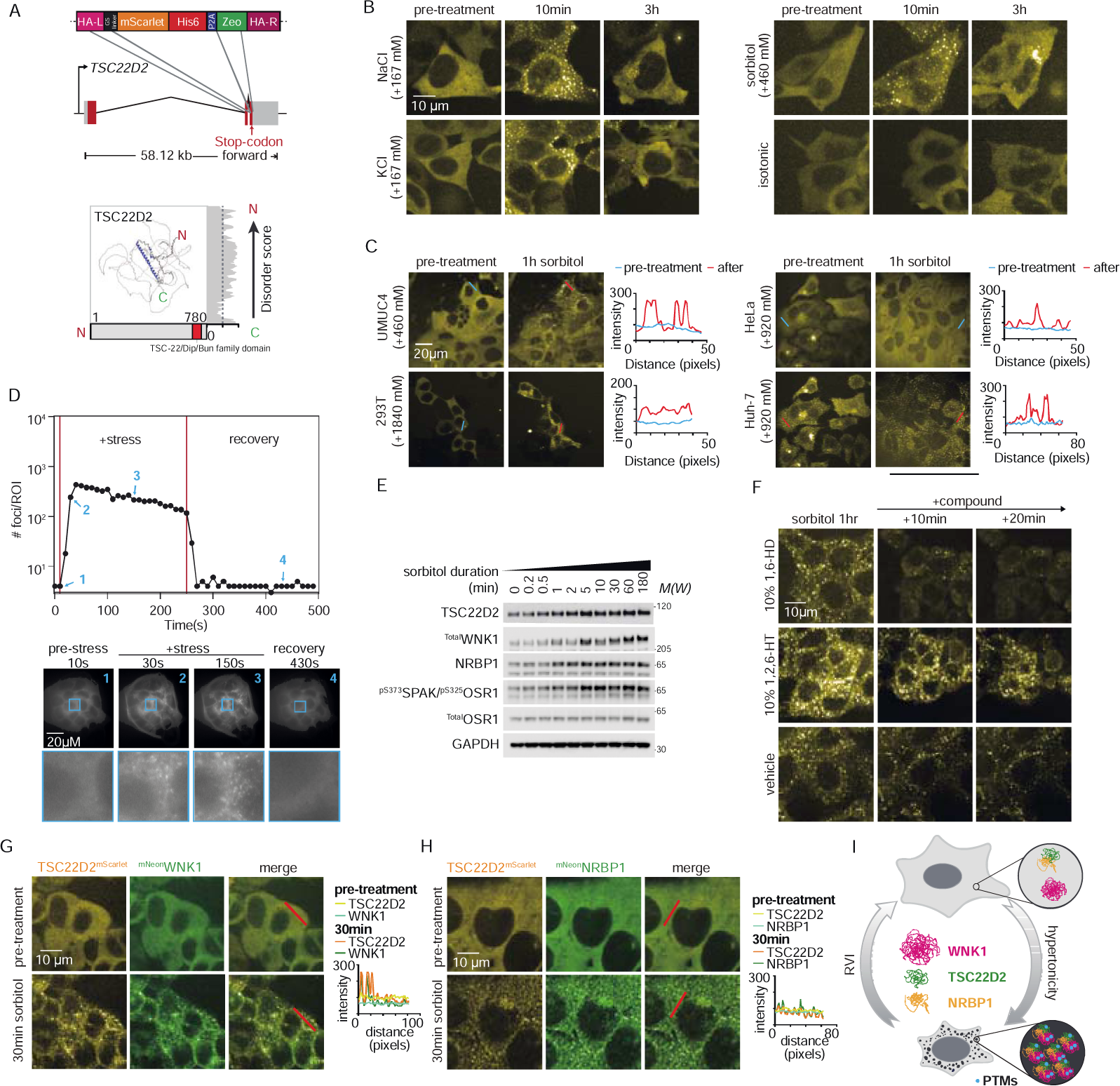
Endogenous TSC22D2, WNK1 and NRBP1 form cytoplasmic biomolecular condensates and co-localize following hyperosmotic stress. **A**. The top panel describes strategy for tagging the C-terminus of TSC22D2 with mScarlet at the endogenous locus; the bottom panel shows predicted canonical TSC22D2 structure. Sidebar of the TSC22D2 structure shows disorder score predicted by IUPRED^58^. **B**. Time-lapse fluorescence microscopy of HAP1 *TSC22D2^mScarlet^* cells with the indicated hyperosmotic stress (i.e. NaCl, KCl, or sorbitol). **C**. Fluorescence microscopy of *TSC22D2^mScarlet^* endo-tagged cell lines including HeLa-Kyoto, Huh-7, HEK293T and UM-UC-4 before and after treatment with the indicated concentrations of sorbitol. Line graphs to the right represent image intensity values as a function of the line indicated in the pre-treatment (green) or 1h sorbitol (red) images. Lines for intensity variation measurements are located at the same position in the pre- and post-treatment images and processed with ImageJ. **D**. Quantification of TSC22D2 foci formation in the TIRF movies of HAP1 *TSC22D2^mScarlet^*cells following sorbitol stress and recovery. Representative frames pre-treatment, post-treatment and post-recovery are indicated by the four arrows and numbers in the graph and shown at the bottom. The blue box indicated in the example frames is expanded to show the TSC22D2 foci from stress. Movies files are included as Supplementary Movie files. **E**. Immunoblots for TSC22D2, WNK1, NRBP1, OSR1, and phosphorylated OSR1^S325^/SPAK^S373^. GAPDH serves as the loading control. **F**. Microscope images of a time course of TSC22D2 foci after sorbitol treatment followed by treatment of cells with either 1,6-Hexanediol (1,6-HD) or 1,2,6-Hexanetriol (1,2,6-HT). See also Fig. S2 and Movies S1-3. Representative microscope images of HAP1 **G.** *^mNeon^WNK1:TSC22D2^mScarlet^* or **H.** *^mNeon^NRBP1:TSC22D2^mScarlet^* cells before and after 30 min of hyperosmotic stress with 460mM sorbitol. Line graphs to the right represent intensity values as a function of the red line indicated in the before and after merged images. Line colors are defined in the inset. **I**. Model for TSC22D2, WNK1 and NRBP1 contribution to regulatory cell volume control. See also Fig. S2 and S3.

To confirm our observations were not HAP1 dependent, *TSC22D2* was endo-tagged with mScarlet at the C-terminus in four additional human cell lines originating from different tissues including Huh-7 (liver), HeLa (cervix), HEK293T (kidney), and UM-UC-4 (bladder). Using live cell imaging, TSC22D2 foci were evident following hyperosmotic stress in all four cell lines, comparable to what was observed in HAP1 cells (**Fig. 2C**).

TSC22D2 co-localization studies were next performed with sub-cellular markers to identify whether TSC22D2^mScarlet^ cytoplasmic foci co-localize with known structures including CLTC for clathrin-mediated endocytic vesicles, LAMP1 for lysosomes, SQSTM1 for aggresomes, DCP1A for processing bodies, G3BP1 for stress granules, and TOMM20 for mitochondria (**Fig. S2D**). These results indicate that TSC22D2 cytoplasmic foci induced by hyperosmolarity partially co-localize with clathrin but are distinct compared to other known cytoplasmic organelles.

The kinetics of TSC22D2 foci formation following hyperosmotic stress was also examined by total internal reflection fluorescence microscopy and foci were observed within 10 seconds and saturated within 30 seconds after hyperosmotic stress, then disappeared within 10 seconds after removing the stress (**Fig. 2D**; **Movies S1-S3**). These changes were accompanied by a corresponding cell volume decrease and increase, consistent with the effects of hyperosmolarity and volume recovery, respectively (**Fig. 2D**; **Movies S1-S3**). Intriguingly, time-dependent post-translational modifications on both TSC22D2 and WNK1 proteins happen after the formation of TSC22D2 foci. Phosphorylation of SPAK/OSR1 and WNK1, as well as band up-shifts of both TSC22D2 and WNK1, are observed after ∼5 minutes of hyperosmotic stress (**Fig. 2E**). Thus, TSC22D2 rapidly forms transient and reversible cytoplasmic foci within seconds in response to hyperosmotic stress.

Despite some of the conceptual issues employing 1,6-hexanediol (1,6-HD)^36^, this compound has been used to dissolve weak hydrophobic interactions and some biomolecular condensates in live cells ^37^. To examine whether TSC22D2 cytoplasmic foci are phase-separated biomolecular condensates, *TSC22D2^mScarlet^*HAP1 cells were hyperosmotically stressed with sorbitol, and the persistence of foci was monitored following addition of 1,6-HD, or 1,2,6-hexanetriol (1,2,6-HT) as a viscosity control. Dissolution of TSC22D2 foci was clearly observed following 1,6-HD but not 1,2,6-HT treatment, indicating that TSC22D2 foci are consistent with being liquid-liquid phase-separated biomolecular condensates (**Fig. 2F**).

Both TSC22D2 and WNK1 are extensively disordered proteins (**Fig. 2A** and **S3A**) and are subject to post-translational modifications in minutes following sorbitol stress (**Fig. 2E**)^9,27^. To assay whether TSC22D2 co-localizes with WNK1 under hyperosmotic stress, we generated double endo-tagged *^mNeon^WNK1:TSC22D2^mScarlet^*HAP1 cells and observed strong co-localization patterns between TSC22D2 and WNK1 foci in *^mNeon^WNK1:TSC22D2^mScarlet^* HAP1 cells following sorbitol treatment (**Fig. 2G and S3B**). In contrast to TSC22D2 and WNK1, NRBP1 is composed of a short, disordered N terminus (1-54), a highly ordered kinase-like domain (68-327) and an ordered C terminal domain (439-535) (**Fig. S3A**). Nevertheless, we also observed strong patterns of co-localization between TSC22D2 and NRBP1 foci in *^mNeon^NRBP1:TSC22D2^mScarlet^* HAP1cells following hyperosmotic stress (**Fig. 2H** and **S3B**). Thus, TSC22D2 and NRBP1 proteins co-localize to cytoplasmic foci with WNK1 under hyperosmotic stress, indicating that TSC22D2 and NRBP1 are likely constituents of macromolecular crowd sensing ‘WNK bodies’ (**Fig. 2I**).

### *TSC22D2* mediates the translocation of *NRBP1* into biomolecular condensates through conserved binding domains

Both TSC22D2 and WNK1 are largely composed of peptide sequence containing intrinsically disordered regions (IDRs), in contrast to the NRBP1 pseudo-kinase (**Fig. 2A** and **S3A)**. So how does NRBP1 form biocondensates under hyperosmotic stress? Previous studies and publicly available large-scale interaction data indicate that human TSC22Ds directly bind to NRBPs ^38–42^. Moreover, the fly homologs of TSC22D2 (bun) and NRBP1 (Madm) were shown to interact using the yeast two-hybrid assay ^43^. Consistent with these observations, we confirmed that endo-tagged ^mNeon^TSC22D2 interacts with endogenous NRBP1 in HAP1 cells using immunoprecipitation followed by Western blotting (**Fig. 3A**). These results suggest that, under hyperosmotic stress, WNK bodies may contain NRBP1 through its association with TSC22D2. To test this hypothesis, we generated *TSC22D1:TSC22D2:TSC22D4* triple mutant HAP1 cells harbouring endo-tagged *^mNeon^NRBP1*, in order to minimize genetic buffering effects within the *TSC22D* gene family, and to examine endogenous NRBP1 localization (**Fig. 3B**). Strikingly, ^mNeon^NRBP1 fails to partition into visible biomolecular condensates in *TSC22D1:TSC22D2:TSC22D4* triple mutant HAP1 cells, even in strong hyperosmotic conditions (**Fig. 3B** and **S3C**). Therefore, we conclude that *TSC22Ds* are required for NRBP1-mediated phase separation into WNK biocondensates.

**Figure 3.**
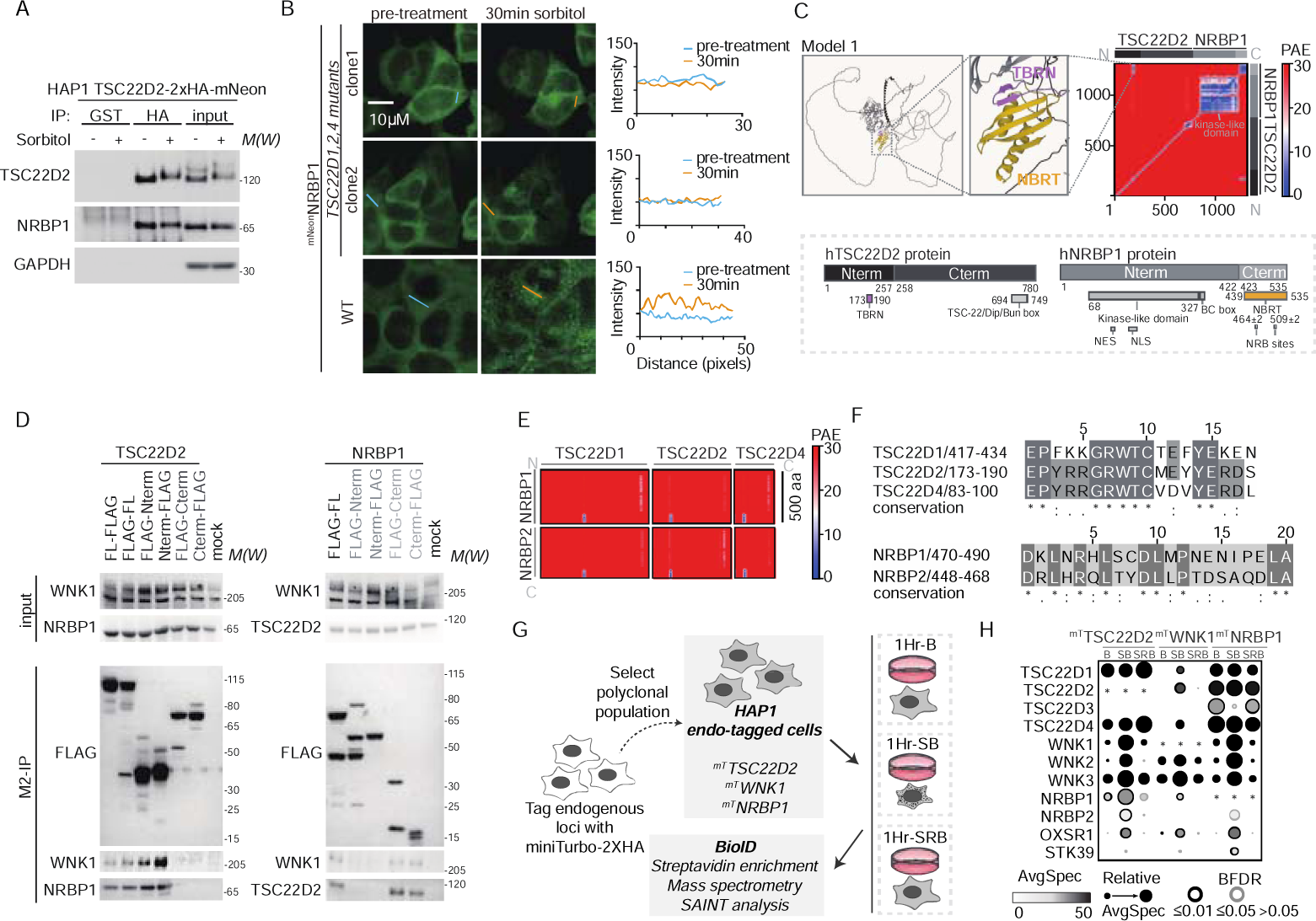
TSC22D2 mediates NRBP1 into biocondensates through conserved binding domains. **A**. Immunoprecipitation from HAP1-*TSC22D2*^2x^*^HAmNeon^*lysates with anti-HA beads followed by immunoblotting for endogenous TSC22D2, NRBP1 or the loading control GAPDH. Control immunoprecipitations used anti-GST beads. Sample loading is indicated including the input lysates. **B**. Microscope images before and after 30 minutes of hyperosmotic stress with 460 mM sorbitol in ^mNeon^NRBP1 endo-tagged HAP1 parental cells and HAP1 cells with *TSC22D1:TSC22D2:TSC22D4* triple gene mutants. The line graphs to the right represent intensity values as a function of the blue lines (pre-treatment) and orange lines (30min sorbitol) indicated in the images. **C**. The top left panel shows AF2 prediction of 3D structure of human TSC22D2 and NRBP1 interaction through TbrN and NbrT domains. The top right panel shows the PAE score matrix (0-31.75Å) of the AF2 model1 prediction for human TSC22D2 and NRBP1. Blue in the colour spectrum shows a low PAE score and red in the colour spectrum shows a high PAE score. The bottom panel shows 1D structures of human TSC22D2 and NRBP1 as well as the truncation designs used in part D. **D**. Co-immunoprecipitation with exogenous FLAG-tagged TSC22D2 or NRBP1 constructs using anti-M2 beads followed by immunoblotting for FLAG, TSC22D2, WNK1, or NRBP1. Overexpression of mClover constructs with no FLAG was used as mock control. Sample loading is indicated including the input lysates. **E**. PAE score matrix (0-31.75Å) of representative AF2 predictions for human TSC22D1, TSC22D2 and TSC22D4 interactions with human NRBP1 and NRBP2. Blue in the colour spectrum shows a low PAE score and red in the colour spectrum shows a high PAE score. **F**. Amino acid sequences of predicted TSC22D-NRBP paralog interaction interface (distance<5Å). For sequence conservation symbols, an * (asterisk) indicates positions which have a single, fully conserved residue, a : (colon) indicates conservation between groups of strongly similar properties, a . (period) indicates conservation between groups of weakly similar properties. See in-depth definition EMBL-EBI (https://www.ebi.ac.uk/seqdb/). **G**. Experimental setup for condition-specific proximity mapping of TSC22D2, WNK1 and NRBP1 associations in HAP1 cells with endogenous N-terminal miniTurbo (mT) tags. *^mT^TSC22D2*, *^mT^WNK1*, and *^mT^NRBP1* cells were treated with 1hr 50mM biotin (1Hr-B), 1hr 460 mM sorbitol containing 50mM biotin (1Hr-SB) and 1hr 460mM sorbitol then followed by recovery of isotonic medium containing 50mM biotin (1Hr-SRB) prior to mass-spectrometry. **H**. Dot-plot analysis of the proximity interactions for each of TSC22D2, WNK1, and NRBP1 baits in the no stress (1Hr-B), sorbitol stress (1Hr-SB) and recovery (1Hr-SRB) conditions (n=3). TSC22D, WNK, and NRBP paralogs are recovered as preys, as well as OXSR1 (OSR1) and STK39 (SPAK). Dots are plotted as a function of the Bayesian False Discovery Rate or BFDR and spectral counts, as indicated in the legend.

To further understand the structural basis for the TSC22D2-NRBP1 interaction, we used AlphaFold2-Multimer (AF2-M) to predict potential interaction regions (**Fig. 3C**)^44^. Notably, 24 out of the top 25 AF2-M structural models predict a strong interaction interface (average PAE=7.42 in model1) between an 18 amino acid stretch on the N-terminus of TSC22D2 (173-190), which we refer to as TbrN (<u>T</u>SC22D <u>B</u>inding <u>R</u>egion with <u>N</u>RBP), and the structured C-terminus of NRBP1, which we refer to as NbrT (<u>N</u>RBP <u>B</u>inding <u>R</u>egion with <u>T</u>SC22D)(**Fig. 3C**). One previous study made reference to NbrT as the MIf1-binding region ^45^. To validate that the TbrN and NbrT sequences mediate the physical interaction between TSC22D2 and NRBP1 in cells, we generated a series of expression constructs representing truncated proteins with C- or N-terminal FLAG epitope tags including TSC22D2-Nterm (1-257), TSC22D2-Cterm (258-780), NRBP1-Cterm (1-422), and NRBP1-Nterm (423-535) (**Fig. 3C**). These constructs were subsequently transfected into HEK293T cells and immunoprecipitations were performed with cell lysates using anti-FLAG M2 beads. Endogenous WNK1 and NRBP1 were immunoprecipitated with full-length (FL) TSC22D2 and the TSC22D2-Nterm mutant, but not with TSC22D2-Cterm mutants or the mock control (**Fig. 3D**). Furthermore, endogenous WNK1 and TSC22D2 were immunoprecipitated with FL NRBP1 and the NRBP1-Cterm mutant, but not with the NRBP1-Nterm mutant or the mock control (**Fig. 3D**). These results provide experimental evidence that the predicted TbrN within TSC22D2 likely interacts with the predicted NbrT within NRBP1, and supports the predictive power of AF2-M.

### WNK biocondensates contain TSC22D and NRBP proteins

To further investigate whether the TSC22D2 TbrN and NRBP1 NbrT extend to other TSC22D and NRBP paralogs, and possibly promote pairwise interactions between all members of these two gene families (i.e. TSC22D1, TSC22D2, TSC22D3, TSC22D4, NRBP1, and NRBP2), we rendered AF2-M predictions for all possible TSC22D-NRBP heterodimers, with the exception of TSC22D3 as it does not contain a predicted TbrN (**Fig. S6D**). TbrNs are conserved across TSC22D1, TSC22D2, and TSC22D4 and was predicted by AF2-M to bind with NbrTs on both NRBP1 and NRBP2 (**Fig. 3E and 3F**), raising the possibility that TSC22D and NRBP family members may associate promiscuously to form biocondensates upon hyperosmotic stress, providing some support for the extensive genetic buffering observed in genome-wide pooled CRISPR fitness screens (**Fig. 1D**).

To examine the physical association landscape for TSC22D2, WNK1 and NRBP1, we performed biotinylation proximity association mapping (BioID) in HAP1 cell lines that were endo-tagged with miniTurbo-2xHA (mT) and express ^mT^TSC22D2, ^mT^WNK1 or ^mT^NRBP1 as bait proteins (**Fig. 3G, S3D, S3E** and). Biotinylation, mass spectrometry and SAINT analysis were performed in each of these engineered cell lines in biological triplicate under three conditions: biotin for one hour (1Hr-B); biotin plus sorbitol for one hour (1Hr-SB); or one hour of sorbitol followed by recovery in biotin-containing iso-osmotic media for an additional hour (1Hr-SRB) (**Fig. 3G**). Streptavidin signals colocalize with bait proteins containing an HA epitope tag, indicating that BioID is capable of capturing proteins in proximity to our bait of interest (**Fig. S3E**) ^46–49^. Significant preys from each of the BioID experiments were used to generate condition-specific networks of protein associations and gene set enrichment identified bioprocesses related to stress granule formation, mitotic spindle, RNA binding, clathrin coated vesicles and cytoskeleton-related functions (**Fig. S3F**). More specifically, BioID detected significant associations between members of the TSC22D, WNK, and NRBP families as well as the WNK substrates OXSR1 (OSR1) and STK39 (SPAK), particularly following sorbitol stress (**Fig. 3H**). NRBP1 and TSC22D1,2,4 associations were observed in all conditions (**Fig. 3H**), consistent with the predicted interactions above amongst family members. Curiously, we also detected an NRBP1-TSC22D3 association that decreases under sorbitol stress, suggesting that there may be higher order interactions (e.g. hetero-dimerization amongst TSC22D proteins) that could bring TSC22D3 in proximity to NRBP1 (**Fig. 3H**). These results provide evidence that TSC22D and NRBP protein families associate with WNKs and their downstream effector kinases OXSR1 and STK39 and are constituents of WNK bodies.

To extend the BioID results, we further generated endo-tagged HAP1 cell lines for additional *TSC22D* and *WNK* genes including *^mNeon^TSC22D1:TSC22D2^mScarlet^*, *^mNeon^TSC22D4:TSC22D2^mScarlet^*, and *^mNeon^WNK3:TSC22D2^mScarlet^*cells for co-localization studies in live cells. Consistent with the BioID results, patterns of co-localization were clearly observed between TSC22D2 and TSC22D1, TSC22D2 and TSC22D4, and TSC22D2 and WNK3 following hyperosmotic stress (**Fig. S4A**). Therefore, TSC22D, WNK, and NRPB family members physically associate and co-localize. However, the composition of individual biocondensates that are formed within a cell following osmotic stress appears variable in terms of stoichiometries for each of the TSC22D, WNK, and NRBP proteins. Several prey proteins with available antibodies were also chosen for validation by co-localization. For example, endogenous CLINT1 and WNK1 are mostly colocalized with ^mScarlet^TSC22D2 foci (**Fig. S4B**). To recap, hyperosmotic stress causes the formation of biomolecular condensates that are formed within seconds and contain <u>T</u>SC22D, <u>W</u>NK, and <u>N</u>RBP (i.e., TWN) proteins, which we will refer to as ‘TWN bodies’ hereafter for simplicity. In other words, TWN bodies are biomolecular condensates that form in response to hyperosmotic stress and minimally contain some combination TSC22D, WNK, and NRBP proteins.

To examine the relationship between TWN-bodies and stress granules, we assayed G3BP1 localization by immunofluorescence in *^mNeon^WNK1:TSC22D2^mScarlet^*and *^mNeon^NRBP1:TSC22D2^mScarlet^* HAP1 cells following stress granule formation triggered by either sodium arsenate or sorbitol ^50^. Thirty minutes of sodium arsenate treatment triggered G3BP1 stress granules, but not TWN bodies. In contrast, sorbitol treatment triggered both TWN bodies and stress granules, with only partial co-localization between G3BP1 and TWN bodies (**Fig. S4C** and **S4D**). Using double endo-tagged *^mNeon^TSC22D2:G3BP1^mScarlet^* HAP1 cells coupled with live cell microscopy, G3BP1*^mScarlet^*- and *^mNeon^*TSC22D2-containing condensates were observed to have distinct formation kinetics (**Fig. S5A** and **S5B**). In fact, sorbitol treatment of *^mNeon^G3BP1:TSC22D2^mScarlet^* cells showed that TSC22D2 condensates formed within ∼10 seconds, concurrent with a rapid decrease in cell volume, whereas G3BP1 condensates formed with slower kinetics in ∼5-10 minutes, indicating that TWN bodies and cell volume decrease occur faster than activation of WNK1 kinase and formation of stress granules.

### *TWN* bodies likely evolved in metazoans

The IDR fraction of proteins that associate with TSC22Ds, WNKs, and NRBPs derived from the BioID studies described above was significantly higher than the background set of HAP1-expressed genes (**Fig. 4A**). Thus, the IDRs within the TSC22D family of proteins may help to facilitate multivalent interactions promoting condensate formation, as has been shown already for the WNK1 C-terminal IDR ^9^, and reported previously for other types of phase-separated condensates, such as stress granules and P-bodies ^50,51^.

**Figure 4.**
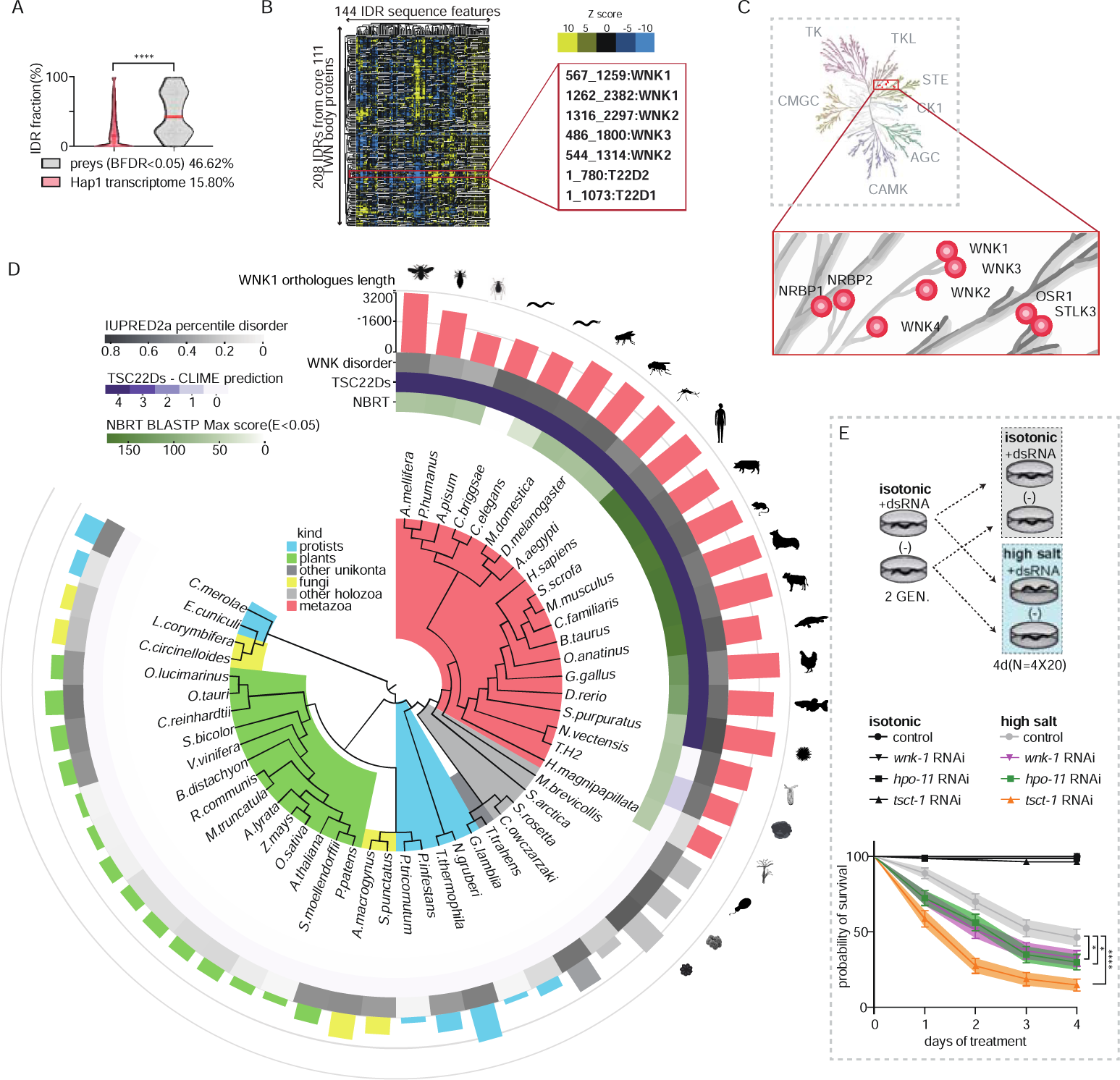
Emergence of TSC22D family of proteins in metazoans. **A**. Percentile IDR fraction of the HAP1 transcribed proteome (mean=15.80%) and a union of BioID preys (mean=46.62%) with BFDR<0.05 from all baits and conditions predicted by SPOT-DISORDER. Wilcoxon test is used for calculation of p value (****p<0.0001). **B**. The left panel shows a global heatmap consisting of a 144×208 z-score matrix of evolutionary signatures with the core set proximity associations with TSC22D2, against total human proteome. The z-score legends are selected against (blue) or for (yellow) amongst the indicated IDRs. The right panel shows a selected portions of heatmap presented in **Fig. S6C** highlighting key evolutionary features that are selected against (blue) or for (yellow) amongst the indicated disordered subsequences containing N terminal WNK1-3 and TSC22D1-2 (cluster1, corresponding to disorder S1 in Fig. S6D). **C**. Schematic of human kinome tree with major families indicated. The expanded box highlights the region in the STE20 family that contains WNK and NRBP kinases. **D.** Circular phylogenetic plot highlighting species from all major kingdoms and including intrinsic disordered predictions for WNK gene products using IUPRED2a, predicted polypeptide lengths for WNK gene products, CLIME predictions for presence/absence of *TSC22D* genes, and prediction of presence of NbrT domain on NRBP homologues by BLASTP matrix (E<0.05). **E**. Survival curves showing the effects of RNAi knock-down of *tsct-1*, *wnk-1* and *hpo-11* in *C. elegans* N2 worms under isotonic or high salt conditions. Kaplan-Meier survival estimate is used for p-values where *p<0.05 and ****p<0.0001. See also Fig. S4, S5, Tables S1, S2 and Data S2.

A recent proteome-wide analysis of human IDRs (i.e., human IDR-ome), focusing on 144 features for each IDR (**Data S2**), demonstrated that evolutionary patterns of bulk molecular features of the human IDR-ome enable classification and functional annotation ranging from sub-cellular localization and membraneless organelles to the constitution of the cytoskeleton and transmembrane signalling ^52,53^. Using these same features, clustering analyses of IDR sequences from TWN body-associated proteins demonstrated that TSC22D proteins share striking features with WNK C-terminal regions (disordered S1), such as strong depletion in charged residues and enrichment of hydrophobic and aliphatic residues, consistent with prion-like low complexity regions (**Fig. 4B**, **S6A-S6D**). Moreover, the N-terminal IDRs of WNK proteins (disordered S2) showed an increased propensity for glutamic acid residues and shared similarities to NRBP1 (**Fig. 4D, S6A-S6D**).

WNK and NRBP kinases are part of the STE20 family of kinases (**Fig. 4C**)^54^. To further address the evolutionary ancestries of the *TSC22D*, *WNK*, and *NRBP* genes, we turned to clustering by inferred models of evolution (CLIME)^55,56^. In contrast to previous reports^8^, CLIME predicts the presence of WNK1 and NRBP1 proteins across all kingdoms of life. However, the *TSC22D* gene family first appears at the metazoan branch of evolution, with exceptions including *N. vectensis* (starlet sea anemone) and *H. magnipapillata* (hydra) (**Fig. 4D** and **S6E**). Despite our best efforts, evidence for TSC22D proteins were not found in choanoflagellates, filasteria or icthyosporea – the closest unicellular organisms related to multicellular animals. Correspondingly, BLASTP analysis of the human NbrT described above shows conservation only in metazoans (E<0.05), except for *C. briggsae* (**Fig. 4D**). The maximum BLASTP^57^ score between the NbrT from human NRBP1 and qualifying metazoan species ranges from 174 (human NRBP1, E=1E-49) to 27.7(*C.elegans* hpo-11, E=5E-05) (**Fig. 4D**), suggesting the binding of NRBP and TSC22D proteins emerged in metazoans, unlike WNK kinases which appear to be more ancient.

We also performed a phylogenetic analysis and show that WNK1 kinase domains co-cluster based on their phylogenetic family. All WNK1 orthologs were next analyzed for ‘IDR fraction’ by IUPRED2a ^58–60^ and this analysis revealed that plant WNK1 orthologs have short polypeptide sequence lengths and low IDR fraction, while metazoans have long WNK IDR sequences and high IDR fraction (**Fig. 4D**). Therefore, the longer WNK IDR regions are positively correlated with the appearance of TSC22D genes in metazoans, consistent with these two gene families co-evolving to encode proteins containing similar IDR sequence features (**Fig. 4B**).

Previous work in model systems indicates that orthologs of *TSC22D*, *WNK*, and *NRBP* genes are involved in cell volume control and osmotic regulation. For example, *Drosophila melanogaster* and *Caenorhabditis elegans* have single orthologs of the TWN genes including *bunched/tsct-1/TSC22D, dWnk/wnk-1/WNK*, and *Madm/hpo-11/NRBP*, respectively (**Data S2**) and a previous study showed that *wnk-1* in worms was important for viability in hyperosmotic conditions ^61^. To extend these observations, we hypothesized that *tsct-1* and *hpo-11* would also be important for surviving hyperosmotic stress and performed a survival assay in *C. elegans* with either *tsct-1*, *hpo-11* or *wnk-1* knocked down using RNA interference. There was no observable difference in viability in wild-type N2 worms with depletion of *tsct-1*, *wnk-1* or *hpo-11* under normal culture conditions; however, knockdown of these genes resulted in substantially reduced viability under hyperosmotic conditions (**Fig. 4E**). These results support the idea that TWN genes are general regulators of hyperosmotic stress across multicellular animals.

### *WNK* inhibition triggers *TWN* body formation in vivo

Decisive evidence indicates that WNK signaling acts upstream of OXSR1, NKCC1 and p38-MAPK for RVI in response to hyperosmotic stress^31^, but the role of TSC22D or NRBP proteins in this signaling cascade is poorly understood. To examine how key cell signalling pathways impact the formation of TWN bodies, we screened endo-tagged *TSC22D2^mScarlet^* HAP1 cells for TSC22D2 foci formation with 21 different small molecule inhibitors targeting common cell signaling pathways (**Data S3)**. Strikingly, the pan-WNK inhibitor WNK463, which inhibits WNK proteins at nanomolar concentrations *in vitro*^32^ and micromolar concentrations in cells^31^, was found to trigger the formation of TSC22D2 foci in the absence of hyperosmotic stress (**Fig. 5A**, **Data S3**). The kinetics of TSC22D2 foci formation was delayed with WNK463 in comparison to salt- or sorbitol-induced TSC22D2 foci formation and occurred between 10-30 minutes following the addition of WNK463 to cells (**Fig. 5B** and **S7A**). WNK463-induced TSC22D2 foci also contained NRBP1 (**Fig. 5B** and **S7A**). Similar observations were made with the small molecule inhibitor WNK-IN-11 (**Fig. S7B**).

**Figure 5.**
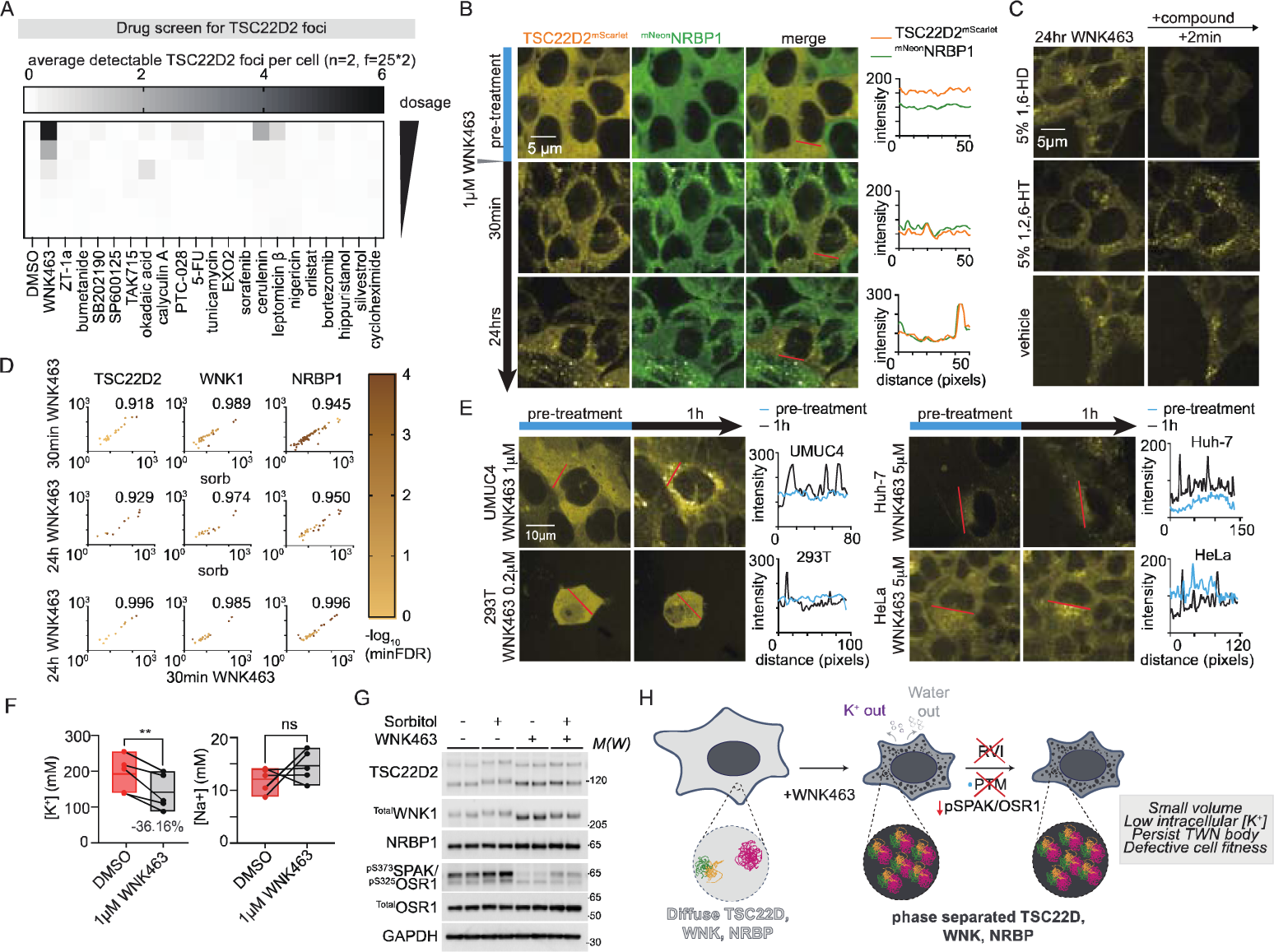
WNK inhibition triggers formation and persistence of TWN bodies. **A**. Heatmap showing a mini microscopy screen for chemical induction of TWN bodies. *TSC22D2^mScarlet^*HAP1 cells were treated with increasing doses of the indicated compounds and the formation of TSC22D2 foci was monitored by automated image analyses with Spotcount. See Data S3 for details of drugs and drug doses. **B.** Images from time-lapse microscopy following the addition of 1µM WNK463 to *^mNeon^NRBP1:TSC22D2^mScarlet^*HAP1 cells. **C.** Live cell imaging after 24 hrs treatment of *TSC22D2^mScarlet^*HAP1 cells with 1µM WNK463 at the indicated doses followed by treatment with 5% 1,6-HD or 5% 1,2,6-HT and imaging 2 minutes later. **D.** Scatterplots and pairwise correlation comparisons for the results of the proximity labelling experiments using endogenously tagged TSC22D2, WNK1, and NRBP1 baits. To be thorough, PCCs are calculated from preys with BFDR<0.25 and indicated in and indicated in the top right of each scatterplot. **E.** Representative microscope images and intensity line plots for alternative endogenously tagged *TSC22D2^mScarlet^*cell lines including UM-UC-4, HEK 293T, Huh-7 and HeLa-Kyoto before and after treatment with WNK463 at indicated doses. **F.** Box-and-Whisker plot for change in [K^+^]_intracellular_ and [Na^+^]_intracellular_ following treatment of parental HAP1 cells with WNK463. **G**. Immunoblotting for HAP1 parental cells treated with no treatment, 1hr 460mM sorbitol and/or 24hrs 10μM WNK463 in two biological replicates. 24hrs treatment of 10μM WNK463 prevents sorbitol-induced protein modification of TSC22D2 and WNK1, and phosphorylation levels of SPAK/OSR1. **H.** Model describing the mechanism of WNK inhibition drives the formation of non-functional TWN bodies and results in failure of cell volume control. See also Fig. S7 and S8.

Both WNK463- and WNK-IN-11-induced TSC22D2 foci disappeared following 1,6-HD treatment, indicating that these structures may be liquid-liquid phase separated foci (**Fig. 5C** and **S7B**). To check the composition of WNK kinase inhibitor-induced foci, a second set of BioID experiments using *^mT^TSC22D2*, *^mT^WNK1*, and *^mT^NRBP1* HAP1 cells was performed in four different conditions: untreated; 30 minutes + sorbitol; 30 minutes + WNK463; and 24 hrs + WNK463. BioID associations across all the experimental treatments with all three baits correlated strongly following the treatment of the cell lines with either sorbitol or WNK463 for different lengths of time (**Fig. 5D**, **S7C**). Together, these observations indicate that the composition of the biomolecular condensates formed by sorbitol or WNK463 treatment is comparable (**Fig. S7D**, PCC>0.97), and that TWN bodies are formed by active inhibition of WNK kinases. WNK463 also induced the formation of TSC22D2 foci in multiple other endo-tagged *TSC22D2^mScarlet^* cell lines including UM-UC-4, HEK293T, Huh-7 and HeLa, indicating that this may be a general mechanism for the formation of TWN bodies across cell types (**Fig. 5E**).

WNK463 treatment also causes a significant decrease in [K^+^]_intracellular_ but not [Na^+^]_intracellular_ (**Fig. 5F**), coupled with a decrease in cell volume and no observable formation of stress granules (**Fig. S1G-H, S7E**). This suggests that TWN body formation is triggered by tonic imbalance. Notably, in an osmotically balanced sorbitol solution, which causes ion and water efflux without changing osmolarity, we observed induced TWN body formation (**Fig. S7F**). Moreover, water efflux caused by a combination of diuretics also induced visible but transient TWN bodies (**Fig. S8A**). The persistence of TWN bodies induced by chronic WNK463 treatment (**Fig. 5B** and **S7A**) is coupled with reduced size-upshift (i.e. PTM modifications) on TSC22D2 and WNK1, as well as downregulation of ^pS373^SPAK/^pS325^OSR1 (**Fig. 5G**), consistent with the idea that WNK kinase activity is needed for RVI and the dissolution of TWN bodies (**Fig. 5H**).

### Chemical-genetics reveals extensive buffering between TSC22D, WNK and NRBP gene families

To survey genetic dependencies following acute chemical inhibition of WNK kinases, we performed a genome-wide chemical genetic pooled CRISPR screen in the presence of 2.5µM of WNK463 (**Fig. S8B**), which is cytotoxic with an IC_50_∼5 µM in HAP1 cells. The strongest negative chemical-genetic interaction with WNK463 was the perturbation of *WNK1* (**Fig. 6A**; FDR=5.38E-8). We also observed a negative chemical-genetic interaction with *NRBP1*, providing the first evidence that *NRBP1* buffers acute chemical inhibition of *WNK* genes. This indicates that reducing WNK1 activity or NRBP1 levels renders cells highly sensitive to WNK inhibition with WNK463. Drug sensitivity assays in parental HAP1 cells and two single mutant clones of *WNK1* and *NRBP1*, as well as two unique *TSC22D1:TSC22D2:TSC22D4* triple mutant clones confirmed these results. That is, the IC_50_ for WNK463 decreased from 4.5µM(WT) to an average of 0.08µM for *WNK1* mutant clones and 0.56µM for *NRBP1* mutant clones (**Fig. 6B**). Although *TSC22D* single gene interactions were not detected in the genome-wide knockout screen for genetic modifiers of WNK463 cytotoxicity, we found that *TSC22D1:TSC22D2:TSC22D4* triple mutant cells showed strong sensitivity to WNK463 cytotoxicity and the IC_50_ dropped to an average of 0.30µM in these triple mutant clones (**Fig. 6B**). Since WNK463 impairs WNK kinase activity, reduces cell volume, and triggers the formation of TWN bodies, the synergistic fitness effect between WNK463 and *WNK1* or *NRBP1* single mutant HAP1 cells, or *TSC22D1:TSC22D2:TSC22D4* triple mutant HAP1 cells, indicates extensive genetic buffering in the regulation of cell RVI.

**Figure 6.**
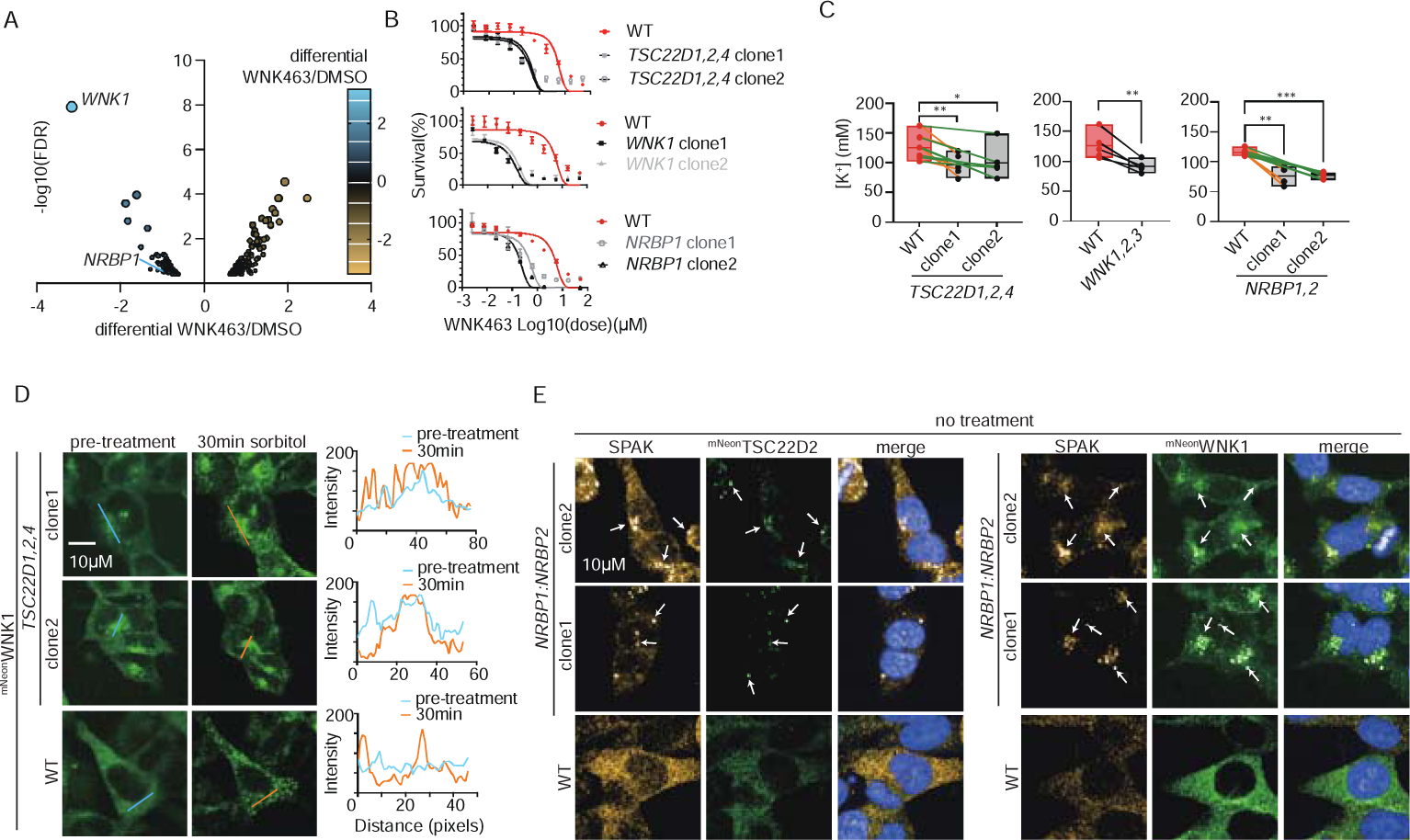
Chemical-genetics with WNK463 reveals buffering between *TSC22D, WNK* and *NRBP* gene families. **A.** Volcano plot showing the results of the genome-wide WNK463 drug screen. **B.** WNK463 dose-response survival curves for different *TSC22D1:TSC22D2:TSC22D4* triple mutant clones, *WNK1* single mutant clones, and *NRBP1* single mutant clones as indicated. **C**. Box-and-Whisker plot for [K^+^]_intracellular_ in parental HAP1 cells, two *TSC22D1:TSC22D2:TSC22D4* triple mutant clones, one *WNK1:WNK2:WNK3* triple mutant clone, and two *NRBP1:NRBP2* double mutant clones. **D.** Microscope images before and after 30min hyperosmotic stress with 460 mM sorbitol in *^mNeon^WNK1* endo-tagged HAP1 parental cells and two clonally-derived *TSC22D1:TSC22D2:TSC22D4* triple mutant cell lines as indicated. The line graphs to the right of each pair of images represent intensity values as a function of the blue lines (pre-treatment) and orange lines (30min sorbitol) as indicated. **E.** Microscope images of *^mNeon^TSC22D2* or *^mNeon^WNK1* endo-tagged (green) signals in HAP1 parental cells and two clonally-derived *NRBP1:NRBP2* double mutant cell lines as indicated. Blue signal represents DAPI stain (i.e. nucleus) and orange signal represents immuno-stained SPAK kinase.

To evaluate between-family genetic interactions across *TWN* genes, we constructed *WNK1:WNK2:WNK3* triple mutant and *NRBP1:NRBP2* double mutant HAP1 cells by selecting for long term clonal survivors. The *TSC22D1:TSC22D2:TSC22D4* and *WNK1:WNK2:WNK3* triple mutant cells, as well as the *NRBP1:NRBP2* double mutant cells, are significantly smaller, have slower doubling rates, and show a net efflux of cations (i.e., reduced intracellular [K^+^]), while the intracellular [Na^+^] remains unchanged (**Fig. 6C** and **S8C**). The observation that *TSC22D1:TSC22D2:TSC22D4*, *WNK1:WNK2:WNK3*, and *NRBP1:NRBP2* mutant cells are viable in standard culture conditions suggests a high degree of genetic buffering that can occur for cell fitness.

As WNK kinase inhibition with WNK463 or WNK-IN-11 results in TWN-body formation (**Fig. 5B** and **S7A-B**), we postulated that mutations in WNK kinases or NRBP pseudokinases would result in diminished phosphorylation activities during RVI. Mutations of *TSC22D1:TSC22D2:TSC22D4*, which failed to partition NRBP1 into TWN-bodies (**Fig. 3B**), led to visible ^mNeon^WNK1 foci formation in iso-osmotic conditions (**Fig. 6D**). Similarly, mutations of *NRBP1:NRBP2* also lead to visible ^mNeon^TSC22D2 and ^mNeon^WNK1 foci formation containing SPAK and OSR1 (**Fig. 6E and S9A**) under standard growth conditions (i.e. no osmotic stress). TSC22D2^mScarlet^ foci formation shows similar kinetics with wild-type in WNK1 depleted cells (**Fig. S9B**), supporting paralog redundancy amongst WNK genes. In contrast, visible TSC22D2 and NRBP1 phase-separated condensates in *WNK1:WNK2:WNK3* triple mutant HAP1 cells are only detectable at high doses of sorbitol (**Fig. S9C**), but severely compromised at lower doses (**Fig. S9D**). These data indicate that TWN bodies are exquisitely sensitive to concentrations of the components driving phase separation, with a sharp switch-like boundary with dependence on the collective concentrations of the contributing proteins to reach the necessary “saturation concentration” for phase separation, consistent with multi-component condensates.

## DISCUSSION

TSC22D family proteins were first identified as transcription factors due to the conserved presence of a leucine zipper motif and their repressive activity when fused to a heterologous DNA-binding domain^62^. However, no direct evidence has been reported for the transcription regulation activity of TSC22D family proteins under physiological conditions. While TSC22D1 (TSC-22) and TSC22D3 (GILZ) are the most studied of the TSC22D proteins (reviewed by Pépin et al^63^), more recent studies have demonstrated a role for TSC22D4 in liver metabolism and hepatic metabolic dysfunction in liver cancer models^64–66^. Little is known about TSC22D2 except for its possible suppressive role in colorectal tumorigenesis^67^. Although *TSC22D* genes may have distinct roles in specific tissues or cell types, we find the protein sequence features and expression patterns^63^ shared by the *TSC22D* gene products (at least for TSC22D1, TSC22D2, and TSC22D4) show striking similarities and behave like cytoplasmic crowding sensors. Our work demonstrates that the *TSC22D* gene family is a stark example of evolution creating redundancy amongst the TSC22D proteins to allow intermingling with NRBPs and WNKs for cytoplasmic crowd sensing and cell volume regulation. Finding the underlying molecular features that drive TWN body formation will help to further clarify the foundation of cytoplasmic crowd sensing in metazoan cells.

It is typical in studies of biocondensates to concentrate on a singular protein entity and scrutinize the modulatory influence exerted by its discrete domains, often characterized by intrinsic disorder, on conditional phase separation dynamics^68^, such as with WNK1^9^, NFAT5^69^, CFTR^70^, FUS and TDP-43^71^. Here we report a distinctive scenario wherein a protein family whose members are mostly disordered (i.e. TSC22D1, TSC22D2, and TSC22D4) constitutively interact with the NRBP pseudo-kinases through distinct sequence regions that evolved in metazoans we have termed TbrN (TSC22D binding region with NRBP) and NbrT (NRBP binding region with TSC22D). The emergence of TSC22D family proteins, long C-terminal IDRs encoded within WNK kinases, and NbrT exhibit a pattern of co-evolution exclusive to metazoans. Our finding that TWN bodies are formed through inhibition of WNK kinases provides new insight into what triggers the formation of WNK containing condensates. It is thought that chloride stabilizes the inactive conformation of WNKs, preventing kinase autophosphorylation and activation, rendering WNKs intracellular chloride sensors through direct binding of a regulatory chloride ion to the active site^72–74^. Thus, large increases in intracellullar chloride are thought to promote WNK inhibition, which is also consistent with the formation of TWN bodies.

Additionally, it’s crucial to note that decades of research on WNK kinases, the downstream effector kinases OXSR1 and SPAK, and the cation chloride co-transporters (NCCs, NKCCs, and KCCs) have failed to uncover functional links with TSC22D and NRBP family members. Gene co-functionality or co-essentiality studies from functional screening data is a powerful concept to link genes to functions for systematic annotation efforts.

Lastly, the evolution of multicellularity in animals required a series of key innovations. It is worth noting that the functionally dominant genes and their families that we identified in our co-essentiality analysis fall into several cellular functionalities that emerged at animal multicellularity including cell signalling and regulation, proteins underlying cellular differentiation that are rich in intrinsically disordered regions, and many proteins that regulate alternative splicing and post-translational modifications on proteins ^75^. The evolution of regulatory circuitry for controlling macromolecular crowding and cell volume in response to hyperosmolarity is a major innovation for metazoans and TWN bodies appear to be a key component of cell volume control.

## LIMITATIONS OF THE STUDY

There are several limitations to the study. First, genetic interactions are inferred from pooled genome-wide CRISPR screens where single mutant fitness profiles in wild-type or parental HAP1 cells are compared to single mutant fitness profiles from clonally derived isogenic mutant HAP1 cell lines. One caveat to this approach is that isogenic HAP1 query mutant cell lines may be amplified under selective pressure introduced by the mutation. This scenario could elicit adaptive dependencies that may not be observed in the acute situation where multiple mutations are introduced simultaneously, or collateral genomic changes that would create a more complex background. We also note that acute di- or tri-genic mutations may not provide insight into the adaptive mechanisms at play for cell fitness. Second, our parental HAP1 cells do not express *WNK4* at detectable levels. We anticipate that WNK4 is also part of TWN bodies, as the work of Boyd-Shiwarski^9^ has demonstrated that WNK1 biomolecular condensates also contain WNK4, as well as OXSR1 and SPAK. Third, *TSC22D3* is an outlier in the *TSC22D* family, in terms of function and predicted protein structure. For example, we do not detect TSC22D3 as a genetic interaction with other TSC22D family members. *TSC22D3* appears to have strong co-dependency profiles in DepMap data but is not essential in any cell line to our knowledge. TSC22D3 is largely predicted to be an alpha-helix and is 134 amino acids in length, in contrast to TSC22D1 (1073 aa), TSC22D2 (780 aa) and TSC22D4 (395 aa) which largely contain intrinsically disordered sequence and a predicted alpha-helix. Nevertheless, we do detect proximity interactions between TSC22D3 (prey) and endo-tagged NRBP1 (bait), so it remains possible that TSC22D3 has residual function in HAP1 cells. Moreover, TSC22D3 interactions with NRBPs have been detected in large-scale protein interaction studies using overexpressed bait proteins and co-immunoprecipitation followed by western blotting^38–41,76^. Fifth, we did not detect ASK3(*MAP3K15*) as a genetic or physical interaction in our study, despite detectable mRNA levels for *MAP3K15*. ASK3 was previously reported to phase separate and to bind and suppress WNK1 and SPAK/OSR1 function in the kidney^77–79^.

## ACKNOWLEDGEMENTS

We thank all members of the Moffat lab for advice and comments on the manuscript, Benjamin Petite for help visualizing PPI data in Cytoscape, Michael Nosella and Tai Hun Kim from the Forman-Kay lab for experimental advice, and Brett Larsen and Cassandra Wong from the Gingras lab for mass spectrometry advice.

## AUTHOR CONTRIBUTIONS

Conceptualization: YXX, JM

Methodology: YXX, RS, SYL, MAU, AA, ZTL, RC, KC, PM, CM, AM, WBD, DR, IP, CY, ACG, JM

Investigation: YXX, RS, SYL, MAU, AA, YK, ZTL, RC, KA, JW, AGF, TR, AH, JQH, MB, MR

Visualization: YXX, JM, PM, SYL

Funding acquisition: JM, CM, BA, ACG, JYY, AMM, JDF, WBD Project administration: JM

Supervision: CM, BJA, JYY, DR, WBD, JDF, AM, IP, ACG, JM

Writing – original draft: YXX, JM

Writing – review & editing: YXX, JM, SYL, PM, KC, KB, RS, ZTL, AGF, BJA, JYY, CY, DR, WBD, JDF, AM, IP, ACG

## DATA AND MATERIALS AVAILABILITY

All data are available in the main text or the supplementary materials.

---

**Fig. S1:**
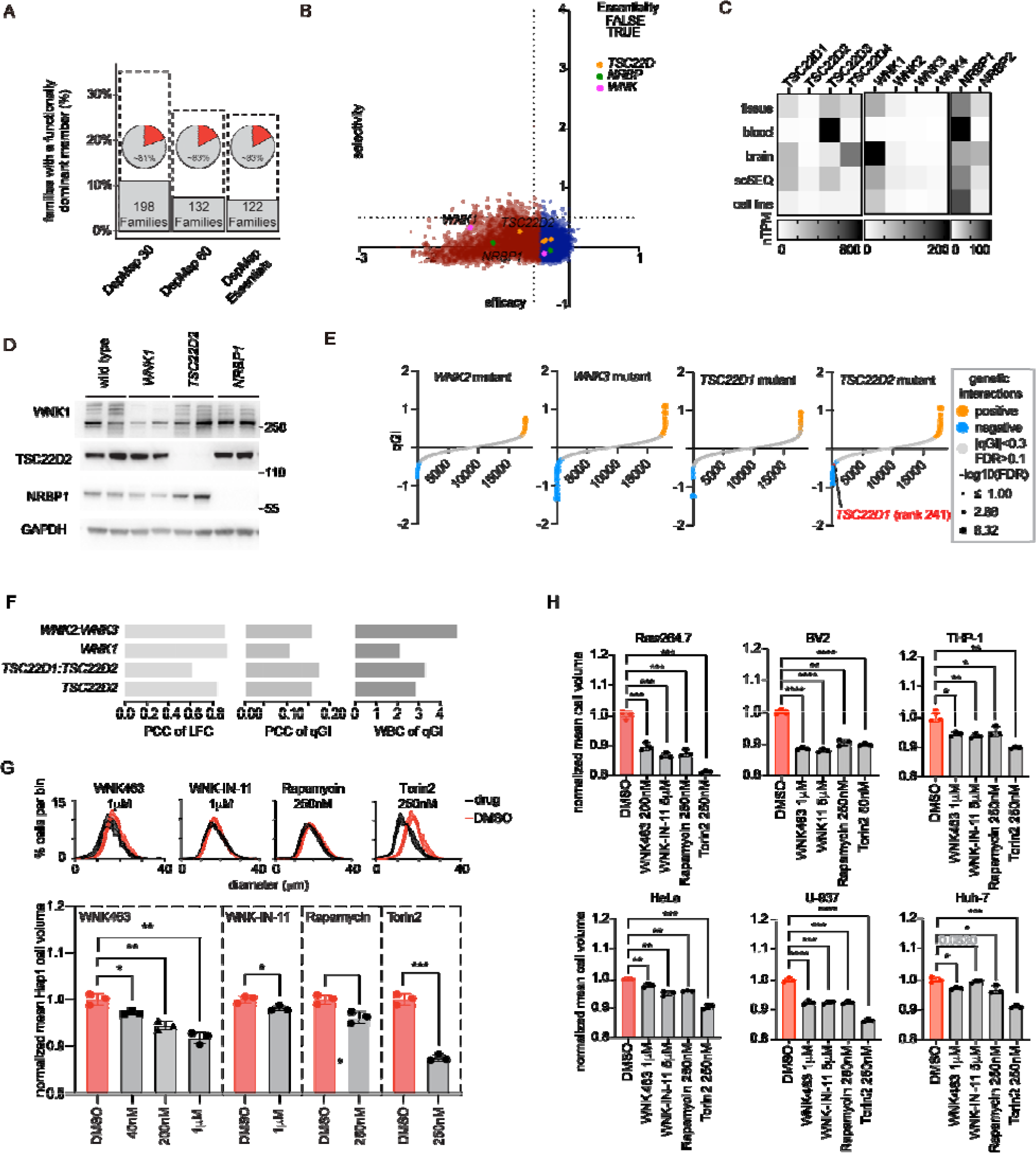
*TSC22D2*, *WNK1* and *NRBP1* are functionally dominant paralogs in human cells, related to Fig. 1. **A.** Bar plot showing the percentage of functionally dominant ohnolog gene families as a function of essentiality in the DepMap CRISPR data, where background gene essentiality is indicated by the dotted bar. Pie charts indicate the percentage of functionally dominant genes that are the highest expressed in each class where an average of 9.5% (max=11%, min=7%) of multigene families include a single essential gene. **B.** ShinyDepMap shows *TSC22D2, WNK1,* and *NRBP1* have the highest absolute efficacy in their sensitive cell lines among their paralogues(*56*). X-axis represents efficacy (i.e., the degree to which perturbation of the gene reduces cell growth in sensitive lines) and y-axis represents selectivity (i.e., the degree to which its essentiality varies across lines). Legend defining dot colour is shown on the right. **C.** Human Protein Atlas RNA expression for *TSC22D, WNK,* and *NRBP* paralogs from tissue, blood, brain, single-cell sequencing (scSEQ), and cell line datasets. **D.** Immunoblot confirmation of protein levels in *WNK1*, *TSC22D2* and *NRBP1* mutant HAP1 query cell lines (n=2). **E.** Rank plot of qGI scores for representative *WNK2*(n=1), *WNK3*(n=1)*, TSC22D1*(n=1), and *TSC22D2*(n=2) single mutant HAP1 query screens. Legend defining dot size and dot colour is shown on the right. **F.** Reproducibility of two biological replicates of *WNK1*, *TSC22D2, WNK2:WNK3*, and *TSC22D1:TSC22D2* mutant query screens using the within-between correlation (WBC) score described in(*12*). **G.** Distribution plots and bar plots for the cell size data from **Fig. 1G**. WNK463, WNK-IN-11, rapamycin and Torin2 treatments are shown in greys and are performed for 24hrs with indicated dosages. DMSO treatment distributions are shown in red for comparison. **H.** Bar plots of normalized mean cell volume of RAW264.7, BV-2, THP-1, HeLa-Kyoto, U-937, and Huh-7 cells. Drug treatments were performed for 24hrs.

**Fig. S2:**
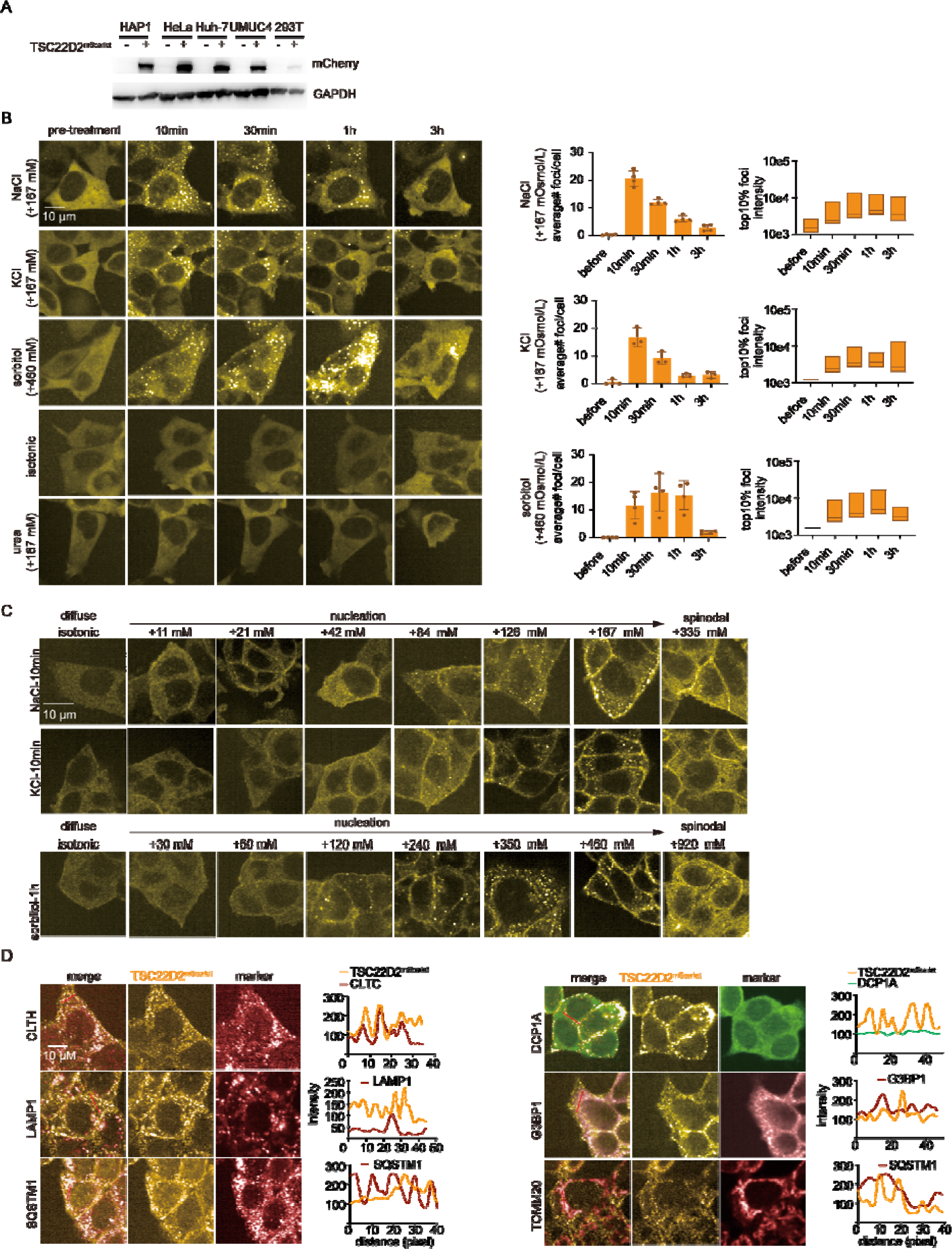
Kinetics of TSC22D2 foci formation, related to Fig. 2. **A.** Immunoblots for mCherry polyclonal antibody to detect mScarlet that is endogenously tagged to the c-terminus of TSC22D2 in indicated cell lines. GAPDH served as the loading control. **B.** The left panel shows time-lapse images of HAP1 *TSC22D2^mScarlet^*cells following the indicated hypertonic stress including NaCl, KCl, and sorbitol. Small molecule crowding reagent urea is used for comparison. The right panel shows the quantification of average TSC22D2^mScarlet^ foci number per cell and the intensities of top 10% of TSC22D2^mScarlet^ foci under indicated conditions. **C.** Representative microscope images for TSC22D2^mScarlet^ foci formation under indicated concentrations of hypertonic stress. **D.** Representative microscope images for endo-tagged TSC22D2^mScarlet^ with known sub-cellular markers of distinct compartments including clathrin heavy chain (CLTC) for endosomes, LAMP1 for lysosomes, SQSTM1 for aggresomes, DCP1A for RNA granules, G3BP1 for stress granules and TOMM20 for mitochondria. The line graphs to the right represent intensity values as a function of the red line indicated in the merged images. The colors in the line graphs correspond to the proteins and are indicated in the legends. Pixel intensity show good localization with CLTC which is consistent with previous publications (*3*).

**Fig. S3:**
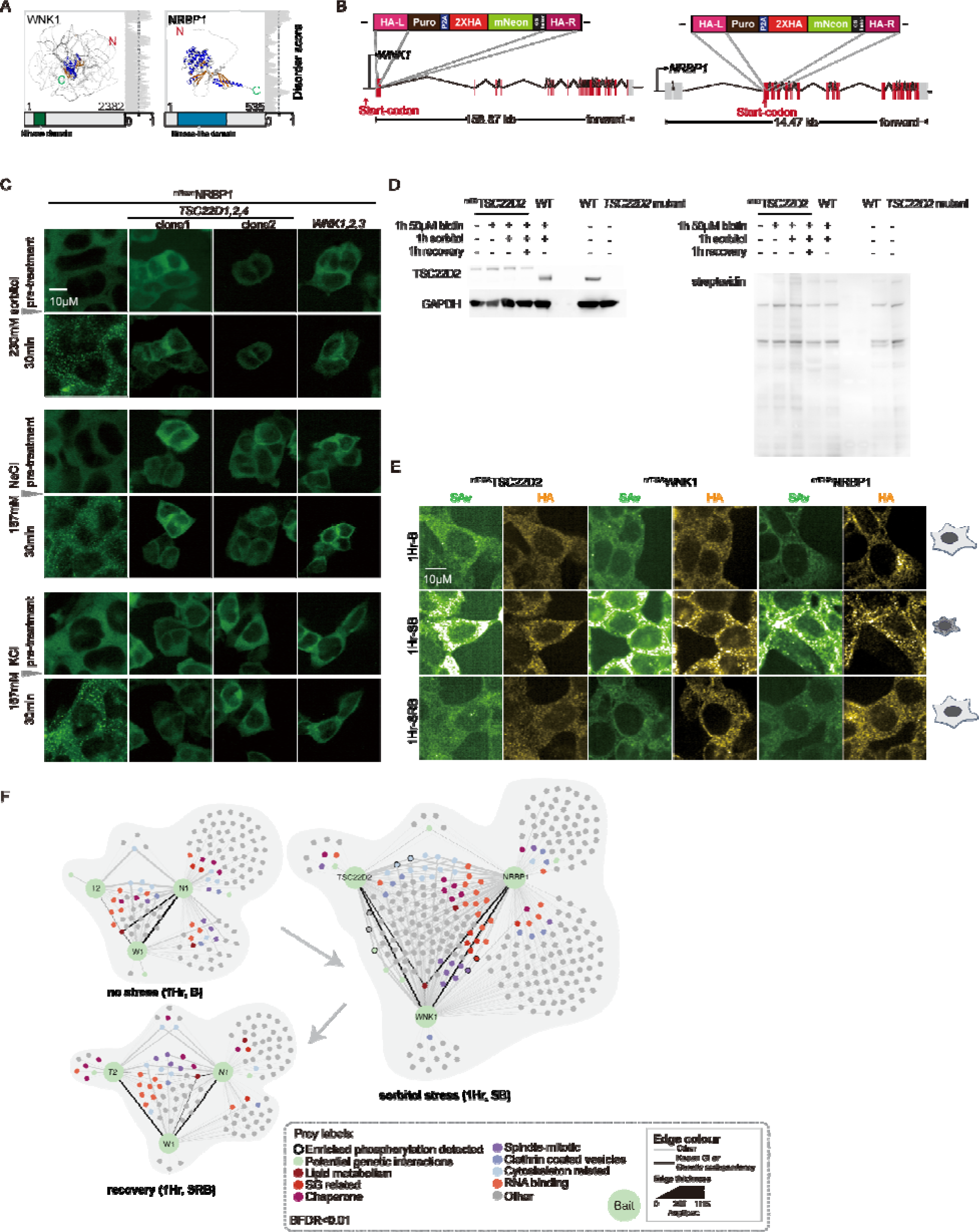
Proximity interactions of TWN proteins, related to Fig. 2 and 3. **A.** Predicted canonical WNK1 and NRBP1 structure. **B.** Strategies for tagging the N-terminus of WNK1 and NRBP1 with mNeon at their endogenous locus; the bottom panel shows. **C.** Representative time-lapse images before and after 30min hypertonic stress with various hypertonicity stress, including 230mM sorbitol, 167mM NaCl, and 167mM KCl, in ^mNeon^NRBP1 endo-tagged HAP1 parental cells and HAP1 cells with *TSC22D1:TSC22D2:TSC22D4* triple gene mutants. **D.** Immunoblotting for ^mT^TSC22D2 HAP1 cells to detect TSC22D2 and streptavidin. Tagged TSC22D2 shows an upshifted band compared with untagged wild-type (WT) while *TSC22D2* mutant line is deficient of TSC22D2. GAPDH served as the loading control. **E.** Co-immunofluorescence staining validates that the mT tags in *^mT^TSC22D2*, *^mT^WNK1*, and *^mT^NRBP1* cells do not interfere with formation of biomolecular condensates following a 1hr treatment with 460 mM sorbitol. **F.** Network analysis of the proximity interactions for each of TSC22D2, WNK1 and NRBP1 baits in the no stress (1Hr, B), sorbitol stress (1Hr, SB) and recovery (1Hr, SRB) conditions. The legend at the bottom defines the node and edge colors, thickness and size.

**Fig. S4:**
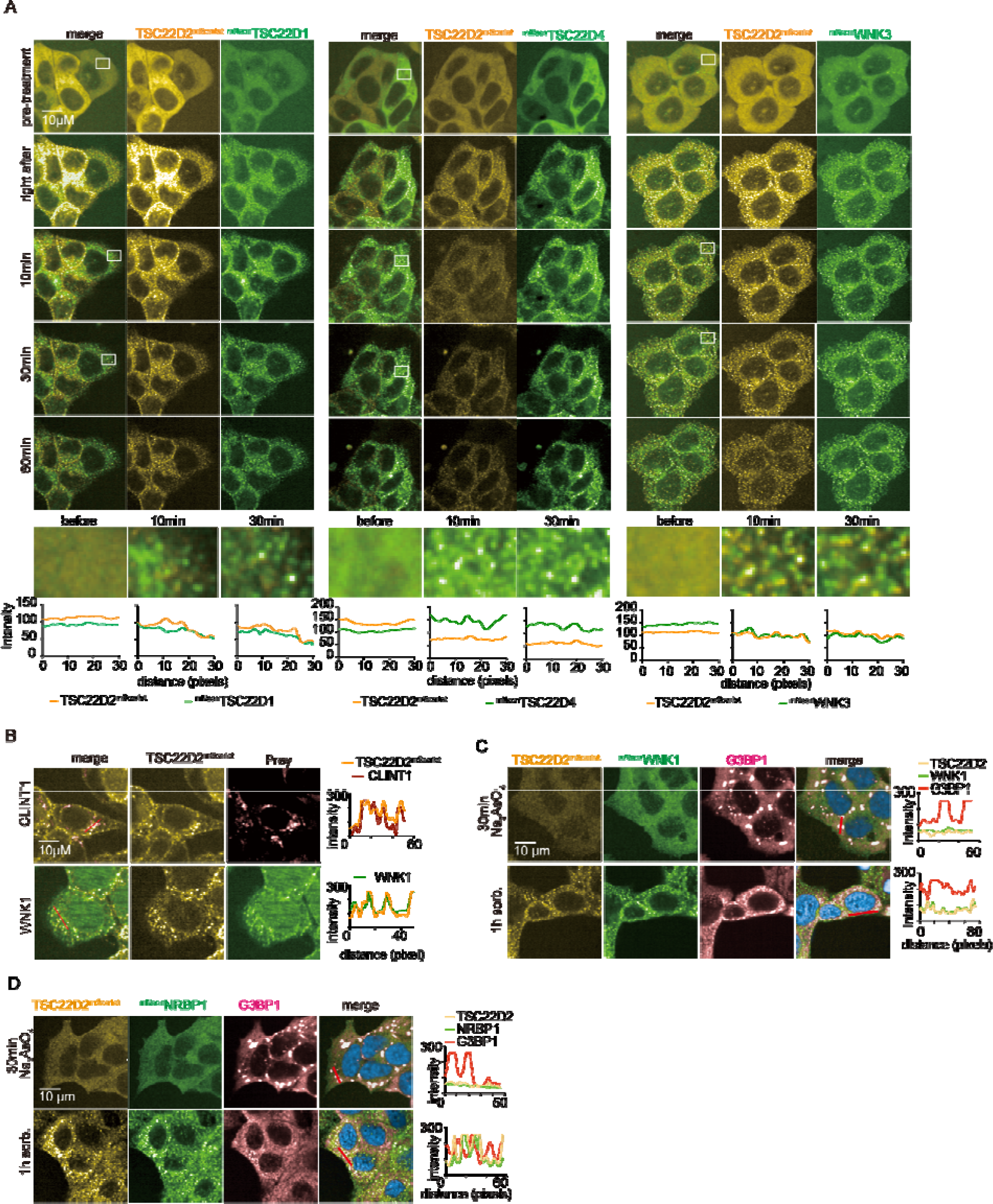
Validation of proximity associations, related to Fig. 3. **A**. Representative time-lapse images for co-tagging N-terminus of *TSC22D1*, *TSC22D4*, or *WNK3* with mNeon in *TSC22D2^mScaret^* HAP1 cells. The line graphs on the bottom represent intensity values as a function of the white box indicated in the merged images. The colors in the line graphs correspond to the proteins and are indicated in the legends. **B.** Representative immunofluorescence images staining preys of interest (i.e., WNK1 and CLINT1) with antibodies detecting endogenous proteins in TSC22D2^mScarlet^ HAP1 cells. The line graphs to the right represent intensity values as a function of the red line indicated in the merged images. The colors in the line graphs correspond to the proteins and are indicated in the legends. Representative immunofluorescence images of *^mNeon^WNK1:TSC22D2^mScarlet^*or *^mNeon^NRBP1:TSC22D2^mScarlet^* HAP1 cells following **C.** 30 min treatment with 0.5 mM sodium arsenate or **D.** 1 hr treatment with 460 mM sorbitol (bottom panel). TSC22D2 (yellow), WNK1 (green), G3BP1 (pink) and the merged images including nuclei (blue) are indicated. The line graphs to the right represent intensity values as a function of the red line indicated in the merged images. The colours in the line graphs correspond to the proteins and are indicated in the legends.

**Fig. S5:**
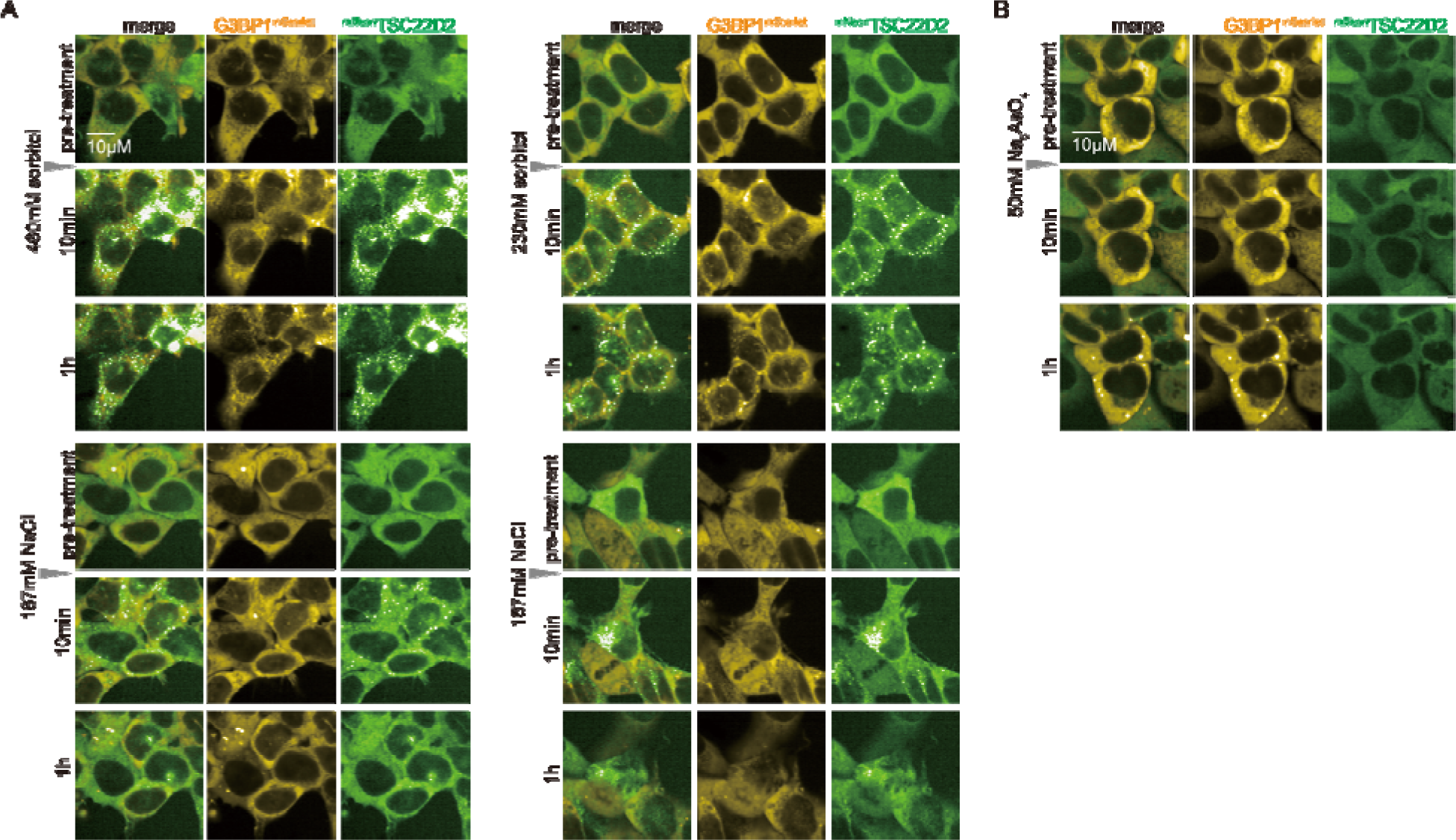
Kinetics of TWN bodies and stress granules (SGs), related to Fig. 3. **A.** Representative time-lapse images for ^mNeon^TSC22D2 and G3BP1^mScarlet^ dual tagged cells under hypertonic stresses including 460mM sorbitol, 230mM sorbitol, 167mM NaCl and 167mM KCl. **B.** Representative time-lapse images for ^mNeon^TSC22D2 and G3BP1^mScarlet^ dual tagged cells under 50mM sodium arsenate.

**Fig. S6:**
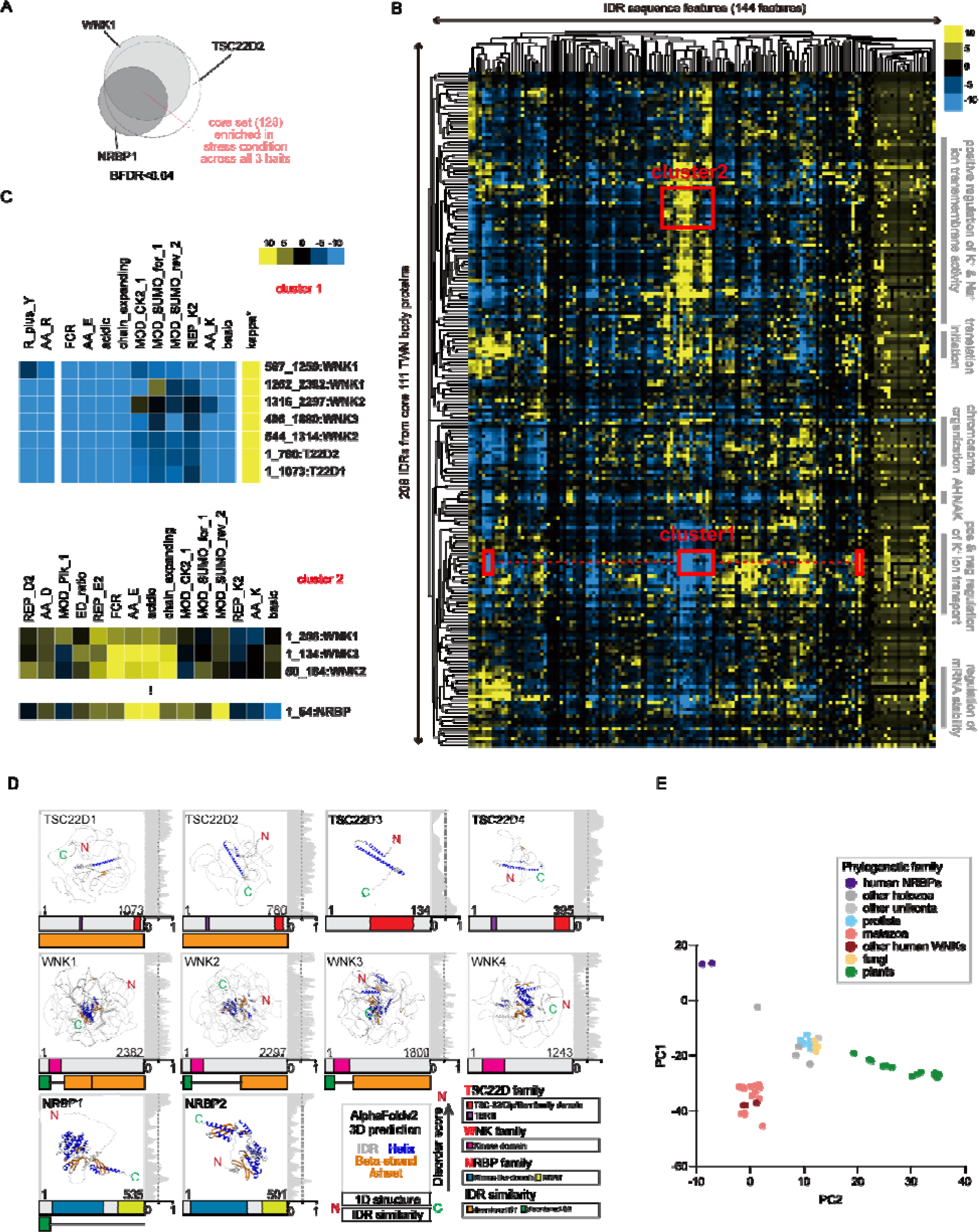
Evolutionary signature and TWN protein structure analysis, related to Fig. 4. **A.** Schematics outlining the core set of proximity associations that were examined for specific sequence features in panel B. **B.** The 144×208 z-score matrix of evolutionary signatures global heatmap with the core set against total human proteome. The z-score legends are selected against (blue) or for (yellow) amongst the indicated IDRs. **C.** Selected portions of heatmap presented in **Fig. S6B** highlighting key evolutionary features that are selected against (blue) or for (yellow) amongst the indicated disordered subsequences containing N terminal WNK1-3 and TSC22D1-2(cluster1, corresponding to disorder S1 in **Fig. S6D**), or WNK1-3 and NRBP (cluster2, corresponding to disorder S2 in **Fig. S6D**). The column labels are defined in **Data S2**. Z-scores were calculated as described in the Methods. **D.** TWN protein sequence features including protein domains summarized using InterPRO, disorder scrores predicted by IUPRED2A, regions of similar IDR features (disordered S1 and S2), and 3-dimensional structures predicted by Alpha-fold-2. **E.** PCA analysis on sequence percentile identity with input WNK kinase domains from 51 model organisms from **Fig. 4D**.

**Fig. S7:**
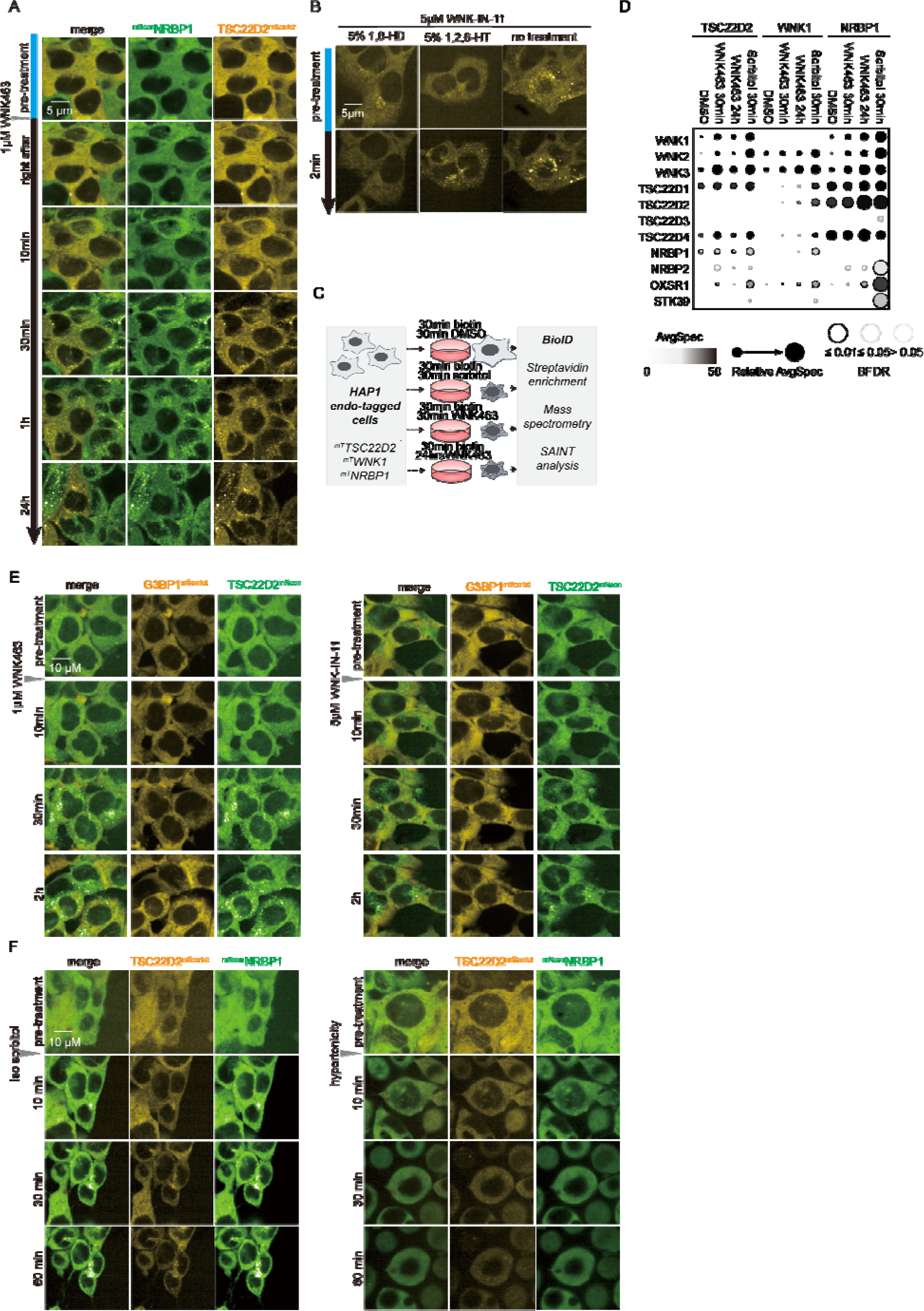
WNK463 inhibition and ion imbalance trigger TWN body formation, related to Fig. 5. **A.** Images of additional time-lapse microscopy following the addition of 1 mM WNK463 to *^mNeon^NRBP1:TSC22D2^mScarlet^*HAP1 cells. **B.** Live cell imaging after 24hrs treatment of *TSC22D2^mScarlet^*HAP1 cells 5µM WNK-IN-11 at the indicated doses followed by treatment with 1,6-HD or 1,2,6-HT and imaging 2 minutes later. **C.** Experimental setup for condition-specific proximity mapping with 1µM WNK463 or 460mM sorbitol of TSC22D2, WNK1 and NRBP1 associations in HAP1 cells with endogenous N-terminal miniTurbo (mT) tags. **D.** Dot-plot graph showing recovery of TSC22D, WNK, and NRBP paralogs as preys, as well as OXSR1 and STK39 under indicated conditions. Dots are plotted as a function of the Bayesian False Discovery Rate or BFDR and spectral counts, as indicated in the legend. **E.** Representative time-lapsed microscope images for endogenously dual tagged G3BP1^mScarlet^ and ^mNeon^TSC22D2 cells following 1µM WNK463 (left) or 5µM WNK-IN-11 (right) treatment. WNK kinase inhibition induces TWN body formation but not SG. **F.** Images from time-lapse microscopy following the addition of hypotonic medium (IMDM:H_2_O=1:1) (left) or Isotonic (iso) sorbitol solution (300mOsmol/L) (right) to *^mNeon^NRBP1:TSC22D2^mScarlet^*HAP1 cells.

**Fig. S8:**
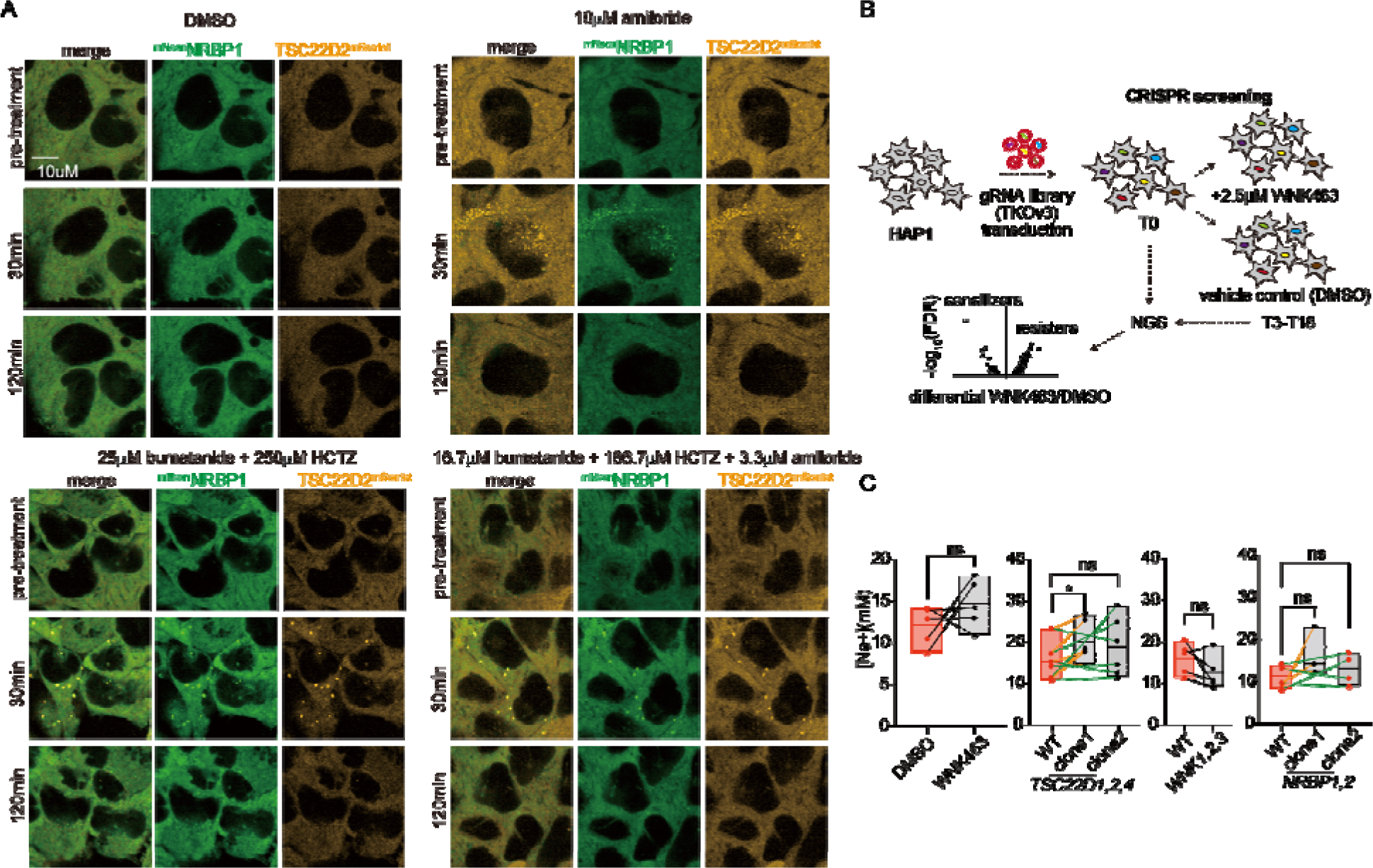
TWN body is induced by tonicity imbalance, related to Fig. 5 and 6. **A.** Representative time-lapse images of *^mNeon^NRBP1:TSC22D2^mScarlet^* cells treated with amiloride, a combination of bumetanide and hydrochlorothiazide (HCZT) and a combination of bumetanide, HCZT and amiloride with indicated dosages and time. **B.** Schematics for WNK463 chemical-genetics CRISPR screening. gRNA library(TKOv3) were transduced to HAP1 wild-type cells at MOI of 0.3. Transduced cell populations were then purified with puromycin selection and split at T0 into two arms. One arm is treated with 2.5µM WNK463 and the other arm is treated with vehicle control DMSO all the way to T18. T0 and T18 cell pellets are subjected to next-generation sequencing for determination of sensitizers and resisters of WNK463. **C**. Box-and-Whisker plot for change in [Na^+^]_intracellular_ comparing *TSC22D1:TSC22D2:TSC22D4*, *WNK1:WNK2:WNK3*, and *NRBP1:NRBP2* mutant clones and HAP1 parental cells.

**Fig. S9:**
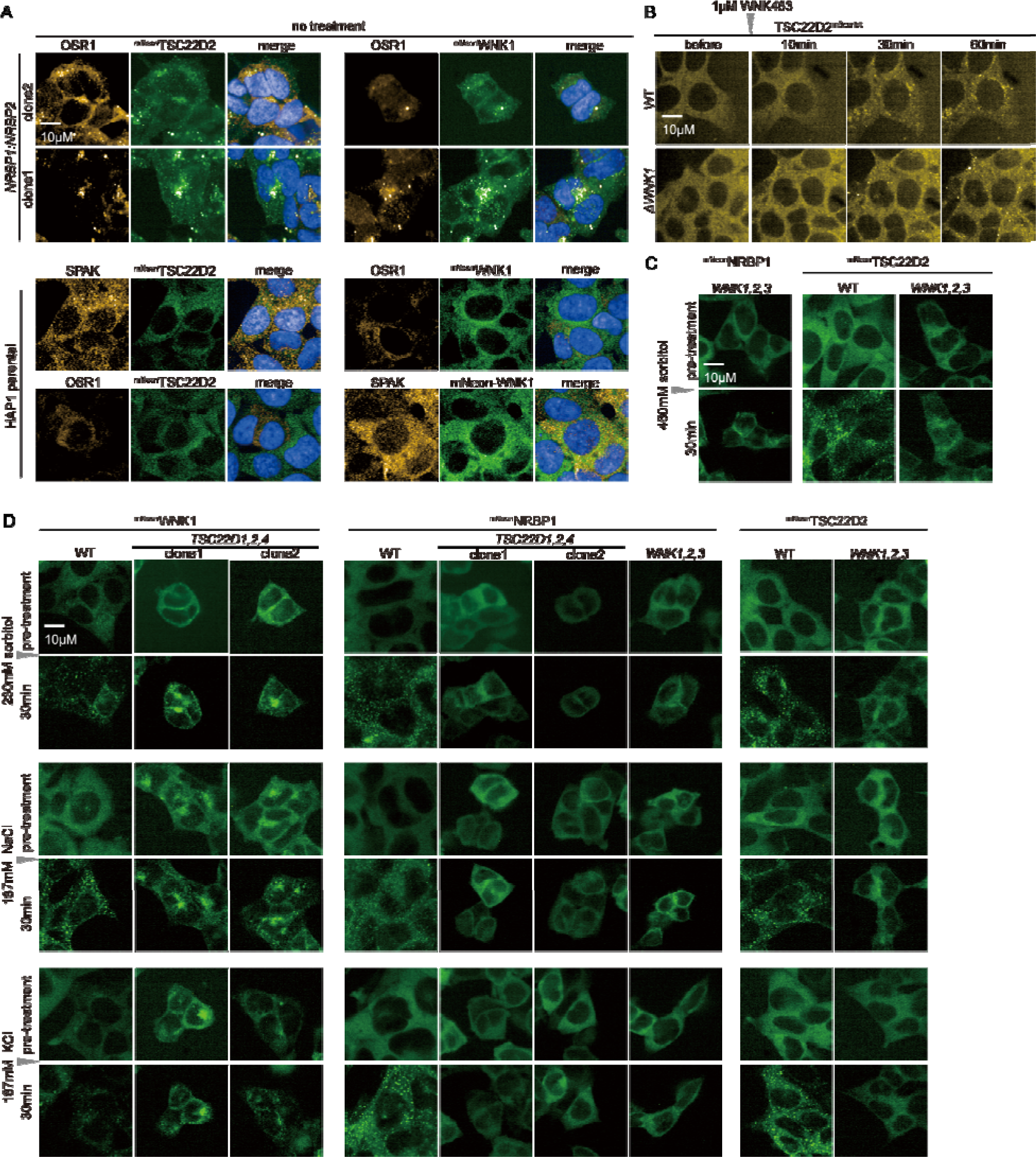
TSC22Ds, WNKs and NRBPs are essential for proper TWN body formation, related to Fig. 6. **A.** Microscope images of ^mNeon^TSC22D2 or ^mNeon^WNK1 endo-tagged (green) HAP1 parental cells and *NRBP1:NRBP2* mutant cells. Blue signal represent DAPI(nucleus), and the orange signal represents co-stained SPAK or OSR1 kinases. **B.** Representative time-lapse images showing TSC22D2^mScarlet^ foci still form in WNK1 depletion cells after the addition of 1µM WNK463. **C.** Representative time-lapse images showing ^mNeon^NRBP1 and ^mNeon^TSC22D2 foci are compromised in *WNK1:WNK2:WNK3* triple depletion cells after the addition of 460mM sorbitol. **D.** Representative time-lapse images showing ^mNeon^NRBP1 and ^mNeon^TSC22D2 form compromised foci in *WNK1:WNK2:WNK3* mutant under 30min treatment of 230mM sorbitol, 167mM NaCl or 167mM KCl. ^mNeon^WNK1 forms foci under isotonic stress in *TSC22D1:TSC22D2:TSC22D4* mutant cells.

## REFERENCES

1. Ponder, E., and Saslow, G. (1930). The measurement of red cell volume: II. Alterations in cell volume in solutions of various tonicities. J Physiol 70, 169–181. 10.1113/jphysiol.1930.sp002685.

2. MacAulay, N. (2021). Molecular mechanisms of brain water transport. Nat Rev Neurosci 22, 326–344. 10.1038/s41583-021-00454-8.

3. MacAulay, N. (2021). Reply to ‘Aquaporin 4 and glymphatic flow have central roles in brain fluid homeostasis’. Nat Rev Neurosci 22, 651–652. 10.1038/s41583-021-00515-y.

4. Pardo, L.A., and Stuhmer, W. (2014). The roles of K(+) channels in cancer. Nat Rev Cancer 14, 39–48. 10.1038/nrc3635.

5. Verkman, A.S., Anderson, M.O., and Papadopoulos, M.C. (2014). Aquaporins: important but elusive drug targets. Nat Rev Drug Discov 13, 259–277. 10.1038/nrd4226.

6. O’Neill, W.C. (1999). Physiological significance of volume-regulatory transporters. Am J Physiol 276, C995–C1011. 10.1152/ajpcell.1999.276.5.C995.

7. Saddhe, A.A., Karle, S.B., Aftab, T., and Kumar, K. (2021). With no lysine kinases: the key regulatory networks and phytohormone cross talk in plant growth, development and stress response. Plant Cell Rep 40, 2097–2109. 10.1007/s00299-021-02728-y.

8. Verissimo, F., and Jordan, P. (2001). WNK kinases, a novel protein kinase subfamily in multi-cellular organisms. Oncogene 20, 5562–5569. 10.1038/sj.onc.1204726.

9. Boyd-Shiwarski, C.R., Shiwarski, D.J., Griffiths, S.E., Beacham, R.T., Norrell, L., Morrison, D.E., Wang, J., Mann, J., Tennant, W., Anderson, E.N., et al. (2022). WNK kinases sense molecular crowding and rescue cell volume via phase separation. Cell. 10.1016/j.cell.2022.09.042.

10. Parker, J.C. (1993). In defense of cell volume? Am J Physiol 265, C1191–1200. 10.1152/ajpcell.1993.265.5.C1191.

11. Walter, H., and Brooks, D.E. (1995). Phase separation in cytoplasm, due to macromolecular crowding, is the basis for microcompartmentation. FEBS Lett 361, 135–139. 10.1016/0014-5793(95)00159-7.

12. Zimmerman, S.B., and Harrison, B. (1987). Macromolecular crowding increases binding of DNA polymerase to DNA: an adaptive effect. Proc Natl Acad Sci U S A 84, 1871–1875. 10.1073/pnas.84.7.1871.

13. Ellison, D.H. (1993). Pseudohypoaldosteronism Type II. In GeneReviews((R)), M.P. Adam, G.M. Mirzaa, R.A. Pagon, S.E. Wallace, L.J.H. Bean, K.W. Gripp, and A. Amemiya, eds.

14. Mabillard, H., and Sayer, J.A. (2019). The Molecular Genetics of Gordon Syndrome. Genes 10. 10.3390/genes10120986.

15. Rodan, A.R., and Jenny, A. (2017). WNK Kinases in Development and Disease. Current topics in developmental biology 123, 1–47. 10.1016/bs.ctdb.2016.08.004.

16. Ewen-Campen, B., Mohr, S.E., Hu, Y., and Perrimon, N. (2017). Accessing the Phenotype Gap: Enabling Systematic Investigation of Paralog Functional Complexity with CRISPR. Dev Cell 43, 6–9. 10.1016/j.devcel.2017.09.020.

17. Gu, Z., Steinmetz, L.M., Gu, X., Scharfe, C., Davis, R.W., and Li, W.H. (2003). Role of duplicate genes in genetic robustness against null mutations. Nature 421, 63–66. 10.1038/nature01198.

18. Atchley, W.R., Fitch, W.M., and Bronner-Fraser, M. (1994). Molecular evolution of the MyoD family of transcription factors. Proc Natl Acad Sci U S A 91, 11522–11526. 10.1073/pnas.91.24.11522.

19. Laruson, A.J., Yeaman, S., and Lotterhos, K.E. (2020). The Importance of Genetic Redundancy in Evolution. Trends Ecol Evol 35, 809–822. 10.1016/j.tree.2020.04.009.

20. Morimoto, M., Nishinakamura, R., Saga, Y., and Kopan, R. (2012). Different assemblies of Notch receptors coordinate the distribution of the major bronchial Clara, ciliated and neuroendocrine cells. Development 139, 4365–4373. 10.1242/dev.083840.

21. Costanzo, M., Kuzmin, E., van Leeuwen, J., Mair, B., Moffat, J., Boone, C., and Andrews, B. (2019). Global Genetic Networks and the Genotype-to-Phenotype Relationship. Cell 177. 10.1016/j.cell.2019.01.033.

22. Boehm, J.S., Garnett, M.J., Adams, D.J., Francies, H.E., Golub, T.R., Hahn, W.C., Iorio, F., McFarland, J.M., Parts, L., and Vazquez, F. (2021). Cancer research needs a better map. Nature 589, 514–516. 10.1038/d41586-021-00182-0.

23. Dempster, J.M., Pacini, C., Pantel, S., Behan, F.M., Green, T., Krill-Burger, J., Beaver, C.M., Younger, S.T., Zhivich, V., Najgebauer, H., et al. (2019). Agreement between two large pan-cancer CRISPR-Cas9 gene dependency data sets. Nature communications 10, 5817–5817. 10.1038/s41467-019-13805-y.

24. Hahn, W.C., Bader, J.S., Braun, T.P., Califano, A., Clemons, P.A., Druker, B.J., Ewald, A.J., Fu, H., Jagu, S., Kemp, C.J., et al. (2021). An expanded universe of cancer targets. Cell 184, 1142–1155. 10.1016/j.cell.2021.02.020.

25. Meyers, R.M., Bryan, J.G., McFarland, J.M., Weir, B.A., Sizemore, A.E., Xu, H., Dharia, N.V., Montgomery, P.G., Cowley, G.S., Pantel, S., et al. (2017). Computational correction of copy number effect improves specificity of CRISPR-Cas9 essentiality screens in cancer cells. Nature genetics 49, 1779–1784. 10.1038/ng.3984.

26. Pacini, C., Dempster, J.M., Boyle, I., Gonçalves, E., Najgebauer, H., Karakoc, E., van der Meer, D., Barthorpe, A., Lightfoot, H., Jaaks, P., et al. (2021). Integrated cross-study datasets of genetic dependencies in cancer. Nature communications 12, 1661–1661. 10.1038/s41467-021-21898-7.

27. Zagórska, A., Pozo-Guisado, E., Boudeau, J., Vitari, A.C., Rafiqi, F.H., Thastrup, J., Deak, M., Campbell, D.G., Morrice, N.A., Prescott, A.R., and Alessi, D.R. (2007). Regulation of activity and localization of the WNK1 protein kinase by hyperosmotic stress. The Journal of cell biology 176, 89–100. 10.1083/jcb.200605093.

28. Singh, P.P., and Isambert, H. (2020). OHNOLOGS v2: a comprehensive resource for the genes retained from whole genome duplication in vertebrates. Nucleic acids research 48, D724–D730. 10.1093/nar/gkz909.

29. Aregger, M., Lawson, K.A., Billmann, M., Costanzo, M., Tong, A.H.Y., Chan, K., Rahman, M., Brown, K.R., Ross, C., Usaj, M., et al. (2020). Systematic mapping of genetic interactions for de novo fatty acid synthesis identifies C12orf49 as a regulator of lipid metabolism. Nature Metabolism 2. 10.1038/s42255-020-0211-z.

30. Billmann, M., Ward, H.N., Aregger, M., Costanzo, M., Andrews, B.J., Boone, C., Moffat, J., and Myers, C.L. (2023). Reproducibility metrics for context-specific CRISPR screens. Cell Syst 14, 418–422 e412. 10.1016/j.cels.2023.04.003.

31. Liu, Z., Demian, W., Persaud, A., Jiang, C., Subramanaya, A.R., and Rotin, D. (2022). Regulation of the p38-MAPK pathway by hyperosmolarity and by WNK kinases. Scientific reports 12, 14480–14480. 10.1038/s41598-022-18630-w.

32. Yamada, K., Park, H.-M., Rigel, D.F., DiPetrillo, K., Whalen, E.J., Anisowicz, A., Beil, M., Berstler, J., Brocklehurst, C.E., Burdick, D.A., et al. (2016). Small-molecule WNK inhibition regulates cardiovascular and renal function. Nature chemical biology 12, 896–898. 10.1038/nchembio.2168.

33. Eltschinger, S., and Loewith, R. (2016). TOR Complexes and the Maintenance of Cellular Homeostasis. Trends in cell biology 26, 148–159. 10.1016/j.tcb.2015.10.003.

34. Yamada, K., Levell, J., Yoon, T., Kohls, D., Yowe, D., Rigel, D.F., Imase, H., Yuan, J., Yasoshima, K., DiPetrillo, K., et al. (2017). Optimization of Allosteric With-No-Lysine (WNK) Kinase Inhibitors and Efficacy in Rodent Hypertension Models. Journal of medicinal chemistry 60, 7099–7107. 10.1021/acs.jmedchem.7b00708.

35. Shimobayashi, S.F., Ronceray, P., Sanders, D.W., Haataja, M.P., and Brangwynne, C.P. (2021). Nucleation landscape of biomolecular condensates. Nature 599, 503–506. 10.1038/s41586-021-03905-5.

36. Wheeler, J.R., Matheny, T., Jain, S., Abrisch, R., and Parker, R. (2016). Distinct stages in stress granule assembly and disassembly. Elife 5. 10.7554/eLife.18413.

37. Gao, Y., Li, X., Li, P., and Lin, Y. (2022). A brief guideline for studies of phase-separated biomolecular condensates. Nature Chemical Biology 18, 1307–1318. 10.1038/s41589-022-01204-2.

38. Buljan, M., Ciuffa, R., van Drogen, A., Vichalkovski, A., Mehnert, M., Rosenberger, G., Lee, S., Varjosalo, M., Pernas, L.E., Spegg, V., et al. (2020). Kinase Interaction Network Expands Functional and Disease Roles of Human Kinases. Mol Cell 79, 504–520 e509. 10.1016/j.molcel.2020.07.001.

39. Cho, N.H., Cheveralls, K.C., Brunner, A.D., Kim, K., Michaelis, A.C., Raghavan, P., Kobayashi, H., Savy, L., Li, J.Y., Canaj, H., et al. (2022). OpenCell: Endogenous tagging for the cartography of human cellular organization. Science 375, eabi6983. 10.1126/science.abi6983.

40. Huttlin, E.L., Bruckner, R.J., Navarrete-Perea, J., Cannon, J.R., Baltier, K., Gebreab, F., Gygi, M.P., Thornock, A., Zarraga, G., Tam, S., et al. (2021). Dual proteome-scale networks reveal cell-specific remodeling of the human interactome. Cell 184, 3022–3040.e3028. 10.1016/j.cell.2021.04.011.

41. Huttlin, E.L., Bruckner, R.J., Paulo, J.A., Cannon, J.R., Ting, L., Baltier, K., Colby, G., Gebreab, F., Gygi, M.P., Parzen, H., et al. (2017). Architecture of the human interactome defines protein communities and disease networks. Nature 545, 505–509. 10.1038/nature22366.

42. Huttlin, E.L., Ting, L., Bruckner, R.J., Gebreab, F., Gygi, M.P., Szpyt, J., Tam, S., Zarraga, G., Colby, G., Baltier, K., et al. (2015). The BioPlex Network: A Systematic Exploration of the Human Interactome. Cell 162, 425–440. 10.1016/j.cell.2015.06.043.

43. Gluderer, S., Brunner, E., Germann, M., Jovaisaite, V., Li, C., Rentsch, C.A., Hafen, E., and Stocker, H. (2010). Madm (Mlf1 adapter molecule) cooperates with Bunched A to promote growth in Drosophila. Journal of biology 9, 9–9. 10.1186/jbiol216.

44. Evans, R., O’Neill, M., Pritzel, A., Antropova, N., Senior, A., Green, T., Žídek, A., Bates, R., Blackwell, S., Yim, J., et al. (2022). Protein complex prediction with AlphaFold-Multimer. bioRxiv, 2021.2010.2004.463034. 10.1101/2021.10.04.463034.

45. Lim, R., Winteringham, L.N., Williams, J.H., McCulloch, R.K., Ingley, E., Tiao, J.Y.H., Lalonde, J.-P., Tsai, S., Tilbrook, P.A., Sun, Y., et al. (2002). MADM, a Novel Adaptor Protein That Mediates Phosphorylation of the 14-3-3 Binding Site of Myeloid Leukemia Factor 1. Journal of Biological Chemistry 277, 40997–41008. 10.1074/jbc.M206041200.

46. Go, C.D., Knight, J.D.R., Rajasekharan, A., Rathod, B., Hesketh, G.G., Abe, K.T., Youn, J.-Y., Samavarchi-Tehrani, P., Zhang, H., Zhu, L.Y., et al. (2021). A proximity-dependent biotinylation map of a human cell. Nature 595, 120–124. 10.1038/s41586-021-03592-2.

47. Go, C.D., Knight, J.D.R., Rajasekharan, A., Rathod, B., Hesketh, G.G., Abe, K.T., Youn, J.-Y., Samavarchi-Tehrani, P., Zhang, H., Zhu, L.Y., et al. (2022). Author Correction: A proximity-dependent biotinylation map of a human cell. Nature 602, E16–E16. 10.1038/s41586-021-04308-2.

48. Teo, G., Koh, H., Fermin, D., Lambert, J.-P., Knight, J.D.R., Gingras, A.-C., and Choi, H. (2016). SAINTq: Scoring protein-protein interactions in affinity purification - mass spectrometry experiments with fragment or peptide intensity data. Proteomics 16, 2238–2245. 10.1002/pmic.201500499.

49. Teo, G., Liu, G., Zhang, J., Nesvizhskii, A.I., Gingras, A.-C., and Choi, H. (2014). SAINTexpress: Improvements and additional features in Significance Analysis of INTeractome software. Journal of Proteomics 100, 37–43. 10.1016/j.jprot.2013.10.023.

50. Youn, J.-Y., Dyakov, B.J.A., Zhang, J., Knight, J.D.R., Vernon, R.M., Forman-Kay, J.D., and Gingras, A.-C. (2019). Properties of Stress Granule and P-Body Proteomes. Molecular cell 76, 286–294. 10.1016/j.molcel.2019.09.014.

51. Mittag, T., and Parker, R. (2018). Multiple Modes of Protein-Protein Interactions Promote RNP Granule Assembly. J Mol Biol 430, 4636–4649. 10.1016/j.jmb.2018.08.005.

52. Wang, J., Choi, J.M., Holehouse, A.S., Lee, H.O., Zhang, X., Jahnel, M., Maharana, S., Lemaitre, R., Pozniakovsky, A., Drechsel, D., et al. (2018). A Molecular Grammar Governing the Driving Forces for Phase Separation of Prion-like RNA Binding Proteins. Cell 174, 688–699 e616. 10.1016/j.cell.2018.06.006.

53. Zarin, T., Strome, B., Peng, G., Pritisanac, I., Forman-Kay, J.D., and Moses, A.M. (2021). Identifying molecular features that are associated with biological function of intrinsically disordered protein regions. Elife 10. 10.7554/eLife.60220.

54. Manning, G., Whyte, D.B., Martinez, R., Hunter, T., and Sudarsanam, S. (2002). The Protein Kinase Complement of the Human Genome. Science 298, 1912–1934. 10.1126/science.1075762.

55. Hernández-Plaza, A., Szklarczyk, D., Botas, J., Cantalapiedra, Carlos P., Giner-Lamia, J., Mende, D.R., Kirsch, R., Rattei, T., Letunic, I., Jensen, Lars J., et al. (2023). eggNOG 6.0: enabling comparative genomics across 12 535 organisms. Nucleic Acids Research 51, D389–D394. 10.1093/nar/gkac1022.

56. Li, Y., Calvo, S.E., Gutman, R., Liu, J.S., and Mootha, V.K. (2014). Expansion of biological pathways based on evolutionary inference. Cell 158, 213–225. 10.1016/j.cell.2014.05.034.

57. Altschul, S.F., Madden, T.L., Schaffer, A.A., Zhang, J., Zhang, Z., Miller, W., and Lipman, D.J. (1997). Gapped BLAST and PSI-BLAST: a new generation of protein database search programs. Nucleic Acids Res 25, 3389–3402. 10.1093/nar/25.17.3389.

58. Erdős, G., and Dosztányi, Z. (2020). Analyzing Protein Disorder with IUPred2A. Current protocols in bioinformatics 70, e99–e99. 10.1002/cpbi.99.

59. Mészáros, B., Erdos, G., and Dosztányi, Z. (2018). IUPred2A: context-dependent prediction of protein disorder as a function of redox state and protein binding. Nucleic acids research 46, W329–W337. 10.1093/nar/gky384.

60. Uversky, V.N. (2020). Analyzing IDPs in Interactomes. Methods in molecular biology (Clifton, N.J.) 2141, 895–945. 10.1007/978-1-0716-0524-0_46.

61. Lee, E.C.-H., and Strange, K. (2012). GCN-2 dependent inhibition of protein synthesis activates osmosensitive gene transcription via WNK and Ste20 kinase signaling. American journal of physiology. Cell physiology 303, C1269–1277. 10.1152/ajpcell.00294.2012.

62. Kester, H.A., Blanchetot, C., den Hertog, J., van der Saag, P.T., and van der Burg, B. (1999). Transforming growth factor-beta-stimulated clone-22 is a member of a family of leucine zipper proteins that can homo- and heterodimerize and has transcriptional repressor activity. J Biol Chem 274, 27439–27447. 10.1074/jbc.274.39.27439.

63. Pepin, A., Biola-Vidamment, A., Latre de Late, P., Espinasse, M.A., Godot, V., and Pallardy, M. (2015). [TSC-22D proteins: new regulators of cell homeostasis?]. Med Sci (Paris) 31, 75–83. 10.1051/medsci/20153101016.

64. Demir, S., Wolff, G., Wieder, A., Maida, A., Buhler, L., Brune, M., Hautzinger, O., Feuchtinger, A., Poth, T., Szendroedi, J., et al. (2022). TSC22D4 interacts with Akt1 to regulate glucose metabolism. Sci Adv 8, eabo5555. 10.1126/sciadv.abo5555.

65. Ekim Ustunel, B., Friedrich, K., Maida, A., Wang, X., Krones-Herzig, A., Seibert, O., Sommerfeld, A., Jones, A., Sijmonsma, T.P., Sticht, C., et al. (2016). Control of diabetic hyperglycaemia and insulin resistance through TSC22D4. Nat Commun 7, 13267. 10.1038/ncomms13267.

66. Jones, A., Friedrich, K., Rohm, M., Schafer, M., Algire, C., Kulozik, P., Seibert, O., Muller-Decker, K., Sijmonsma, T., Strzoda, D., et al. (2013). TSC22D4 is a molecular output of hepatic wasting metabolism. EMBO Mol Med 5, 294–308. 10.1002/emmm.201201869.

67. Liang, F., Li, Q., Li, X., Li, Z., Gong, Z., Deng, H., Xiang, B., Zhou, M., Li, X., Li, G., et al. (2016). TSC22D2 interacts with PKM2 and inhibits cell growth in colorectal cancer. Int J Oncol 49, 1046–1056. 10.3892/ijo.2016.3599.

68. Chong, P.A., and Forman-Kay, J.D. (2016). Liquid-liquid phase separation in cellular signaling systems. Curr Opin Struct Biol 41, 180–186. 10.1016/j.sbi.2016.08.001.

69. Khandwala, C.B., Sarkar, P., Schmidt, H.B., Ma, M., Kinnebrew, M., Pusapati, G.V., Patel, B.B., Tillo, D., Lebensohn, A.M., and Rohatgi, R. (2023). Direct ionic stress sensing and mitigation by the transcription factor NFAT5. bioRxiv. 10.1101/2023.09.23.559074.

70. Abu-Arish, A., Pandzic, E., Luo, Y., Sato, Y., Turner, M.J., Wiseman, P.W., and Hanrahan, J.W. (2022). Lipid-driven CFTR clustering is impaired in cystic fibrosis and restored by corrector drugs. J Cell Sci 135. 10.1242/jcs.259002.

71. Alberti, S., Gladfelter, A., and Mittag, T. (2019). Considerations and Challenges in Studying Liquid-Liquid Phase Separation and Biomolecular Condensates. Cell 176, 419–434. 10.1016/j.cell.2018.12.035.

72. Chen, J.-C., Lo, Y.-F., Lin, Y.-W., Lin, S.-H., Huang, C.-L., and Cheng, C.-J. (2019). WNK4 kinase is a physiological intracellular chloride sensor. Proceedings of the National Academy of Sciences of the United States of America 116, 4502–4507. 10.1073/pnas.1817220116.

73. Piala, A.T., Moon, T.M., Akella, R., He, H., Cobb, M.H., and Goldsmith, E.J. (2014). Chloride sensing by WNK1 involves inhibition of autophosphorylation. Science signaling 7, ra41–ra41. 10.1126/scisignal.2005050.

74. Saha, B., Leite-Dellova, D.C.A., Demko, J., Sørensen, M.V., Takagi, E., Gleason, C.E., Shabbir, W., and Pearce, D. (2022). WNK1 is a chloride-stimulated scaffold that regulates mTORC2 activity and ion transport. Journal of cell science 135. 10.1242/jcs.260313.

75. Vernon, R.M., Chong, P.A., Tsang, B., Kim, T.H., Bah, A., Farber, P., Lin, H., and Forman-Kay, J.D. (2018). Pi-Pi contacts are an overlooked protein feature relevant to phase separation. Elife 7. 10.7554/eLife.31486.

76. Yasukawa, T., Tsutsui, A., Tomomori-Sato, C., Sato, S., Saraf, A., Washburn, M.P., Florens, L., Terada, T., Shimizu, K., Conaway, R.C., et al. (2020). NRBP1-Containing CRL2/CRL4A Regulates Amyloid β Production by Targeting BRI2 and BRI3 for Degradation. Cell reports 30, 3478–3491.e3476. 10.1016/j.celrep.2020.02.059.

77. Morishita, K., Watanabe, K., Naguro, I., and Ichijo, H. (2023). Sodium ion influx regulates liquidity of biomolecular condensates in hyperosmotic stress response. Cell Rep, 112315. 10.1016/j.celrep.2023.112315.

78. Naguro, I., Umeda, T., Kobayashi, Y., Maruyama, J., Hattori, K., Shimizu, Y., Kataoka, K., Kim-Mitsuyama, S., Uchida, S., Vandewalle, A., et al. (2012). ASK3 responds to osmotic stress and regulates blood pressure by suppressing WNK1-SPAK/OSR1 signaling in the kidney. Nat Commun 3, 1285. 10.1038/ncomms2283.

79. Watanabe, K., Morishita, K., Zhou, X., Shiizaki, S., Uchiyama, Y., Koike, M., Naguro, I., and Ichijo, H. (2021). Cells recognize osmotic stress through liquid-liquid phase separation lubricated with poly(ADP-ribose). Nat Commun 12, 1353. 10.1038/s41467-021-21614-5.

