## Supplemental Methods for "*TSC22D*, *WNK* and *NRBP* gene families exhibit functional buffering and evolved with Metazoa for macromolecular crowd sensing"

**RESOURCE AVAILABILITY**

Materials availability: All unique reagents in this study are available from the Lead Contact and will be provided upon request.

Data and code availability: Data including additional supplemental videos and tables have been deposited at Mendeley and are publicly available as of the date of publication. DOI is listed in the key resources table and method section.

**METHODS DETAILS**

**Cell culture**

HAP1 cells (Horizon Discovery) were maintained in IMDM (Wisent). HEK293T (ATCC), HeLa-kyoto (Cellosaurus), Huh-7 (ATCC) and RAW264.7 (ATCC) cells were maintained in Dulbecco's Modified Eagle Medium (DMEM) (Wisent). UM-UC-4 (ECACC) cells were maintained in Eagle's Minimum Essential Medium (EMEM) (Wisent). BV-2 (ACCEGEN), U-937 (ATCC), THP-1 (ATCC) cells were maintained in Roswell Park Memorial Institute 1640 Medium (RPMI 1640) (Wisent). All cell lines were supplemented with 10% FBS (Gibco) and as needed supplemented with 1% penicillin–streptomycin (PS) (Gibco). All cell lines were maintained at 37°C with 5% CO2 and passaged regularly. Adherent cell lines (HEK293T, HeLa-kyoto, Huh-7, RAW264.7, UM-UC-4, and BV2 cells) were dissociated with 0.25% trypsin-EDTA (Gibco). All cell lines were regularly monitored for mycoplasma contamination using the e-Myco PLUS Mycoplasma PCR Detection kit (iNtRON Bio). See **KEY RESOURCE TABLE** for detailed cell line information.

**Gene co-dependency analysis**

Gene families arising through whole genome duplication were obtained from the OhnologsV2 database^1^ which comprised 1827 gene families using ‘strict’ criteria for generation of **Fig. 1A** and **S1A** (**Data S1**). Small-scale duplicates assigned to families in the database (i.e. paralogs) were not removed from the gene families before analysis. The “Chronos_Combined” dataset (version 22Q2) from the Cancer Dependency Map (DepMap) was accessed using the ‘depmap’ Bioconductor package (v1.12.0) and used to assess essentiality. Three sets of essential genes were created by applying different criteria/thresholds to the DepMap data. First, we extracted genes labelled as “Common Essential” by DepMap (excluding those that were also labelled as “Strongly Selective”). Second, we extracted genes that were essential in ≥ 60% of the DepMap screens (i.e. DepMap60). Finally, we extracted genes that were essential in ≥ 30% of the DepMap screens (i.e. DepMap 30). Each gene set was intersected with the ohnolog families, and the number of essential genes in each family was enumerated. Bootstrap distributions (n = 1000) were created by randomly sampling equivalent sized gene sets from the DepMap with no threshold criteria and assigning them to groups of the same size and number as the families in the given gene set. The 25th percentile of the permuted data was chosen as to represent the lower bound of random expectation. Expression data was downloaded from DepMap (OmicsExpressionProteinCodingGenesTPMLogp1.csv; release 22Q4), as were the gene perturbation cell viability scores (CRISPR_gene_effect.csv; release 22Q4) used for the co-essentiality analysis. The correlation matrix was created using the “cor” function in R, with the method parameter set to “pearson”. Hierarchical clustering was performed using the pheatmap function, with the clustering distance for both rows and columns set to 1- the Pearson correlation coefficient (**Data S1**).

For generation of **Fig. 1B and S1B** (**Data S1**), DepMap Public 22Q4 was analyzed to generate co-dependency profiles against all human genes for *TSC22D2*, *NRBP1* and *WNK1* knockouts in cancer cell lines (<https://depmap.org/>). For this, the Pearson correlation of knockout effects (i.e.: CERES scores) between the query gene and all other genes was calculated, and genes with Pearson correlation > 0.28 were considered “co-dependent”. The co-dependencies were plotted as interactions with Cytoscape v3.6.1^2^. Interactions between co-dependent genes were also added if they shared a Pearson correlation > 0.28. *TSC22D2*, *WNK1* and *NRBP1* expression and essentiality analyses were performed with ShinyDepmap ^3^ and visualized using GraphPad (PRISM software).

**Generation of CRISPR mutant clones**

HAP1 gene mutation cell lines for *TSC22D1*, *TSC22D2*, *NRBP1*, *WNK1*, *WNK2*, and *WNK3* were generated by electroporation of either [1] plasmid systems that express target gRNAs and spCas9 or [2] Cas9-ribonucleoprotein (RNP) complexes. For plasmid systems, target gRNAs were cloned into a modified version of the pSpCas9(BB)−2A-Puro (PX459v2) plasmid (Addgene #62988).

For Cas9-RNP systems, CRISPR guide RNA sequences (gRNAs) were synthesized by Integrated DNA Technologies (IDT), complexed at 1:1 molar ratio with tracrRNA (IDT) and then combined with Alt-R Cas9 enzyme (IDT) at 1:3 ratio to form RNP complexes. Parental HAP1 cells were electroporated using the Neon™ Transfection System with either PX459v2 plasmid or Cas9-RNP containing the target sgRNAs against the above genes using the following parameters: 1575V, 10ms and 3 pulses. For plasmid electroporated cells, puromycin selection was applied 24 h post-electroporation to select for cells expressing the plasmid. Cells were allowed to recover for 72hrs post-electroporation, then both plasmid and Cas9-RNP perturbed cells were seeded at average 0.5 cell/well for single-cell cloning in 96-well plates by seeding 0.5 cells per well. Following the expansion of single-cell clones, genomic DNA was extracted (Extracta™ DNA Prep for PCR, Quantabio), PCR amplified across the guide region, and sequenced by Sanger sequencing to identify clones with premature termination codons. Single knock-out clones were further validated by western blot to confirm the knockout of the protein. Sequential gene-knockout was done in selected single knockout clones for the generation of dual and triple gene mutation cell lines: *TSC22D2:TSC22D1, WNK1:WNK2:WNK3, TSC22D2:TSC22D1:TSC22D4.* See **KEY RESOURCE TABLE** for all guide sequences.

**Pooled CRISPR screens**

The Toronto knockout (TKOv3) genome-scale CRISPR library (Addgene #90294) was used to perform pooled CRISPR knockout screens in HAP1 parental and knockout cell lines to identify genetic interactions^4^. For each screen, a total of 9 x 10^7^ cells were transduced with TKOv3 lentivirus at an MOI of 0.3 and > 200-fold library coverage. 24 hours after infection, transduced cells were selected with a medium containing 1 μg/ml puromycin for 48 hours. Cells were then harvested, pooled, and split into three replicates of at least 1.5 x 10^7^ cells maintaining 200-fold library coverage. Approximately 3 – 5 x 10^7^ cells were also collected for gRNA extraction and determination of library representation at day 0 (T0 reference). Cells were passaged every 3-4 days at > 200-fold library coverage until 15 population doublings were reached. Cell pellets for gRNA extraction were collected at every cell passage for each replicate. For WNK463 drug-treated screens, replicates were separated into drug or vehicle (DMSO, <0.1% v/v) treatment screening arms at passage T4. Both WNK463-treated and DMSO control replicates were passaged at > 200-fold coverage on the same day until drug dosing at IC50 reached three rounds. Cell pellets for treated and control replicates were also collected at each passage.

gRNA abundance was measured at T0 and endpoints of each screen by next-generation sequencing (NGS). To prepare sequencing libraries, genomic DNA was extracted from cell pellets using the Wizard Genomic DNA Purification kit (Promega). Sequencing libraries were prepared from 50 µg gDNA (200-fold library coverage of haploid genome) via a 2-step nested PCR using primers that include Illumina TruSeq adapters with i5 and i7 indices. Barcoded libraries were gel purified using PureLink Quick Gel Extraction kit (Thermo Fisher Scientific) and sequenced on an Illumina HiSeq2500 or NovaSeq6000 using single-read sequencing and standard primers for dual-indexing with HiSeq SBS Kit v4 or NovaSeq v1.5 reagents. T0 samples were sequenced at read depths of 500-fold library coverage, while all other samples were sequenced at 200-fold. Amplicons were sequenced using a custom sequencing recipe comprising DC: 21 cycles; R1: 36 cycles; Index read 1: 8 cycles; Index read 2: 8 cycles.

All screens were analyzed using methods described previously for HAP1 drop-out screens and for WNK463 drug screens^5,6^. Briefly, reads from each sample were mapped to the TKOv3 reference library using Bowtie (v0.12.8) allowing up to two mismatches and one exact alignment (specific parameters: -v2, -m1, -p4 -sam-nohead). Successfully aligned reads were counted and combined into a matrix with guide annotations. Read counts were normalized to 10 million reads per sample and log fold-changes (LFC) of each gRNA between each timepoint and T0 were determined. Guides with fewer than 30 or greater than 10,000 absolute read counts at T0 were removed. To identify genetic interactions, the qGI algorithm^5^ was used to score the differential effects between HAP1 parental and gene knockout screens, while the Bioconductor package limma^6^ was used to examine drug-treated versus DMSO effects. Interactor genes with FDR (or minimum FDR across replication screens) > 0.2 were plotted with GraphPad (PRISM software).

**Cell size measurements**

To prepare cells for cell volume measurements, cells were seeded at low densities (e.g., 5 x 10^4^ cells on one well of 6-well plates) on day0 to minimize contact inhibition. After overnight incubation, cells were quickly washed with PBS and dissociated using 0.25% trypsin-EDTA (Gibco) for 2 minutes followed by resuspension in 500 µL of media. Cell volumes were measured using the Multisizer 4e Coulter Counter (Beckman-Coulter) to determine the diameter and distribution of average cell volume under different conditions. For drug-induced cell volume changes, cells were seeded at low density on day 0 and allowed to settle overnight. Fresh medium containing the drug or DMSO control was replaced on day 1 and incubated for 24 hrs. For adherent cell types, the cells were quickly washed once with PBS at room temperature, trypsinized for 2 min and resuspended with 500 µL culture medium prior to size measurement.

**Intracellular ion concentration quantification**

For intracellular [Na^+^] and [K^+^] measurements, HAP1 cells were grown and treated in 60 mm tissue culture plates at confluency. Cells were washed three times with an osmotically balanced solution of cold 150 mM LiCl (300 mOsm) on ice and lysed with 1 ml cold 1% HNO_3_ lysis buffer. [Na^+^] and [K^+^] were measured using a PinAAcle 900F atomic absorption spectrometer (PerkinElmer), and [Na^+^] and [K^+^] were then normalized to cell number and volume.

**Inhibitor studies**

The effect of WNK463 on cell growth and proliferation was determined by AlamarBlue™ (ThermoFisher Scientific) staining according to the manufacturer’s instructions. Briefly, 4 x 10^3^ cells were seeded in a 96-well plate and treated with WNK463 serially diluted three-fold in DMSO to a final concentration of <0.5% DMSO. Treated cells were incubated at 37°C with 5% CO_2_ for three days. After three days, the culture medium was removed and incubated with 10% AlamarBlue™ solution for one hour. Following incubation, AlamarBlue™ fluorescence was quantified at 560nm excitation and 590nm emission wavelength using a Varioskan™ LUX multimode microplate reader (ThermoFisher Scientific). Half-maximal inhibitory concentration (IC50) was determined by converting relative fluorescence units (RFU) into percentages relative to the levels of untreated controls (defined as 100%) and fitted by non-linear regression using GraphPad (PRISM software) for macOS.

**Endogenous protein tagging**

### CRISPR-Cas9-mediated homology-directed repair was used to fluorescently tag endogenous proteins. Donor repair plasmids and guide RNA designs used for cell line construction are described in detail in STAR protocol and KEY RESOURCE TABLE. Briefly, donor repair plasmids containing the desired fluorescent tag were flanked by 400-1000 bp homology arms targeting either the C- or N-terminus of the protein of interest. Gene fragments containing desired homology arms were synthesized by TWIST Biosciences and cloned into pUC19 by digesting with EcoRI and HindIII and cloning insertion fragments (both homology arms and PCR products containing designated fluorescent protein or miniTurbo) by Gibson assembly. Cas9 guides were cloned into pX459v2 as described above. Target cells were transfected with both Cas9-targeting and donor repair plasmids at a molar ratio of 3:1 using X-tremeGene 9™ (Roche) transfection reagent or the Neon™ Transfection System (ThermoFisher Scientific) using the same parameters described above. Cells were kept in culture for 48-72 hours, then subjected to puromycin (1µg/ml), zeocin (200µg/ml), or blasticidin (10µg/ml) selection to isolate cells expressing the target protein fused to the fluorescent tag. Genomic DNA was extracted, and PCR amplified in homology arm regions to confirm the proper insertion of tags at the C- or N-terminus of the protein by Sanger sequencing. Clones with correct endogenous tagged gene sequences were then examined by confocal imaging as described below.

**Live cell imaging and analysis**

For confocal live-cell imaging, endogenously tagged HAP1 cells were seeded in Perkin Elmer LLC CellCarrier-96 Ultra Microplates (PerkinElmer) 18 hours prior to imaging. Cells were then imaged at indicated treatment and time points on the Opera Phenix High-Content Screening System (PerkinElmer). Images were processed and analyzed with Columbus and Harmony (PerkinElmer) software for spot number quantifications. Intensity fluctuation measurements were performed using Fiji-ImageJ.

For TIRF live-cell imaging, endogenously tagged HAP1 cells were seeded in 8-well chamber slides (Nunc® Lab-Tek® II chambered cover glass). Normal culture medium for live imaging experiments consisted of Phenol-red free IMDM medium (Gibico) supplemented with 10% FBS and 1% PS. Stress medium included addition of 700 mOsmol/L sorbitol. Media was changed prior to imaging using 3D-printed adaptors and pumps designed for medium change. Stress medium was changed between frames 2 and 3, and normal phenol-red free IMDM medium changed between frames 26-27. Highly inclined laminated optical sheet (HILO) imaging was performed on a modular imaging system built around an IX-83 base (Olympus, Canada) using a 100X oil-immersion TIRF objective (UApoN 100×/1.49 Oil, Olympus, Japan). mScarlet was excited with a 561nm laser (GEM 200 mW, Laser Quantum, UK) with the laser power modulated to 5 mW at the objective. Images were captured using a sCMOS camera (Prime BSI, Photometrics, USA) and, at 100x magnification, resulted in an effective pixel size of 65 nm x 65 nm. A detailed snapshot of the optical system is described in the following file (<https://github.com/YipLab/IX83-Modules/releases/tag/1.2>). Each experiment consisted of 50 frames and was imaged at 10 seconds per frame, resulting in a total acquisition time of 500 seconds. Acquired movies were then deconvolved and analyzed with Fiji-ImageJ for foci number quantifications. Macro scripts from ImageJ were written for image processing and foci number quantification and are available upon request. Overlaid supplemental movie is generated by ImageJ, Keynote, and iMovie.

**Immunofluorescence**

Cells were seeded and exposed to stress conditions in black 96-well PerkinElmer LLC CellCarrier Ultra Microplates (PerkinElmer). After treatments, cells were fixed with 4% paraformaldehyde solution in PBS for 15 minutes at room temperature. After fixation, cells were washed with PBS and permeabilized with 0.1% Triton-X 100 solution in PBS for 10 minutes. Cells were then washed three times with PBS and blocked for one hour at room temperature in buffer containing 0.3% Triton-X 100, 5% FBS in PBS. Primary antibodies were diluted in buffer containing 0.3% Triton-X 100, 1% BSA in PBS and incubated with cells for overnight staining at 4°C. The next day, cells were washed three times with PBS and stained with matching secondary antibodies for one hour at room temperature. Cells were then washed with PBS and incubated with 1 µg/ml DAPI as needed prior to imaging. Antibody and staining reagent specifications are listed in **KEY RESOURCE TABLE**. Cells were imaged on the Opera Phenix QHES (PerkinElmer) automated high-content screening system. Intensity fluctuation at indicated regions was performed in Fiji-ImageJ.

**Protein-protein interaction domain prediction with AF2**

Amino acid sequences of the canonical isoforms of human NRBP1, NRBP2, TSC22D1, TSC22D2, and TSC22D4 was used to generate fasta files to input into AlphaFold2-Multimer (v2.X)^7,8^. AlphaFold2-Multimer was run using the multimer model with full database size and including all databases needed to model multimers, with a maximum template date of January 14th 2023, and running 5 seeds per model to a total of 25 predictions per protein pair (Sample script). A total of 2 GPUs and 8 CPUs were used per prediction, running times and memory usage were determined empirically for each dataset. Computations were performed on the Graham supercomputer at the SciNet HPC Consortium^9,10^. Models were evaluated based on the predicted aligned error (PAE) matrix, extracted from the .pkl file of each run, and by visualizing the colored 3D structures based on the pLDDT scores using the .pdb files. PAE matrices were generated for all 25 predictions using python and 3D structures were visualized with Protein Viewer^11^.

**Proximity-based labelling of proteins captured with mass spectrometry.**

BioID coupled to mass spectrometry was similarly performed essentially as previously described^12-15^. Briefly, HAP1 cell lines were endogenously tagged with 2xHA-miniTurbo on the N-terminus of *TSC22D2*, *WNK1* or *NRBP1* as described above. HA1 parental cells and HAP1 parental transduced with overexpressed miniTurbo are included as background controls. Successfully tagged cell lines were seeded on 150mm tissue culture plates (1 x 10^7^ cells) and incubated overnight. The next day cells were treated with 460 mM sorbitol and/or 50 μΜ biotin for one hour, or 1 μM WNK463 inhibitor and/or 50 μΜ biotin for 30 min to label proximal proteins under stress and normal conditions. Cells were then washed twice on ice with cold PBS and directly lysed in plates with modified RIPA lysis buffer [50 mM Tris-HCl pH 7.4, 150 mM NaCl, 0.5mM EDTA, 1mM 150 mM MgCl2, 0.1% (wt/vol) SDS, 1% NP-40, 0.5 mM EGTA, 0.4% sodium deoxycholate, and Halt™ Protease Inhibitor Cocktail (100X) (Thermo Fisher Scientific)]. Cells were sonicated for 30 seconds (5 seconds on, 5 seconds off for three cycles) at high amplitude using a 4°C water bath sonicator (Bioruptor® Plus). Subsequently, 1 μl Benzonase nuclease (Thermo Fisher Scientific) was added to each sample and the mixture nutated at 4°C for 30 minutes. 10% SDS was added to bring each sample’s final SDS concentration to 0.5%, and samples were further rotated at 4°C for 15 minutes and centrifuged at 15,000g (Eppendorf centrifuge 5424R) for 20 min at 4°C to clarify the lysate. Total proteins were normalized to 3mg/ml using a BCA assay (Pierce™ BCA Protein Assay Kit). 1.2 ml of normalized supernatant was added to pre-washed Streptavidin Sepharose beads (GE Healthcare 17-5113-01) and rotated at 4°C for 3 hours. Beads were washed once with 2% SDS wash buffer (2% SDS, 50 mM Tris-HCL pH 7.5), twice in RIPA wash buffer, once in TNNE wash buffer (25 mM Tris-HCl, pH 7.4, 150 mM NaCl, 0.1% NP40, 1 mM EDTA) and three times in 50 mM ammonium bicarbonate pH 8.0 (ABC) wash buffer. Following purification, 1 μg (in 50 μl ABC) of sequencing-grade trypsin (Sigma-Aldrich) was added and incubated overnight at 37°C with agitation. Subsequently, another 0.5 μg trypsin was added, and the sample was incubated for three more hours to ensure complete digestion. Peptides were collected into a new tube, the beads were washed with MS-grade H2O, and the washes were combined with the peptide fraction. Peptides in solution were acidified with formic acid (5% final concentration). Samples were then lyophilized and stored at -80°C until analysis by mass spectrometry.

**Mass spectrometry**

LC-MS/MS analysis was conducted on a TripleTOF 6600 (SCIEX) mass spectrometer with a nanoelectrospray ion source coupled in-line to a 425 Nano-HPLC (Eksigent). A fused silica column (15 cm x ID 100 μm, OD 360 μm) with integrated emitter tip was prepared and packed in-house using a laser puller (Sutter Instruments Co.) and ~15 cm of C18 (3 μm) resin (Reprosil-Pur). Samples were resuspended in 10 μl 5% formic acid with 4 μl of sample used per injection for data-dependent acquisition. Samples were loaded onto the column via an autosampler and an organic phase gradient was delivered over 90 min at 400 nl/min (2-35% acetonitrile with 0.1% formic acid). The mass spectrometer was operated in data-dependent acquisition mode with 1 MS scan (250 ms; mass range 400-1250 m/z) followed by up to 20 MS/MS scans (50 ms each). Ions were dynamically excluded for 10 s with a 50 mDa window and only candidate ions between two and five charge states were considered. Each sample was run with 2 blank samples (5% formic acid, 3 rapid gradient cycles at 1500 nl/min over 30 min) and 1 BSA quality control sample run over 30 min prior to the next sample. A 30 min BSA mass calibration run was also performed before each sample.

Mass spectrometry data reported in this paper is available on MassIVE (http://massive.ucsd.edu) at the following accession numbers: MSV000092162; MSV000092163. Data has been deposited as a complete submission to the MassIVE repository (https://massive.ucsd.edu/ProteoSAFe/static/massive.jsp) and assigned the accession numbers MSV000092162 and MSV000092163. The datasets are currently available for reviewers at ftp://MSV000092162@massive.ucsd.edu and ftp://MSV000092163@massive.ucsd.edu. Please login with username MSV000092162_reviewer; password: TWNBODYWNK1 for dataset 1 (VS2) and username MSV000092163_reviewer; password: TWNBODYWNK2 for dataset 2 (VS3). The dataset will be made public upon acceptance of the manuscript.

**Peptide and protein identification**

MS raw files (.WIFF and .WIFF.SCAN) were converted to MGF and mzML formats using ProteoWizard (v3.0.4468) and the AB SCIEX MS Data Converter (V1.3 beta) through ProHits^16^. The search database contained forward and reversed human and adenovirus sequences (RefSeq protein database v57) supplemented with “common contaminants” from the Max Planck Institut (<http://lotus1.gwdg.de/mpg/mmbc/maxquant_input.nsf/7994124a4298328fc125748d0048fee2/$FILE/contaminants.fasta>) and the Global Proteome Machine as well as common epitope tags was used. MS spectra were analyzed separately using Mascot (2.3.02 Matrix Science) and Comet^17^(2012.01 rev.3) , with trypsin specificity and up to two missed cleavages specified. Deamidation (NQ) and oxidation (M) were selected as variable modifications, with no fixed modifications selected. Fragment mass tolerance was set to 0.15 Da, and a mass window of ±40 ppm for the precursor with charges of 2+ to 4+ (monoisotopic) was selected. Comet and Mascot results were processed by PeptideProphet90 and combined into an iProphet^18^ and combined into an iProphet^19^ output using the Trans-Proteomic Pipeline (TPP; Linux version, v0.0 Development trunk rev 0, Build 201303061711). General TPP options were –p0.05, -x20, -PPM, -dDECOY

**SAINT analysis of high-confidence proximity partners**

The probability that an identified protein (prey) was significantly enriched in a given bait profile over a series of negative control runs was calculated using SAINTexpress (v3.6.1)^15^. SAINTexpress uses a semi-supervised spectral counting model to determine this probability and only proteins with an iProphet protein probability >0.95 and a minimum of two unique peptides were considered. Baits and controls were processed in biological triplicates. The negative controls consisted of streptavidin purifications from lysates generated from HAP1 cells without a bait-miniTurbo fusion (which models endogenous biotinylation) as well as each bait-miniTurbo fusion cell line with no biotin treatment (which models the biotinylation noise of each bait). 42 independent controls were used separately without compression. The three biological replicates were compressed into two, and SAINTexpress scores were averaged across these duplicates to calculate a Bayesian False Discovery Rate (BFDR). Unless otherwise specified, a stringent BFDR of 1% or less is considered high confidence.

**Reproducibility metrics**

Calculation of R values was based on the high-confidence proximal associations (BFDR<0.05) detected with either of the baits compared (e.g., with or without stress) and was performed in EXCEL using PEARSON function (not normalized or transformed) extracted from the SAINTexpress file^14,15^ . Prey ACACA is excluded from PCC analysis because ACACA has overly high spectral counts and is often detected as background noise. Details for assessments are available in **Data S3**.

**Bait-prey proximity interaction visualization**

Dot plots and heatmap in **Fig. 3H**, and **S7D** are generated by ProHits-viz^20^. SAINTexpress^14,15^ analyzed output files that were used as input. Preys with more than two average spectral counts (AvgSpec) from three technical replicates were displayed with the BFDR primary filter of 0.01 and the secondary filter of 0.05. AvgSpec displays were capped at 50. The bait-prey network in **Fig. S3F** was produced using Cytoscape v3.6.1^2^.

**Conditional proximity protein component correlation analysis**

SAINTexpress output files were filtered with a loose BFDR<0.25 cutoff to be conservative ^15^. AvgSpec for each prey captured by the same bait but under different conditions was inputted to calculate the Pearson correlation performed by GraphPad (PRISM software). minBFDR between the two conditions in comparison was used to colour each bait dot. Prey ACACA is excluded from Pearson correlation analysis because ACACA has overly high spectral counts and is often detected as background noise.

**Evolutionary signature analysis with protein intrinsic disordered regions**

For data preparation, the core set (128 preys) are defined in **Fig. S5A** and **Data S2**. Protein sequences of interest were retrieved from UniProt using ID mapping from UniProt/Gene Name to UniProtKB in Homo sapiens [9606]. Reviewed (Swiss-Prot) sequences were adopted. Sequences were then manually examined for missing or duplicated sequences, re-examined with alternative names, and adopted canonical variants. For the “HAP1 expressed” set, the corresponding genes were filtered based on RNA expression with counts per million (cpm)>1 in HAP1 cells.

To define intrinsic disorder regions, SPOT-Disorder predictor v1.0^21^ was used to predict the probability of intrinsic disorder on a residue level for every protein sequence in the human proteome. A disorder probability above 0.5 was used to define disordered residues. Only the protein regions with 30 or more consecutively predicted intrinsically disordered residues were considered as IDRs in all subsequent analyses.

Evolutionary signature identification and clustering of the matrix of Z-scores was done as previously described ^22^. The maxtrix of Z-scores representing evolutionary signatures of intrinsically disordered regions of the core set was extracted from that of the total human proteome and represented as a heatmap and visualized using Treeview^23^. GO enrichment analyses were performed with PANTHER^24^ using Homo sapiens as background and annotated manually.

**Coevolution analysis**

For data preparation, the evolutionary histories of TSC22D, WNK, NRBP family proteins as well as OXSR1 and SPAK were predetermined by CLIME software^25^. WNK1 homologues were then identified by BLASTP search against each target proteome, followed by domain architecture analysis and manual inspection in species of interest. Hits were selected with an Expect value < 0.001, retaining only the top homolog. Putative WNK homologs were accepted as valid if: a) BLASTP of object protein sequence against the human proteome returned WNK paralogues as the top result; b) contains a kinase domain that is similar to human WNK paralog kinase domain; c) the sequence is intact with methionine as the start; d) more similar in sequence percentile identity to human WNK paralogs than to human NRBP paralogs (**Fig. S5C**). The sequences were cross-referenced against the EggNOG v6 ^26^databases. PCA analysis for selected protein sequences was performed with JalView using the BLOSUM62 substitution matrix and percentile identity as input; the first two principal components were plotted with GraphPad (PRISM software).

For the determination of WNK paralog intrinsic disorder fraction, IUPRED2a scores^27-29^ were used for predicting intrinsically disordered residues, and a cut-off of >0.5 was applied. Intrinsic disorder fractions were then calculated as the number of disordered residues divided by the total protein length.

For phylogenic tree construction, WNK full-length kinase domain sequences from selected organisms were used to perform multiple sequence alignment with MUSCLE^30^. The phylogenetic tree was then generated using the default settings. Tree visualization was performed by ChiPlot^31^ (https://www.chiplot.online) and manually labelled with taxonomic branch names for clarity.

**RNAi knockdown of worms and saline sensitivity assay**

Wild-type C. elegans (N2 Bristol) used in these studies were maintained at 20°C. RNA interference (RNAi) was performed by feeding a library of bacteria expressing double stranded RNA (dsRNA; Source Bioscience) to worms. RNAi bacteria were streaked onto LB media plates (with ampicillin and tetracycline) and incubated for 18 hours at 37°C. Individual colonies were selected with a sterile p200 tip and were transferred to tubes containing liquid LB + ampicillin + tetracycline. These were grown for 24 hours (at 37°C) in an orbital shaker, prior to induction of dsRNA synthesis with isopropyl-β-D-thiogalactoside (0.1 M) for 4 hours (37°C, orbital shaker). Next, the bacteria were plated on RNAi media plates (containing isopropyl-β-D-thiogalactoside and carbenicillin). The worms were bleach synchronized at the L1 stage and then grown for two generations on RNAi plates before F1 young adults were transferred to either control RNAi or high salinity RNAi plates as previously described^32^. Survival was assessed every 24 hours using responsiveness to touch and pharyngeal pumping as viability indicators. Survival plot and Gehan-Breslow-Wilcoxon tests are generated by GraphPad (PRISM software).

**Co-Immunoprecipitation**

For co-IP between endogenous NRBP1 and TSC22D2 (**Fig. 3A**), HAP1 cells expressing ^mNeon-2xHA^TSC22D2 were utilized. Cells were cultured 48 hours prior to one hour in D-sorbitol containing medium. Cells were lysed using IP lysis buffer (Thermo Fisher Scientific) supplemented with combined EDTA-free Halt Protease and Phosphatase Inhibitor Cocktail (Pierce). Lysates were pre-cleared for non-specific binding with Protein A/G PLUS agarose (Thermo Fisher Scientific). 500 µg of pre-cleared protein was then incubated with an anti-HA antibody (1:100) overnight at 4°C to bind with endogenous mNeon-2xHATSC22D2 or an anti-GST antibody (1:100) that was used as a non-specific binding control. Protein-antibody immunocomplexes were further enriched with Protein A/G PLUS agarose. Collected proteins were washed three times with IP lysis buffer and then eluted from beads using sample buffer mixture containing 5:3:2 ratio of NuPAGE LDS sample buffer (4X):IP lysis buffer:NuPAGE sample reducing agent (10X) and incubating for 5 min at 95°C. See **KEY RESOURCE TABLE** for detailed antibody specifications.

For co-IP between exogenous TSC22D2 and NRBP1 truncations with endogenous WNK1, TSC22D2 and NRBP1 (**Fig. 3D**), 293T cells were transfected with CMV promoter-driven overexpression plasmids containing exogenous protein truncations tethered with either C-terminus or N-terminus FLAG, including TSC22D2-Nterm (1-257), TSC22D2-Cterm (258-780), NRBP1-Cterm (1-422), NRBP1-Nterm (423-535) for 48hrs before harvesting with IP lysis buffer (Thermo Fisher Scientific) supplemented with combined EDTA-free Halt Protease and Phosphatase Inhibitor Cocktail (Pierce). 500 µg of protein was then incubated with an anti-M2 antibody (1:100) overnight at 4°C to bind with overexpression constructs tethered with FLAG. 293T lysates containing transfection of CMV-mClover construct with absence of FLAG tag was used as a mock control. Collected proteins were washed three times with IP lysis buffer and then eluted from beads using sample buffer mixture containing 5:3:2 ratio of NuPAGE LDS sample buffer (4X):IP lysis buffer:NuPAGE sample reducing agent (10X) and incubating for 5 min at 95°C. See **KEY RESOURCE TABLE** for detailed antibody specifications.

**Western Blotting**

Cells were lysed in RIPA buffer containing protease and phosphatase inhibitors (Pierce) for 10 minutes on ice. Next, 10–30 µg protein was resolved on 4–12% Bis‐Tris gels or 3-8% Tris-Acetate gels (Life Technologies) and transferred to Immobilon‐P nitrocellulose membrane (Millipore) at 66 V for 90 minutes. Subsequently, proteins were detected using primary antibodies and HRP-conjugated secondary antibodies, and proteins were visualized on iBright™ imaging system using SuperSignal West Pico Enhanced chemiluminescence (ECL) substrates (Thermo Fisher Scientific) and Western Lightning Pro ECL substrates (PerkinElmer). See **KEY RESOURCE TABLE** for detailed antibody specifications.

**Statistical analysis**

The number of technical and/or biological replicates is listed in the figure legends or text for all experiments. Indicated statistical analyses were performed using GraphPad (PRISM Software) or the R language programming environment.

**KEY RESOURCE TABLE**

| REAGENT or RESOURCE | SOURCE | IDENTIFIER |
| --- | --- | --- |
| Antibodies and cell stains | | |
| Streptavidin, Alexa Fluor™ 488 conjugate | Invitrogen | S11223 |
| HA-Tag (6E2) | Cell Signaling | 2367S |
| GST Antibody (B-14) | Santa Cruz Biotechnology | sc-138 |
| TSC22D2 | Novus | NBP1-81491 |
| WNK1 | Abcam | ab137687 |
| NRBP1 | Origene | TA500437 |
| OSR1 | Atlas | HPA008237 |
| ^pS373^SPAK/^pS325^OSR1 | Millipore Sigma | 07-2273 |
| mCherry | Thermo Fisher Scientific | PA5-34974 |
| G3BP1 | BD Transduction Labs | AB_398437 |
| EEA1 | Cell Signaling | cs2411 |
| GOLGA2 | Sigma-Aldrich | HPA021799 |
| Clathrin-HC | Cell Signaling | cs4796S |
| RAB11 | Cell Signalling | cs5589 |
| TOMM20 | Cell Signaling | cs42406 |
| SQSTM1 | Abcam | ab207305 |
| LAMP1 | Cell Signaling | cs9091 |
| RAB7 | Cell Signalling | cs9367 |
| CLINT1 | Abcam | ab251892 |
| FASN | Abcam | ab128870 |
| PRRC2C | Sigma-Aldrich | HPA025766 |
| DCP1A | Thermo Fisher Scientific | A303-590A |
| Streptavidin Agarose beads | GE Healthcare Life Science | 17511301 |
| Protein A/G PLUS agarose | Thermo Fisher Scientific | 20423 |
| Anti-DYKDDDDK Magnetic Agarose | Pierce | A36797 |
| Goat anti-Mouse IgG (H+L) Cross-Adsorbed Secondary Antibody, Alexa Fluor™ | Thermo Fisher Scientific |  |
| Goat anti-Rabbit IgG (H+L) Cross-Adsorbed Secondary Antibody, Alexa Fluor™ | Thermo Fisher Scientific |  |
| Alexa Fluor™ 647 Phalloidin | Invitrogen | A22287 |
| Goat anti-Rabbit IgG (H+L) Secondary Antibody, HRP | Invitrogen | RRID: AB_228341 |
| Goat anti-Mouse IgG (H+L) Secondary Antibody, HRP | Invitrogen | RRID: AB_228307 |
| DAPI (4',6-Diamidino-2-Phenylindole, Dihydrochloride) | Invitrogen | D1306 |
| ER-Tracker™ Blue-White DPX, for live-cell imaging | Invitrogen | E12353 |
| Bacterial and virus strains | | |
| ElectroMAX™ Stbl4™ Competent Cells | Invitrogen | 11635018 |
| One Shot™ Stbl3™ Chemically Competent *E. coli* | Invitrogen | C737303 |
| Chemicals, peptides, and recombinant proteins | | |
| T4 DNA Ligase | NEB | M0202S |
| T4 PNK | NEB | M0201S |
| 2X MyTaq™ Red Mix | Bioline | BIO-25043 |
| 2 X KAPA Taq ReadyMix | Sigma-Aldrich | KK1006 |
| WNK463 | Selleckchem | S8358 |
| WNK-IN-11 | Selleckchem | S0015 |
| rapamycin | Selleckchem | S1039 |
| Torin2 | Selleckchem | S2817 |
| 1,6-Hexanediol | Millipore Sigma | 240117 |
| 1,2,6-Hexanetriol | Millipore Sigma | T66206 |
| SB202190 | Selleckchem | S1077 |
| SP600125 | Selleckchem | S1460 |
| TAK715 | Selleckchem | S2928 |
| calyculin A | Millipore Sigma | 208851 |
| okadaic acid | Millipore Sigma | 459620 |
| ZT-1a | MedChemExpress | HY-136532 |
| PTC-028 | Selleckchem | S8662 |
| 5-FU | Selleckchem | S1209 |
| tunicamycin | Selleckchem | S7894 |
| sorafenib | Selleckchem | S7397 |
| cerulenin | MedChemExpress | HY-A0210 |
| Leptomycin B solution from Streptomyces sp. | Millipore Sigma | L2913-.5UG |
| Nigericin | Selleckchem | NSC 292567 |
| Sodium arsenite solution | Millipore Signma | 1.06277 |
| bumetanide | Selleckchem | S1287 |
| Hydrochlorothiazide | Selleckchem | S1708 |
| amiloride | Selleckchem | No.S1811 |
| cycloheximide | Millipore Signma | 01810-1G |
| hipporestanol | MedKoo | 561328 |
| rSAP | NEB | M0371S |
| EcoRI-HF | NEB | R3101S |
| HindIII-HF | NEB | R3104S |
| BbsI | NEB | R0539S |
| Critical commercial assays | | |
| GeneArt Gibson Assembly® Cloning | Invitrogen | [A46624](https://www.thermofisher.com/order/catalog/product/A46624) |
| Wizard® HMW DNA Extraction Kit | Promega | A2920 |
| Qubit dsDNA BR Assay Kit, 500 assays | Invitrogen | Q32853 |
| BCA protein assay | Thermo Fisher Scientific | 23225 |
| SuperSignal West Pico Enhanced chemiluminescence (ECL) substrates | Thermo Fisher Scientific | 34580 |
| 4–12% Bis‐Tris gels | Invitrogen | NP0322BOX |
| NuPAGE 3-8% Tris-Acetate gels | Invitrogen | EA03785BOX |
| NuPAGE™ MOPS SDS Running Buffer | Invitrogen | NP0001 |
| NuPAGE Tris-Acetate SDS running buffer | Invitrogen | LA0041 |
| BCA Protein Assay Kit | Thermo Fisher Scientific Chemicals | J63283.QA |
| PureLink™ HiPure Plasmid Maxiprep Kit | Invitrogen | K210006 |
| PureLink™ Quick Gel Extraction Kit | Invitrogen | K210012 |
| PureLink™ Quick Plasmid Miniprep Kit | Invitrogen | K210010 |
| Extracta DNA Prep for PCR | Quantabio | 95091-002 |
| MassIVE | [https://massive.ucsd.edu/ProteoSAFe/static/massive.jsp](https://can01.safelinks.protection.outlook.com/?url=https%3A%2F%2Fmassive.ucsd.edu%2FProteoSAFe%2Fstatic%2Fmassive.jsp&data=05%7C01%7Cjason.moffat%40sickkids.ca%7C8043db71968540b36ce708db6d23d2ea%7C3961553ff47e49eb9f6ccf8518914e9a%7C0%7C0%7C638223772848565673%7CUnknown%7CTWFpbGZsb3d8eyJWIjoiMC4wLjAwMDAiLCJQIjoiV2luMzIiLCJBTiI6Ik1haWwiLCJXVCI6Mn0%3D%7C3000%7C%7C%7C&sdata=ZfY1BWSt5R2UaSP2seNLAh0pFvOhfdZBOJiYPFGOWjA%3D&reserved=0) | MSV000092162; MSV000092163 |
| Experimental models: Cell lines |  |  |
| Human: HAP1 | Horizon | RRID: CVCL_Y019 |
| Human: HeLa (Kyoto) | Cellosaurus | RRID: CVCL_1922 |
| Human: THP-1 | ATCC | Cat#TIB-202; RRID: CVCL_0006 |
| Human: UM-UC-4 | ECACC | RRID: CVCL_2749 |
| Human: HEK293T | ATCC | RRID: CRL-3216 |
| Human: HUH-7 | ATCC | RRID: CVCL_0336 |
| Mouse: RAW264.7 | ATCC | Cat#91062702-1VL, RRID: CVCL_0493 |
| Mouse: BV-2 | ACCEGE | Cat#ABC-TC212S; RRID: CVCL_0182 |
| Human: U-937 | ATCC | Cat#CRL-1593.2; RRID: CVCL_0007 |
| Experimental models: Organisms/strains | | |
| *C. elegans* (N2 Bristol) |  |  |
| Oligonucleotides | | |
| WNK3-N-LHA | TTGTAAAACGACGGCCAGTGAATTCccactgaattgtataattaaaaatggttaaaatggtaaatttgacgatatgtatattttagtacagtaaaagaAACTACTATTGATGAATTCCAGAATTTTAGACAAAAACCACCTAGAAACCATGTGGTTGTTTTACTTACTGTGTTTGTCACATTGTGGCTAAGTGAACTCAGGGTTTGGGTACTCATTTTGAATTTGATGGATTGACAGATGGAGCTGAGAACTAGGTTTCTAACCAATATTCATGCAATATGAAGACAGATTGAGctatatactaagtagtctgctagacactgaggactaatggtaaacaagataagacacagtccctAGAGTCTGCTGGTTCCTAAGATTGGACAGAAAAAAGAATAAACTTCTTATTAAGGAAAAAGGGATTAAAAAATCAGGTGATTTGTTTTCTCCTTAATGATTTCCTTTGTAGGGTGATTTTAAAATATCTTTTTCCCCCCCTTCAGGTTGCTATGGAAATACATGACCACGCAAAAGGAAGTCCATTCTGATAATTCTGATACCTGAGATGTAACTGGACTGAAGAGTAGAAACAGGAAAAATTTTAGTGCCAACTTTAATTACAATGGGCAAGAGCAAGACCGGTAGC | |
| WNK3-N-RHA | AGGAAGCGGAGGATCTGGCGGTACCGCCACTGATTCAGGGGATCCAGCCAGCACAGAAGATTCTGAGAAACCTGATGGAATTTCATTTGAAAACAGAGTTCCCCAGGTCGCTGCAACTTTGACAGTAGAAGCTAGACTAAAGGAGAAAAACAGTACCTTCTCTGCTTCTGGGGAAACTGTAGAAAGGAAGAGATTTTTCCGAAAGAGTGTTGAAATGACGGAAGATGACAAAGTTGCCGAATCATCCCCCAAAGATGAGAGAATTAAGGCTGCAATGAATATTCCAAGAGTAGATAAGCTTCCTTCAAATGTGTTGAGAGGTGGACAAGAAGTTAAATATGAACAGTGTTCAAAGTCAACCTCAGAAATCTCAAAAGATTGTTTCAAGGAGAAAAATGAAAAGGAAATGGAAGAAGAAGCAGAAATGAAGGCTGTAGCTACTTCTCCTAGTGGCAGATTCCTGAAATTTGACATAGAACTAGGAAGAGGAGCATTTAAAACAGTATATAAAGGACTGGACACTGAAACATGGGTTGAGGTTGCTTGGTGTGAGCTGCAGGTAGGTATATAATCTTCTTGGTTATAATAAATTCAAGTTTTAGCTGGCCTGGCCTTCTGTGTGATTCATAAGCTTGGCGTAATCATGGTCATAG | |
| TSC22D1-N-LHA | TTGTAAAACGACGGCCAGTGAATTCCCGGGGGTGGCGAGGGGAAGGCGACATTTGCTTGGCGCTGCCTCCGTGCAGAGCGGCCGGGAGGGCTTTGCAGCGGCCGCCGACGCGGCGGGAGGAGGGGGCTGAGGCGGCTTCGGAGTTGGCCGGAGAAGGTGGGCATTTCTCGCTTTTCCTCCCCTTCCTGCGCTCTCCCCCTCCCTCCTGCCCCGCACCCCACGTGAAGCAGAATATAAAGGGGGGTGTGAGGCTAGAGGGGAAAGTGAATGGCGAAGGACTGAAGGGATCCCCCCTTCGGGTCCCCGGCCGCCCTGTTCACCCTCGTTCATCCTCCTTTCCGAAGCTCGCTCTCGAAGGCAGGAGCGACCGGCGCCTTTGGCTGAGGAGGAGGAGAAGGAGGAATCGCGCCAGGCGGAGCGTCAGGTCCCGTTTTCCTCTCCGGCGTCTCCAATACAAAGATTACGGTGCAGAAGGAAATTGCACTCGTCTCCTCCGCGCCCCCGGTACCCAACACAATGGGCAAGAGCAAGACCGGTAGC | |
| TSC22D1-N-RHA | AGGAAGCGGAGGATCTGGCGGTACCCACCAGCCGCCTGAGTCCACCGCCGCGGCCGCCGCCGCTGCAGACATTAGCGCTAGGAAGATGGCGCACCCGGCAATGTTCCCTCGAAGGGGCAGCGGTAGTGGCAGCGCCTCTGCTCTCAATGCAGCAGGTACCGGCGTCGGTAGTAATGCCACATCTTCCGAGGATTTTCCGCCTCCGTCGCTGCTTCAGCCGCCGCCCCCTGCAGCATCTTCTACGTCGGGACCACAGCCTCCGCCTCCACAAAGCCTGAACCTCCTTTCGCAGGCTCAGCTGCAGGCACAGCCTCTTGCGCCAGGCGGAACTCAAATGAAAAAGAAAAGTGGCTTCCAGATAACTAGCGTTACTCCTGCTCAGATCTCCGCTAGTATCAGCTCTAACAACAGTATAGCAGAGGACACTGAGAGCTATGATGATCTGGATGAATCTCACACGGAAGATCTCTCTTCTTCGGAGATCCTTGATGTGTCACTTTCCAGGGCTACTGACTTAGGGGAGCCCGAACGCAGCTCCTCAGAAGAGACCCTAAATAACTTCCAGGAAGCCGAGACACCTGGGGCAGTCTCTCCCAACCAGCCCCACCTTCCTCAGCCTCATTTGCCAAGCTTGGCGTAATCATGGTCATAG | |
| TSC22D4-N-LHA | TTGTAAAACGACGGCCAGTGAATTCctctctctctctctctctcCCCCTCTTTTTCCAGTTTGCAAACTCAGCTCTGGAGTTCAGCAGCAACAGCAGCAGGAAAAACCTGcccctgctcccccctcccgccacctcccctctcctcttctcccctcacccaGCAGGCACCCCCGGTTCCCGCCAGGCCCTCCTGCCATGTCGGACCCAGACGTCCCCAGGGGCTCGGATGTCCCCGCCATGTGGCCCCCTTGTTCCAGGGGTGCCTGAGCCCCTTCAAGGAGCCCCAACCCACCCCCAACCTTGGCCCAGCCCTGAGCCCCAGGGACCATGGGCAAGAGCAAGACCGGTAGC | |
| TSC22D4-N-RHA | AGGAAGCGGAGGATCTGGCGGTACCAGCGGGGGCAAGAAGAAGAGTAGTTTTCAAATCACCAGCGTCACTACGGACTATGAGGGCCCTGGGAGCCCAGGGGCTTCGGATCCCCCTAccccacagcccccaaccgggcccccgccccgcctgcccaatggggagcccagccccgaTCCGGGGGGCAAGGGCACCCCCCGGAATGGCTCCCCACCACCTGGGGCCCCTTCCTCCCGTTTCCGGGTGGTGAAGCTGCCCCACGGCCTGGGAGAGCCTTATCGCCGCGGTCGCTGGACGTGTGTGGATGTTTATGAGCGAGACCTGGAGCCCCACAGCTTCGGCGGACTCCTGGAGGGAATTCGAGGGGCCTCAGGGGGCGCCGGGGGCAGATCTTTGGATTCCAGGTTGGAGCTGGCCAGCCTCGGCCTGGGCGCCCCCACCCCACCGTCAGGCCTGTCTCAGGGCCCCACCTCCTGGCTCCGTCCACCCCCCACCTCTCCTGGACCTCAGGCCCGCTCCTTCACTGGGGGACTGGGCCAGCTGGTGGTGCCCAGCAAAGCCAAGGCAGAGAAACCCCCACTGTCGGCCTCCTCACCCCAGCAGCGCCCCCCAGAGCCTGAGACCGGTGAGAGTGCGGGCACATCCCGGGCTGCCACGCCCCTGCCCTCTCTGAGGGTGGAAGCGGAGGCTGGGGGCTCAGGGGCCAGGACCCCTCCACTGTCCCGGAGGAAAGCTGTAGACATGCGGCTGCGGATGGAGTTGGGTGCTCCAGAAGAGATGGGGCAGGTAAGACCTGGGTTCTAGGGCTGGCCCATCAGCCCTGGCCTAACCTCTATCTTGAGcctccttcctcctcctcccccccctctcctcctccttctccATCACTTTATAAGATCAGATAGCCCTCTTCTCTGTGCCTGCCAGGTCACTCTGAAGCCCCTTCCTCCTACTTCATTTTCTGTCTCAGCCTCCTTCCTGTTTTTCTGCACCCTCTCTCTTTAAGCTTGGCGTAATCATGGTCATAG | |
| WNK1-N-LHA | TTGTAAAACGACGGCCAGTGAATTCGAGCCGGGCCGCGGCCTTCCCTCGCCCGCCTCGGCCCCTCCCACTCCTCTGCCCCGGGGCCGCCACCGCCCGGGCGTCGGACCTGGTCCCGTGCTCGCGGTGCCGCCGCCCTCTGGGCCTAGCCCGCCCAGCTCGGCGAGCGGCGGCAGTGGGAGCCGCGTCCGCCGCATCCGCCTCGACTCGGTGCCGGCCCCTGGCCCTCCCCTCATGACTGCGGCGCCTCTGCTGCCACCGCCCGCCCGGCCGCCGCTCGCCGCAGGATGGATGCGGACCGTGCGGCGCTAACCCCCGTGGCTCAGCTCCCGAATCGCCCGCCTTCGAGCCCTCCTCGTGAGCCGCAGCAGCCTCGGTGCCAGCCCCCGCCGCAGCTGGGCCCAGCGGTCCGCCTGTCCCTCGTTGCGGCTTGTCGGTGCTGAGTGAGGCGTCGTCCGGGTCGGCGCGAACCCGCCCGGCCGCGGTTCCCTGCAGACCTCTGCGCGGGCGGCTCGGCCCTTCACGCCCTTTTCGTTCACGAATCCGAGCCCGCTCGCCTCTCTCCAGCGAACCGACCATGGGCAAGAGCAAGACCGGTAGC | |
| WNK1-N-RHA | AGGAAGCGGAGGATCTGGCGGTACCTCTGGCGGCGCCGCAGAGAAGCAGAGCAGCACTCCCGGTTCCCTGTTCCTCTCGCCGCCGGCTCCTGCCCCCAAGAATGGCTCCAGCTCCGATTCCTCCGTGGGGGAGAAACTGGGAGCCGCGGCCGCCGACGCTGTGACCGGCAGGACCGAGGAGTACAGGCGCCGCCGCCACACTATGGACAAGGACAGCCGTGGGGCGGCCGCGACCACTACCACCACTGAGCACCGCTTCTTCCGCCGGAGCGTCATCTGTGACTCCAATGCCACTGCACTGGAGCTTCCCGGCCTTCCTCTTTCCCTGCCCCAGCCCAGCATCCCCGCGGCTGTCCCGCAGAGTGCTCCACCGGAGCCCCACCGGGAAGAGACCGTGACCGCCACCGCCACTTCCCAGGTAGCCCAGCAGCCTCCAGCCGCTGCCGCCCCTGGGGAACAGGCCGTCGCGGGCCCTGCCCCCTCGACTGTCCCCAGCAGTACCAGCAAAGACCGCCCAGTGTCCCAGCCTAGCCTTGTGGGGAGCAAAGAGGAGCCGCCGCCGGCGAGAAGTGGCAAGCTTGGCGTAATCATGGTCATAG | |
| TSC22D2-N-LHA | TTGTAAAACGACGGCCAGTGAATTCAGCGCCGAGCTGGACTGACCACGGCTGCCCGGAGACGAGAGAGGAAGCAGCCGGCCCGCCCCTGGGTTCGCGCTCTCGCCGCCTCTGAGGGAATTGAATTGAGGCGCCGCGGCTGCGAGAGCTAAAAAGGAAGGAGGAGCCGCCGCGGGACTGAGACGGGGGCAGAGCCGAAGAGACCGACACAGAGAAGGAAACGAGGAGGAGGATGTCTCACCGGGCGGCCAGCGCCTGGATCAGCCCGTGACTCTTAACAGCGGCGGGCCTCAGACCCCAGCGCAGACTCGGACTTTGTCTTTGGGGGCCCGTGCTCTGCCCTCCCCGGTTTCCGACAGGACCCAGAGGAGCCGGCGTGCCTCTCTGCCCTCCAGCCTTCTTCACCATGGGCAAGAGCAAGACCGGTAGC | |
| TSC22D2-N-RHA | AGGAAGCGGAGGATCTGGCGGTACCTCCAAGATGCCGGCCAAGAAGAAGAGCTGCTTCCAGATCACCAGTGTCACCACGGCCCAGGTGGCCACTAGCATCACCGAGGACACCGAGAGCTTGGACGACCCGGACGAGTCACGCACAGAGGACGTCTCCTCCGAGATTTTCGACGTCTCTCGGGCCACGGATTATGGCCCTGAGGAGGTCTGCGAGCGCAGCTCTTCCGAAGAGACGCTTAACAATGTTGGGGATGCGGAGACTCCCGGGACCGTCTCCCCAAACCTCCTCCTAGATGGGCAGCTGGCAGCGGCGGCTGCTGCTCCCGCCAACGGAGGAGGAGTCGTTTCGGCCCGGAGCGTGTCTGGGGCGCTCGCCAGTACCCTGGCGGCGGCTGCCACTTCGGCCCCCGCCCCCGGAGCACCCGGCGGCCCCCAGCTCGCGGGCTCATCCGCCGGGCCAGTGACTGCAGCCCCATCTCAGCCTCCCACCACATGTAGTTCCCGTTTTCGCGTGATCAAGCTGGACCACGGGAGCGGAGAGCCCTATAGACGCGGCCGATGGACGTGTATGGAATACTATGAGAGGGATTCAGACAGCAGCGTCCTGACTAGATCCGGGGATTGCATTAGACACAGCAGTACTTTTGACCAGACTGCGGAGCGGGACAGCGGCCTGGGCGCCACCGGAGGGTCGGTGGTGGTAGTAGTGGCCTCCATGCAAAGCTTGGCGTAATCATGGTCATAG | |
| NRBP1-N-LHA | TTGTAAAACGACGGCCAGTGAATTCtcactgcagcctcaaactcctgggcaatcaagtgattctcccatcagcctctcgggtagctgggactatagctgtgtaccaccttgccctgctaatttttaaatttttttgtaaagatggggtctcactttgttgcccaggctggtctcaaacacctggcctcaaatgaacctcttgcctcggcttctcaaagtgctgggattacagaggtgagccactgtgctcagcTGGATTACTAAAtttttttttgagatggggtcttgctctgtcacccaggctggagtgcagtggcgtgatctcagctcactgcaacctccacctccctggttcaaaaaaattctcctgccttagtctcctgagtagctaggactacaggcacgcgtcaccacacccagctaattttgtatttttagtagagatggggtttcaccatgttgatcaggctggtcttgaactcctgacctcaggtgatctgcccacctcagcctcccaaagtgctgggattacaggcatgagccaccacgtccagccTGGATTACTAAATCTTTACAGATGTATCCTGACTTTCTCTGCCTTTTCCCTCTGTATTTGCCCCAACCAGTGCAGGCCTGAGTGTTCCTTCCAGCATGGGCAAGAGCAAGACCGGTAGC | |
| NRBP2-N-RHA | AGGAAGCGGAGGATCTGGCGGTACCTCGGAGGGGGAGTCCCAGACAGTACTTAGCAGTGGCTCAGACCCAAAGGTAGAATCCTCATCTTCAGCTCCTGGCCTGACATCAGTGTCACCTCCTGTGACCTCCACAACCTCAGCTGCTTCCCCAGAGGAAGAAGAAGAAAGTGAAGATGAGTCTGAGATTTTGGAAGAGTCGCCCTGTGGGCGCTGGCAGAAGAGGCGAGAAGAGGTAAGGTTATGGTACAGTTACTCTTGGGTGAGTGAATTCTGGAAAGGTGAGAACTGGAAGAGCAAAGTCTTAAAAGAGGCCAACCAAACTTCAATAGGTTGGGGGCTAGGAGGAGATGATCATATGGAGTGTTAAAATGGAGGATTTGTGAGATAAGTTTAATTCTCTTTCATATGAATTTCTTCTCAGGTGAATCAACGGAATGTACCAGGTATTGACAGTGCATACCTGGCCATGGATACAGAGGAAGGTGTAGAGGTTGTGTGGAATGAGGTACAGTTCTCTGAACGCAAGAACTACAAGCTGCAGGAGGTAGGTGATGCTGAAAAGGTGAAGCCTGGGGAATGTTGGAACACTGATGTTAGAAAGCTTGGCGTAATCATGGTCATAG | |
| G3BP1-N-LHA | TTGTAAAACGACGGCCAGTGAATTCGTACTTTGCATAAATATGTACAGTACCATCTTTTCACACTGGTGATGATGAGGTTTTATTCTAAGCCTCCAAATCTATGGAGAAGGTGGCAAttttttttattttttattttttattgggacggagtctcactctgtcgcccatgctggagtgcagtggcacgaccttggctcaccacaacttccgccacccgggttcaagTtgcactccagtctgggtgacgagtgaaactctgtctcattaaaaaaacaaacaaaaaaCTGAACTGTGAGAACTTAGCTGATTTATAATCTGCTGGttctttGGtttttctgagtcggagtctcgctctttttcccagactggagtacaatggcgtgatcttggctcactgcaacctccgcctcctgggttcaagcaattctcctgcctcagcctccttgaatagctggaactacaggcgtgtgtcaccatgtctggctaatttttgtgtttttttagtagagacggggtttcaccatgttggtcaggctggtcttgaactcctgacctcgtaatccacccgtctcggcctcccaaagtgctgggattacaggcatgagtcaccgtgcccagccTCTGCTGGTTCATTATTACAGCTTTCTTTATCTTTGATAGTGATAGCCAGCTCATCAGTACTCTTAAGTCTGGTCACCTTGATTCTTACAGCCCATCATGTTCAGAGGTGAGGTCCGTCTGAATGTCGAAGAGAAGAAGACTCGAGCTGCCAGGGAAGGCGACCGACGAGATAATCGCCTTCGGGGACCTGGAGGCCCTCGAGGTGGGCTGGGTGGTGGAATGAGAGGCCCTCCCCGTGGAGGCATGGTGCAGAAACCAGGATTTGGAGTGGGAAGGGGGCTTGCGCCACGGCAGGGAGGAAGCGGAGGATCTGGCGGTA | |
| G3BP1-N-RHA | ATGGGCGGCCGCTAAGAGCAAGAAGTGAATCTTCATGGATCTTCATGCAGCCATACAAACCCTGGTTCCAACAGAATGGTGAATTTTCGACAGCCTTTGGTATCTTGGAGTATGACCCCAGTCTGTTATAAACTGCTTAAGTTTGTATAATTTTACTTTTTTTGTGTGTTAATGGTGTGTGCTCCCTCTCCCTCTCTTCCCTTTCCTGACCTTTAGTCTTTCACTTCCAATTTTGTGGAATGATATTTTAGGAATAACGGACTTTTAAAGAAGCaaaaaaaaaGACTGAATTTCCTTGCTTACTTTGCATATACAGACTGGAttttttCtttttttttACAGCCATTTCCCCAAAGGAATGTCTTGCATATTACTGACATTTGGTATGTTTCATTCATTGGAATATTTCTTATTTTCTACGTGTTTGAAAAGCCTGTAAGAAATACAGGATTTGATAATATTTTGAAGGCAGGAAAAACCCAAATTGTTTCTTCTTTGAGAGTCATGACTACCTTCTGGTGTGGAGAAATTGCCATTGGAAAATTTGACAATTTTGATTCTCACTGGTATGTTTAAAAACTGAATAAAAGGAATAGAAtttttttttGATAAAGGATCACAAAACAATTCTAAAACCTAACTGTTTTTACCATTGAAATTTAAATTGTGATAATAGGTTTTAAATGTCTAGAATGCAACTGATAGGCTTTTCTTGAACTGTTAGTTTTTTTGAAGTAGTTTTTTCATGTTTAATTTGTATTTGTaaaaaaacaaaaagcaaaaaaaTTCCCAAAACCCAGATAACAACCAGAGCAAAACTGTTGTGCCTTCTATTTATCTTTGATTTCAGTCTTGGCAATTGTTTaaaaCaaaaaTCTAGATTTGTTTTATTAGGTTCAGAGTATGTGGGGAATTATAGAATCCCTCTTTCATCACTTTGTGTATGTCTTTTGTTAACATATTTGTTATGCCTTATTCTAAAATTGAGTCTCAAACTGGAATGCCTTTGAAGACAGATGCTTCTAAAGCTTGGCGTAATCATGGTCATAG | |
| *WNK1 mutant* sgRNA sequence | ATTCTACAGGCACAGTCCCA | |
| *WNK2 mutant* sgRNA sequence | CCTCAAGTTCGACATCGAGC | |
| *WNK3 mutant* sgRNA sequence | GCTGTAGCTACTTCTCCTAG | |
| *TSC22D1* mutant sgRNA sequence | CCGGCAATGTTCCCTCGAAG | |
| *TSC22D2 mutant* sgRNA sequence | ACGGCGGCCACCCTTCCCGT | |
| *NRBP1 mutant* sgRNA sequence | GTGAATCAACGGAATGTACC | |
| *NRBP2 mutant* sgRNA sequence | GGACGAGAGCGACATCCTGG | |
| *WNK1 mutant* synthego guide ID | ENSG00000060237 | |
| *TSC22D4 mutant* synthego guide ID | ENSG00000166925 | |
| *WNK1* endo-tag sgRNA sequence | AGCGAACCGACCATGTCTGG | |
| *TSC22D2* endo-tag sgRNA sequence | CTTGGCCGGCATCTTGGACA | |
| *NRBP1* endo-tag sgRNA sequence | TGTTCCTTCCAGCATGTCGG | |
| *WNK3* endo-tag sgRNA sequence | GTGGCCATTGTAATTAAAGT | |
| *TSC22D1* endo-tag sgRNA sequence | TGGTGCATTGTGTTGGGTAC | |
| *TSC22D4* endo-tag sgRNA sequence | CCGTGGTGACGCTGGTGATT | |
| *G3BP1* endo-tag sgRNA sequence | TCCATGAAGATTCACTGCCG | |
| Recombinant DNA | | |
| Mod-pSpCas9-2A-Puro (PX459) V2.0 | In-house | |
| pUC19 | Invitrogen | |
| Software and algorithms | | |
| Benchling | Benchling.com | |
| Fiji | https://imagej.net/software/fiji/ | |
| Jalview | https://www.jalview.org | |
| Treeview | https://bitbucket.org/TreeView3Dev/treeview3/src/master/ | |
| PRISM 9 | https://www.graphpad.com/ | |
| Chiplot | https://www.chiplot.online | |
| ShinyDepmap | https://labsyspharm.shinyapps.io/depmap/ | |
| ProHits-Viz | https://prohits-viz.org | |
| IUPred2A | https://iupred2a.elte.hu | |
| PANTHER | http://pantherdb.org | |
| Harmony High-Content Imaging and Analysis Software | PerkinElmer | |
| Columbus™ Image Data Storage and Analysis system | PerkinElmer | |
| Other | | |
| Puromycin Dihydrochloride | Gibco | A1113803 |
| Zeocin™ Selection Reagent | Gibco | R25005 |
| Blasticidin S HCl (10 mg/mL) | Gibco | A1113803 |
| D-Biotin | Thermo Fisheer Scientific Chemicals | R25005 |
| D-Sorbitol | Sigma-Aldrich | A1113903 |
| Sodium chloride | Sigma-Aldrich | 230090050 |
| Potassium chloride | Sigma-Aldrich | S1876 |
| Urea | Sigma-Aldrich | S9888 |
| PBS | WISENT | P9541 |
| FBS | Gibco | U5378 |
| IMDM, no phenol red | Gibco | 311-010-CL |
| Modified iscove’s medium | WISENT | A4766801 |
| DMEM medium | WISENT | 21056023 |
| EMEM medium | WISENT | 319-105-XK |
| RPMI medium | WISENT | 219-010-XK |
| RIPA Lysis and Extraction Buffer | Thermo Fisher Scientific | 320-005-CL |
| IP lysis buffer | Thermo Fisher Scientific | 350-000-EL |
| Triton-X 100 | MilliporeSigma | 89901 |
| Lithium chloride | Sigma-Aldrich | 87787 |
| HNO_3_ | CALEDON | 9036-19-5 |
| PerkinElmer LLC CellCarrier-96 Ultra Microplates | PerkinElmer | L9650 |
| Nunc® Lab-Tek® II chambered cover glass | Thermo Fisher Scientific | 7525-8-60 |
| Ammonium Bicarbonate | Bioshop | 6055308 |
| Monoclonal anti-FLAG M2 antibody | Sigma-Aldrich | 171080 |
| Halt™ Protease Inhibitor Cocktail (100X) | Thermo Fisher Scientific | 05-402-7 |
| Halt™ Protease and Phosphatase Inhibitor Cocktail (100X) | Thermo Fisher Scientific | F3165 |
| RNase | Bio Basic | 78430 |
| Trypsin | Sigma-Aldrich | 78440 |
| Turbonuclease | BioVision Inc. | RB0473 |

**SUPPLEMENTAL REFERENCES**

1.	Singh, P.P., and Isambert, H. (2020). OHNOLOGS v2: a comprehensive resource for the genes retained from whole genome duplication in vertebrates. Nucleic Acids Res *48*, D724-D730. 10.1093/nar/gkz909.

2.	Shannon, P., Markiel, A., Ozier, O., Baliga, N.S., Wang, J.T., Ramage, D., Amin, N., Schwikowski, B., and Ideker, T. (2003). Cytoscape: a software environment for integrated models of biomolecular interaction networks. Genome research *13*, 2498-2504. 10.1101/gr.1239303.

3.	Shimada, K., Bachman, J.A., Muhlich, J.L., and Mitchison, T.J. (2021). shinyDepMap, a tool to identify targetable cancer genes and their functional connections from Cancer Dependency Map data. eLife *10*. 10.7554/eLife.57116.

4.	Hart, T., Tong, A.H.Y., Chan, K., Van Leeuwen, J., Seetharaman, A., Aregger, M., Chandrashekhar, M., Hustedt, N., Seth, S., Noonan, A., et al. (2017). Evaluation and design of genome-wide CRISPR/SpCas9 knockout screens. G3: Genes, Genomes, Genetics *7*. 10.1534/g3.117.041277.

5.	Aregger, M., Lawson, K.A., Billmann, M., Costanzo, M., Tong, A.H.Y., Chan, K., Rahman, M., Brown, K.R., Ross, C., Usaj, M., et al. (2020). Systematic mapping of genetic interactions for de novo fatty acid synthesis identifies C12orf49 as a regulator of lipid metabolism. Nature Metabolism *2*. 10.1038/s42255-020-0211-z.

6.	Chan, K., Farias, A.G., Lee, H., Guvenc, F., Mero, P., Brown, K.R., Ward, H., Billmann, M., Aulakh, K., Astori, A., et al. (2023). Survival-based CRISPR genetic screens across a panel of permissive cell lines identify common and cell-specific SARS-CoV-2 host factors. Heliyon *9*, e12744-e12744. 10.1016/j.heliyon.2022.e12744.

7.	Evans, R., O’Neill, M., Pritzel, A., Antropova, N., Senior, A., Green, T., Žídek, A., Bates, R., Blackwell, S., Yim, J., et al. (2022). Protein complex prediction with AlphaFold-Multimer. bioRxiv, 2021.2010.2004.463034. 10.1101/2021.10.04.463034.

8.	Jumper, J., Evans, R., Pritzel, A., Green, T., Figurnov, M., Ronneberger, O., Tunyasuvunakool, K., Bates, R., Zidek, A., Potapenko, A., et al. (2021). Highly accurate protein structure prediction with AlphaFold. Nature *596*, 583-589. 10.1038/s41586-021-03819-2.

9.	Loken, C., Gruner, D., Groer, L., Peltier, R., Bunn, N., Craig, M., Henriques, T., Dempsey, J., Yu, C.-H., Chen, J., et al. (2010). SciNet: Lessons Learned from Building a Power-efficient Top-20 System and Data Centre. Journal of Physics: Conference Series *256*, 012026. 10.1088/1742-6596/256/1/012026.

10.	Ponce, M., Zon, R.v., Northrup, S., Gruner, D., Chen, J., Ertinaz, F., Fedoseev, A., Groer, L., Mao, F., Mundim, B.C., et al. (2019). Deploying a Top-100 Supercomputer for Large Parallel Workloads: the Niagara Supercomputer. Proceedings of the Practice and Experience in Advanced Research Computing on Rise of the Machines (learning). Association for Computing Machinery.

11.	Sehnal, D., Bittrich, S., Deshpande, M., Svobodova, R., Berka, K., Bazgier, V., Velankar, S., Burley, S.K., Koca, J., and Rose, A.S. (2021). Mol* Viewer: modern web app for 3D visualization and analysis of large biomolecular structures. Nucleic Acids Res *49*, W431-W437. 10.1093/nar/gkab314.

12.	Go, C.D., Knight, J.D.R., Rajasekharan, A., Rathod, B., Hesketh, G.G., Abe, K.T., Youn, J.-Y., Samavarchi-Tehrani, P., Zhang, H., Zhu, L.Y., et al. (2021). A proximity-dependent biotinylation map of a human cell. Nature *595*, 120-124. 10.1038/s41586-021-03592-2.

13.	Go, C.D., Knight, J.D.R., Rajasekharan, A., Rathod, B., Hesketh, G.G., Abe, K.T., Youn, J.-Y., Samavarchi-Tehrani, P., Zhang, H., Zhu, L.Y., et al. (2022). Author Correction: A proximity-dependent biotinylation map of a human cell. Nature *602*, E16-E16. 10.1038/s41586-021-04308-2.

14.	Teo, G., Koh, H., Fermin, D., Lambert, J.-P., Knight, J.D.R., Gingras, A.-C., and Choi, H. (2016). SAINTq: Scoring protein-protein interactions in affinity purification - mass spectrometry experiments with fragment or peptide intensity data. Proteomics *16*, 2238-2245. 10.1002/pmic.201500499.

15.	Teo, G., Liu, G., Zhang, J., Nesvizhskii, A.I., Gingras, A.-C., and Choi, H. (2014). SAINTexpress: Improvements and additional features in Significance Analysis of INTeractome software. Journal of Proteomics *100*, 37-43. 10.1016/j.jprot.2013.10.023.

16.	Liu, G., Zhang, J., Larsen, B., Stark, C., Breitkreutz, A., Lin, Z.-Y., Breitkreutz, B.-J., Ding, Y., Colwill, K., Pasculescu, A., et al. (2010). ProHits: integrated software for mass spectrometry-based interaction proteomics. Nature biotechnology *28*, 1015-1017. 10.1038/nbt1010-1015.

17.	Eng, J.K., Jahan, T.A., and Hoopmann, M.R. (2013). Comet: an open-source MS/MS sequence database search tool. Proteomics *13*, 22-24. 10.1002/pmic.201200439.

18.	Keller, A., Nesvizhskii, A.I., Kolker, E., and Aebersold, R. (2002). Empirical statistical model to estimate the accuracy of peptide identifications made by MS/MS and database search. Anal Chem *74*, 5383-5392. 10.1021/ac025747h.

19.	Shteynberg, D., Deutsch, E.W., Lam, H., Eng, J.K., Sun, Z., Tasman, N., Mendoza, L., Moritz, R.L., Aebersold, R., and Nesvizhskii, A.I. (2011). iProphet: multi-level integrative analysis of shotgun proteomic data improves peptide and protein identification rates and error estimates. Mol Cell Proteomics *10*, M111 007690. 10.1074/mcp.M111.007690.

20.	Knight, J.D.R., Choi, H., Gupta, G.D., Pelletier, L., Raught, B., Nesvizhskii, A.I., and Gingras, A.-C. (2017). ProHits-viz: a suite of web tools for visualizing interaction proteomics data. Nature methods *14*, 645-646. 10.1038/nmeth.4330.

21.	Hanson, J., Yang, Y., Paliwal, K., and Zhou, Y. (2017). Improving protein disorder prediction by deep bidirectional long short-term memory recurrent neural networks. Bioinformatics *33*, 685-692. 10.1093/bioinformatics/btw678.

22.	Lu, A.X., Lu, A.X., Pritišanac, I., Zarin, T., Forman-Kay, J.D., and Moses, A.M. (2022). Discovering molecular features of intrinsically disordered regions by using evolution for contrastive learning. PLOS Computational Biology *18*, e1010238-e1010238. 10.1371/journal.pcbi.1010238.

23.	Saldanha, A.J. (2004). Java Treeview—extensible visualization of microarray data. Bioinformatics *20*, 3246-3248. 10.1093/bioinformatics/bth349.

24.	Thomas, P.D., Ebert, D., Muruganujan, A., Mushayahama, T., Albou, L.P., and Mi, H. (2022). PANTHER: Making genome-scale phylogenetics accessible to all. Protein Sci *31*, 8-22. 10.1002/pro.4218.

25.	Li, Y., Calvo, S.E., Gutman, R., Liu, J.S., and Mootha, V.K. (2014). Expansion of biological pathways based on evolutionary inference. Cell *158*, 213-225. 10.1016/j.cell.2014.05.034.

26.	Hernández-Plaza, A., Szklarczyk, D., Botas, J., Cantalapiedra, Carlos P., Giner-Lamia, J., Mende, D.R., Kirsch, R., Rattei, T., Letunic, I., Jensen, Lars J., et al. (2023). eggNOG 6.0: enabling comparative genomics across 12 535 organisms. Nucleic Acids Research *51*, D389-D394. 10.1093/nar/gkac1022.

27.	Erdős, G., and Dosztányi, Z. (2020). Analyzing Protein Disorder with IUPred2A. Current protocols in bioinformatics *70*, e99-e99. 10.1002/cpbi.99.

28.	Mészáros, B., Erdos, G., and Dosztányi, Z. (2018). IUPred2A: context-dependent prediction of protein disorder as a function of redox state and protein binding. Nucleic acids research *46*, W329-W337. 10.1093/nar/gky384.

29.	Uversky, V.N. (2020). Analyzing IDPs in Interactomes. Methods in molecular biology (Clifton, N.J.) *2141*, 895-945. 10.1007/978-1-0716-0524-0_46.

30.	Edgar, R.C. (2004). MUSCLE: multiple sequence alignment with high accuracy and high throughput. Nucleic Acids Research *32*, 1792-1797. 10.1093/nar/gkh340.

31.	Xie, J., Chen, Y., Cai, G., Cai, R., Hu, Z., and Wang, H. (2023). Tree Visualization By One Table (tvBOT): a web application for visualizing, modifying and annotating phylogenetic trees. Nucleic Acids Res. 10.1093/nar/gkad359.

32.	Lant, B., Pal, S., Chapman, E.M., Yu, B., Witvliet, D., Choi, S., Zhao, L., Albiges-Rizo, C., Faurobert, E., and Derry, W.B. (2018). Interrogating the ccm-3 Gene Network. Cell Rep *24*, 2857-2868 e2854. 10.1016/j.celrep.2018.08.039.

33.	Billmann, M., Ward, H.N., Aregger, M., Costanzo, M., Andrews, B.J., Boone, C., Moffat, J., and Myers, C.L. (2023). Reproducibility metrics for context-specific CRISPR screens. Cell Syst *14*, 418-422 e412. 10.1016/j.cels.2023.04.003.

34.	Zagórska, A., Pozo-Guisado, E., Boudeau, J., Vitari, A.C., Rafiqi, F.H., Thastrup, J., Deak, M., Campbell, D.G., Morrice, N.A., Prescott, A.R., and Alessi, D.R. (2007). Regulation of activity and localization of the WNK1 protein kinase by hyperosmotic stress. The Journal of cell biology *176*, 89-100. 10.1083/jcb.200605093.

**SUPPLEMENTAL FIGURE LEGENDS**

**Figure S1.** *TSC22D2*, *WNK1* and *NRBP1* are functionally dominant paralogs in human cells, related to Figure 1. **A.** Bar plot showing the percentage of functionally dominant ohnolog gene families as a function of essentiality in the DepMap CRISPR data, where background gene essentiality is indicated by the dotted bar. Pie charts indicate the percentage of functionally dominant genes that are the highest expressed in each class where an average of 9.5% (max=11%, min=7%) of multigene families include a single essential gene. **B.** ShinyDepMap shows *TSC22D2, WNK1,* and *NRBP1* have the highest absolute efficacy in their sensitive cell lines among their paralogues^3^. X-axis represents efficacy (i.e., the degree to which perturbation of the gene reduces cell growth in sensitive lines) and y-axis represents selectivity (i.e., the degree to which its essentiality varies across lines). Legend defining dot colour is shown on the right. **C.** Human Protein Atlas RNA expression for *TSC22D, WNK,* and *NRBP* paralogs from tissue, blood, brain, single-cell sequencing (scSEQ), and cell line datasets. **D.** Immunoblot confirmation of protein levels in *WNK1*, *TSC22D2* and *NRBP1* mutant HAP1 query cell lines (n=2). **E.** Rank plot of qGI scores for representative *WNK2*(n=1), *WNK3*(n=1)*, TSC22D1*(n=1)*,* and *TSC22D2*(n=2) single mutant HAP1 query screens. Legend defining dot size and dot colour is shown on the right. **F.** Reproducibility of two biological replicates of *WNK1*, *TSC22D2, WNK2:WNK3*, and *TSC22D1:TSC22D2* mutant query screens using the within-between correlation (WBC) score described in^33^. **G.** Distribution plots and bar plots for the cell size data from **Fig. 1G**. WNK463, WNK-IN-11, rapamycin and Torin2 treatments are shown in greys and are performed for 24hrs with indicated dosages. DMSO treatment distributions are shown in red for comparison. **H.** Bar plots of normalized mean cell volume of RAW264.7, BV-2, THP-1, HeLa-Kyoto, U-937, and Huh-7 cells. Drug treatments were performed for 24hrs.

**Figure S2.** Kinetics of TSC22D2 foci formation, related to Figure 2. **A.** Immunoblots for mCherry polyclonal antibody to detect mScarlet that is endogenously tagged to the c-terminus of TSC22D2 in indicated cell lines. GAPDH served as the loading control. **B.** The left panel shows time-lapse images of HAP1 *TSC22D2^mScarlet^* cells following the indicated hypertonic stress including NaCl, KCl, and sorbitol. Small molecule crowding reagent urea is used for comparison. The right panel shows the quantification of average TSC22D2^mScarlet^ foci number per cell and the intensities of top 10% of TSC22D2^mScarlet^ foci under indicated conditions. **C.** Representative microscope images for TSC22D2^mScarlet^ foci formation under indicated concentrations of hypertonic stress. **D.** Representative microscope images for endo-tagged TSC22D2^mScarlet^ with known sub-cellular markers of distinct compartments including clathrin heavy chain (CLTC) for endosomes, LAMP1 for lysosomes, SQSTM1 for aggresomes, DCP1A for RNA granules, G3BP1 for stress granules and TOMM20 for mitochondria. The line graphs to the right represent intensity values as a function of the red line indicated in the merged images. The colors in the line graphs correspond to the proteins and are indicated in the legends. Pixel intensity show good localization with CLTC which is consistent with previous publications ^34^.

**Figure S3.** Proximity interactions of TWN proteins, related to Fig. 2 and 3. **A.** Predicted canonical WNK1 and NRBP1 structure. **B.** Strategies for tagging the N-terminus of WNK1 and NRBP1 with mNeon at their endogenous locus; the bottom panel shows. **C.** Representative time-lapse images before and after 30min hypertonic stress with various hypertonicity stress, including 230mM sorbitol, 167mM NaCl, and 167mM KCl, in ^mNeon^NRBP1 endo-tagged HAP1 parental cells and HAP1 cells with *TSC22D1:TSC22D2:TSC22D4* triple gene mutants. **D.** Immunoblotting for ^mT^TSC22D2 HAP1 cells to detect TSC22D2 and streptavidin. Tagged TSC22D2 shows an upshifted band compared with untagged wild-type (WT) while *TSC22D2* mutant line is deficient of TSC22D2. GAPDH served as the loading control. **E.** Co-immunofluorescence staining validates that the mT tags in *^mT^TSC22D2*, *^mT^WNK1*, and *^mT^NRBP1* cells do not interfere with formation of biomolecular condensates following a 1hr treatment with 460 mM sorbitol. **F.** Network analysis of the proximity interactions for each of TSC22D2, WNK1 and NRBP1 baits in the no stress (1Hr, B), sorbitol stress (1Hr, SB) and recovery (1Hr, SRB) conditions. The legend at the bottom defines the node and edge colors, thickness and size.

**Figure S4.** Validation of proximity associations, related to Fig. 3. **A**. Representative time-lapse images for co-tagging N-terminus of *TSC22D1*, *TSC22D4*, or *WNK3* with mNeon in *TSC22D2^mScaret^* HAP1 cells. The line graphs on the bottom represent intensity values as a function of the white box indicated in the merged images. The colors in the line graphs correspond to the proteins and are indicated in the legends. **B.** Representative immunofluorescence images staining preys of interest (i.e., WNK1 and CLINT1) with antibodies detecting endogenous proteins in TSC22D2^mScarlet^ HAP1 cells. The line graphs to the right represent intensity values as a function of the red line indicated in the merged images. The colors in the line graphs correspond to the proteins and are indicated in the legends. Representative immunofluorescence images of *^mNeon^WNK1:TSC22D2^mScarlet^* or *^mNeon^NRBP1:TSC22D2^mScarlet^* HAP1 cells following **C.** 30 min treatment with 0.5 mM sodium arsenate or **D.** 1 hr treatment with 460 mM sorbitol (bottom panel). TSC22D2 (yellow), WNK1 (green), G3BP1 (pink) and the merged images including nuclei (blue) are indicated. The line graphs to the right represent intensity values as a function of the red line indicated in the merged images. The colours in the line graphs correspond to the proteins and are indicated in the legends.

**Figure S5.** Kinetics of TWN bodies and stress granules (SGs), related to Fig. 3. **A.** Representative time-lapse images for ^mNeon^TSC22D2 and G3BP1^mScarlet^ dual tagged cells under hypertonic stresses including 460mM sorbitol, 230mM sorbitol, 167mM NaCl and 167mM KCl. **B.** Representative time-lapse images for ^mNeon^TSC22D2 and G3BP1^mScarlet^ dual tagged cells under 50mM sodium arsenate**.**

**Figure S6.** Evolutionary signature and TWN protein structure analysis, related to Fig. 4. **A.** Schematics outlining the core set of proximity associations that were examined for specific sequence features in panel B. **B.** The 144x208 z-score matrix of evolutionary signatures global heatmap with the core set against total human proteome. The z-score legends are selected against (blue) or for (yellow) amongst the indicated IDRs. **C.** Selected portions of heatmap presented in **Fig. S6B** highlighting key evolutionary features that are selected against (blue) or for (yellow) amongst the indicated disordered subsequences containing N terminal WNK1-3 and TSC22D1-2(cluster1, corresponding to disorder S1 in **Fig. S6D**), or WNK1-3 and NRBP (cluster2, corresponding to disorder S2 in **Fig. S6D**). The column labels are defined in **Data S2**. Z-scores were calculated as described in the Methods. **D.** TWN protein sequence features including protein domains summarized using InterPRO, disorder scrores predicted by IUPRED2A, regions of similar IDR features (disordered S1 and S2), and 3-dimensional structures predicted by Alpha-fold-2. **E.** PCA analysis on sequence percentile identity with input WNK kinase domains from 51 model organisms from **Fig.** **4D**.

**Figure S7.** WNK463 inhibition and ion imbalance trigger TWN body formation, related to Fig. 5. **A.** Images of additional time-lapse microscopy following the addition of 1 mM WNK463 to *^mNeon^NRBP1:TSC22D2^mScarlet^* HAP1 cells. **B.** Live cell imaging after 24hrs treatment of *TSC22D2^mScarlet^* HAP1 cells 5µM WNK-IN-11 at the indicated doses followed by treatment with 1,6-HD or 1,2,6-HT and imaging 2 minutes later. **C.** Experimental setup for condition-specific proximity mapping with 1µM WNK463 or 460mM sorbitol of TSC22D2, WNK1 and NRBP1 associations in HAP1 cells with endogenous N-terminal miniTurbo (mT) tags. **D.** Dot-plot graph showing recovery of TSC22D, WNK, and NRBP paralogs as preys, as well as OXSR1 and STK39 under indicated conditions. Dots are plotted as a function of the Bayesian False Discovery Rate or BFDR and spectral counts, as indicated in the legend. **E.** Representative time-lapsed microscope images for endogenously dual tagged G3BP1^mScarlet^ and ^mNeon^TSC22D2 cells following 1µM WNK463 (left) or 5µM WNK-IN-11 (right) treatment. WNK kinase inhibition induces TWN body formation but not SG. **F.** Images from time-lapse microscopy following the addition of hypotonic medium (IMDM:H_2_O=1:1) (left) or Isotonic (iso) sorbitol solution (300mOsmol/L) (right) to *^mNeon^NRBP1:TSC22D2^mScarlet^* HAP1 cells.

**Figure S8.** TWN body is induced by tonicity imbalance, related to Fig. 5 and 6. **A.** Representative time-lapse images of *^mNeon^NRBP1:TSC22D2^mScarlet^* cells treated with amiloride, a combination of bumetanide and hydrochlorothiazide (HCZT) and a combination of bumetanide, HCZT and amiloride with indicated dosages and time. **B.** Schematics for WNK463 chemical-genetics CRISPR screening. gRNA library (TKOv3) were transduced to HAP1 wild-type cells at MOI of 0.3. Transduced cell populations were then purified with puromycin selection and split at T0 into two arms. One arm is treated with 2.5µM WNK463 and the other arm is treated with vehicle control DMSO all the way to T18. T0 and T18 cell pellets are subjected to next-generation sequencing for determination of sensitizers and resisters of WNK463. **C**. Box-and-Whisker plot for change in [Na^+^]_intracellular_ comparing *TSC22D1:TSC22D2:TSC22D4*, *WNK1:WNK2:WNK3*, and *NRBP1:NRBP2* mutant clones and HAP1 parental cells.

**Figure S9.** TSC22Ds, WNKs and NRBPs are essential for proper TWN body formation, related to Fig. 6. **A.** Microscope images of ^mNeon^TSC22D2 or ^mNeon^WNK1 endo-tagged (green) HAP1 parental cells and *NRBP1:NRBP2* mutant cells. Blue signal represent DAPI(nucleus), and the orange signal represents co-stained SPAK or OSR1 kinases. **B.** Representative time-lapse images showing TSC22D2^mScarlet^ foci still form in WNK1 depletion cells after the addition of 1µM WNK463. **C.** Representative time-lapse images showing ^mNeon^NRBP1 and ^mNeon^TSC22D2 foci are compromised in *WNK1:WNK2:WNK3* triple depletion cells after the addition of 460mM sorbitol. **D.** Representative time-lapse images showing ^mNeon^NRBP1 and ^mNeon^TSC22D2 form compromised foci in *WNK1:WNK2:WNK3* mutant under 30min treatment of 230mM sorbitol, 167mM NaCl or 167mM KCl. ^mNeon^WNK1 forms foci under isotonic stress in *TSC22D1:TSC22D2:TSC22D4* mutant cells.

**SUPPLEMENTARY MOVIES & DATA TABLES**

**Movies S1-S3**. Kinetics of TSC22D2 foci formation and recovery using TIRF microscopy (frames 1-3). Related to Figure 2D.

**Data S1.** Cell size mini-screen for HAP1 queries. Related to Figure 1E.

**Data S2.** Protein sets identified for IDR feature analysis and z-score table using the core set of proximity associations for TSC22D2, WNK1 and NRBP1 BioID data to calculate correlation profiles based on evolutionary IDR features. Related to Figures 3G-H, 4A-C, and S6A-E.

**Data S3.** Drug mini-screen for effectors of TWN body formation. Related to Figure 5A.
